# Impact of tumor-parenchyma biomechanics on liver metastatic progression: a multi-model approach

**DOI:** 10.1101/2020.05.04.074989

**Authors:** Yafei Wang, Erik Brodin, Kenichiro Nishii, Hermann B. Frieboes, Shannon Mumenthaler, Jessica L. Sparks, Paul Macklin

**Affiliations:** Department of Intelligent Systems Engineering, Indiana University. Bloomington, IN USA; Department of Chemical, Paper and Biomedical Engineering, Miami University. Oxford, OH USA; Department of Bioengineering, University of Louisville. Louisville, KY USA; James Graham Brown Cancer Center, University of Louisville. Louisville, KY USA; Center for Predictive Medicine, University of Louisville. Louisville, KY USA; Lawrence J. Ellison Institute for Transformative Medicine, University of Southern California. Los Angeles, CA USA

## Abstract

Colorectal cancer (CRC) and other cancers often metastasize to the liver in later stages of the disease, contributing significantly to patient death. While the biomechanical properties of the liver parenchyma (normal liver tissue) are known to affect tumor cell behavior in primary and metastatic tumors, the role of these properties in driving or inhibiting metastatic inception remains poorly understood, as are the longer-term multicellular dynamics. This study adopts a multi-model approach to study the dynamics of tumor-parenchyma biomechanical interactions during metastatic seeding and growth. We employ a detailed poroviscoelastic (PVE) model of a liver lobule to study how micrometastases disrupt flow and pressure on short time scales. Results from short-time simulations in detailed single hepatic lobules motivate constitutive relations and biological hypotheses for a minimal agent-based model of metastatic growth in centimeter-scale tissue over months-long time scales. After a parameter space investigation, we find that the balance of basic tumor-parenchyma biomechanical interactions on shorter time scales (adhesion, repulsion, and elastic tissue deformation over minutes) and longer time scales (plastic tissue relaxation over hours) can explain a broad range of behaviors of micrometastases, without the need for complex molecular-scale signaling. These interactions may arrest the growth of micrometastases in a dormant state and prevent newly arriving cancer cells from establishing successful metastatic foci. Moreover, the simulations indicate ways in which dormant tumors could “reawaken” after changes in parenchymal tissue mechanical properties, as may arise during aging or following acute liver illness or injury. We conclude that the proposed modeling approach yields insight into the role of tumor-parenchyma biomechanics in promoting liver metastatic growth, and advances the longer term goal of identifying conditions to clinically arrest and reverse the course of late-stage cancer.

## Introduction

The liver performs critical physiological processes such as macronutrient metabolism, bile creation, and filtration of toxic substances from the blood [1]. Liver cells are organized in an intricate and complex architecture. The main structural units are the hepatic lobules, within which hepatocytes liver cells are arranged in a hexagonal pattern around the central vein. The portal triads (hepatic artery, portal vein, and bile duct) are found at the vertices of these hepatic lobules. See Fig 1. Fluid transport occurs along liver sinusoids, allowing for mixing of oxygen-rich blood from the hepatic artery with nutrient-rich blood from the portal vein while providing reaction surface for metabolic processes [2].

**Figure 1.**
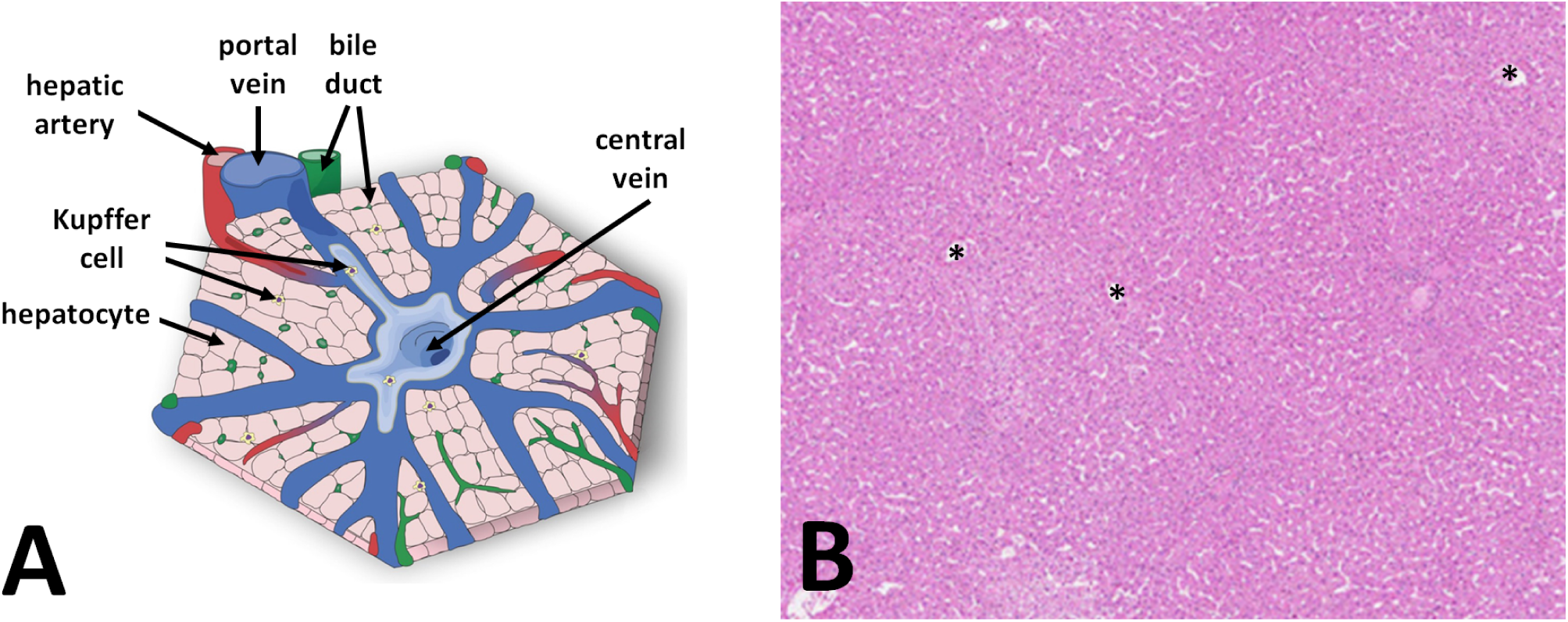
**A:** Schematic of a typical liver lobule. A poroviscoelastic (PVE) model will investigate flow and mechanics at short time scales at this spatial scale. Adapted under CC BY-NC 4.0 from [3]. **B:** Typical human liver histology (hematoxylin and eosin staining), with several central veins marked with **\***; the parenchyma is a mixture of sinusoids (white spaces) and hepatic cords (pink regions). An agent-based model will investigate tumor-parenchyma interactions over longer times in at this larger spatial scale. Adapted under CC BY-SA 3.0 from the Human Protein Atlas available from http://www.proteinatlas.org [4].

While the liver is critically involved in normal physiological functions, it is also a common site for metastatic disease. Liver metastases are a major barrier in cancer management. In particular, for colorectal cancer (CRC), patient mortality is principally due to the spread of disease to distant organs, with the liver being the predominant site in approximately 70% of the cases. This metastatic organotropism is mainly attributed to the drainage of the colon and rectum blood supply directly into the liver through the portal vein. Tumor cells that survive and proliferate in the liver microenvironment do so by replacing resident hepatocytes, co-opting the vasculature, adhering to hepatocyte-derived extracellular matrix (ECM), and communicating with neighboring stromal cells [5]. The reciprocal interactions between resident cells in the liver microenvironment govern the successful formation of metastatic tumor foci. Significant research has focused on the molecular, single-cell alterations that contribute to metastatic disease, yet the biophysical interactions between tumor cells and the metastatic microenvironment remain to be elucidated at multicellular scales over long times. This knowledge gap reflects both the emphasis of reductionist biology on molecular mechanisms of individual cancer cells and the difficulty in longitudinally observing tumor-parenchyma interaction dynamics in animal models or patients. Fresh insights on these interactions could improve therapeutic interventions and patient outcomes.

Tissue stiffness and other biomechanical factors are known to affect primary and metastatic tumor growth [6, 7]. In liver tissues, prior injuries and diseases such as cirrhosis increase fibrosis, and alter tissue stiffness [7]. Aging may also contribute to changes in liver tissue stiffness, as well as increase the likelihood of tissue fibrosis following an acute injury or illness [8–10]. The literature on the effect of such biomechanical alterations on tumor growth in the liver is seemingly inconsistent: some report low incidence of CRC metastases in cirrhotic livers (with accompanying fibrosis) [11], while others have reported that the risk of CRC liver metastases may be higher in patients with cirrhosis [12]. Still others report that stiff tissues can promote tumor growth and metastasis in the liver (e.g., [13]). Some of these conflicting reports may lie in the difference between *invasion* from a primary site at the start of the metastatic cascade (generally characterized as epithelial-mesenchymal transition, or EMT) and *seeding and growth* at a metastatic site (mesenchymal-epithelial transition, or MET) [14]. In EMT, tissue stiffness can up-regulate cell motility and matrix remodeling [15, 16], whereas stiff tissues at a metastatic site may impair tumor growth following MET [14]. Indeed, metastatic “seeds” arriving in a new “soil” may rest dormant for many years: a seeded metastasis may cease proliferating at a small, sub-clinical size or fail to grow at all; unknown later events may “reawaken” long-dormant metastases to resume rapid growth [16]. While there has been some prior study of the molecular-level triggers for metastatic reawakening (e.g., see discussion in [16]), far less is understood of the potential role of direct tumor-parenchyma mechanical interactions, or how mechanical changes following injury, illness, or aging could drive a dormant micrometastasis back to a rapidly growing state. Similarly, most investigations of age-driven changes in liver tissue have focused on immunologic and molecular-scale changes to ECM, rather than more basic mechanical properties (e.g., see [17]).

A more nuanced understanding of mechanobiological feedbacks between a seeded micrometastasis and the surrounding parenchyma could help detangle these inconsistent observations. To arrive at a more detailed understanding, interstitial fluid flow disruption and mechanical pressures imposed on the surrounding parenchyma by growing micrometastases needs further investigation, including how these can lead to parencyhma deformation and death, and how tumor-parenchyma mechanical interactions can alter the growth or arrest of micrometastases. With an improved understanding of these links, critical events most likely to drive emergence of new liver metastases in CRC patients may be identified, and thereby tailor more appropriate monitoring and early interventions for these at-risk patients. The limitations inherent in artificial *in vitro* environments and the confounding interactions present in *in vivo* models may be addressed via mathematical modeling and computational simulations, which allow systematic analysis by building, testing, and refining minimal models until they can fit prior observations. This work seeks to establish a modeling platform to investigate cell-stromal interactions and the associated tissue-scale effects within the context of the liver. In the longer term, such a platform could enable evaluating potential metastatic spread and progression based on patient-specific tumor cell and liver tissue characteristics.

### Prior mathematical modeling

While there have been numerous modeling advances in related areas, to the best of our knowledge, no model has been developed to address tumor-parenchyma biomechanical feedback in large tissues and the impact on tumor growth and parenchyma disruption. Please see *Appendix-Mathematical modeling of liver biology and metastases, and cancer mechanobiology* in the *Supplementary Materials* for an expanded overview of the prior work. Briefly, there is a rich history of mathematical modeling of liver biology, particularly liver tissue regeneration [18, 19], liver fibrosis [20, 21], liver toxicology [22, 23], interstitial flow and mechanics in liver tissues [24–30]. Others have modeled the role of tissue mechanics on (mostly primary) tumor growth [31–34], including some excellent work on the role of mechanosensing in individual tumor cell proliferation [35, 36], although these did not focus specifically on primary or metastatic tumor growth in liver tissues. While there is less history in specific modeling of liver metastases, some have modeled the growth of metastases in the liver [37–39], while others have simulated the impact of ECM remodeling and macrophages in liver metastases [40–42]. There are also good examples of simulating nanotherapy treatments [43–45] and theranostics [46, 47] of liver metastases. Others have developed network models of metastatic seeding [48–53], but without spatiotemporal detail and without considering the impact of tumor-host biomechanical interactions.

In spite of previous empirical and mathematical modeling efforts, the impact of parenchymal mechanics and flow on metastatic growth in the liver remains poorly understood, as is the biology regulating tumor-parenchyma interactions near the tumor boundary. And while emerging biomimetic platforms (e.g., [54, 55]) can evaluate some of these dynamics, it remains difficult to resolve tumor foci at multicellular resolution over long times. Past multiscale modeling investigations have mainly focused on discrete or smaller-scale simulations, or worked to provide insights at larger, continuum scales. However, none have investigated cell-level mechanobiologic effects over large tissues, nor have they been motivated by detailed continuum models of interstitial flow to drive efficient whole-tissue scale models over long times.

### Study approach and summary of novel findings

The modeling framework presented here aims to probe these knowledge gaps. Because many prior parsimonious models have matched observations with simple substrate-driven growth feedbacks, we first (1) evaluate whether disrupted interstitial flow significantly slows tumor growth by limiting the availability of cell oxygen and nutrients, and (2) determine if mechanical pressure feedbacks on tumor cell proliferation are necessary to avoid non-physical model behaviors. Our novel combination of a detailed lobule-scale flow model, mathematical analysis, and an agent-based multicelular model allows us to determine that for metastatic growth into stiff, well-perfused tissues, any minimal model of tumor-parenchyma interactions must combine substrate- and biomechanics-driven birth and death.

We perform a parameter space investigation of the minimal tissue-scale model to (3) identify the conditions under which parenchymal biomechanics results in tumor dormancy, and (4) evaluate whether changes in parenchymal biomechanics (e.g., from aging, injuries, or acute disease) can reawaken a dormant tumor. We find that micrometastases can arrest at subclincical sizes when elastic forces (which stall tumor cell cycle entry) are not relieved by death or plastic relaxation in the parencyhma. We also observe sustained metastatic growth when the parenchyma cannot tolerate significant elastic deformation. These model-driving findings given dynamical context to static clinical observations. Moreover, the model predicts that changes in parenchyma tissue mechanics can push a dormant micrometastasis into sustained growth.

Interestingly, the simple mathematical model predicts these phenomena without the need for complex cell signaling, ECM remodeling, or other molecular-scale phenomena. This work showcases how simple mathematical models can be computationally scaled to larger tissues to assess complex cancer biology. Future incorporation of detailed molecular scale models could help further refine how the cell and tissue agents achieve the basic behaviors in this minimal model, while the emergent multicellular behaviors help explain how molecular- and cell-scale behaviors drive long-time behavior of metastases.

## Results

### Modeling approach

This study combines two modeling approaches to investigate metastatic tumor seeding and growth in liver tissues. A high-resolution poroviscoelastic (PVE) model of interstitial flow and tissue mechanics gives detailed insights on a tumor’s impact on interstitial flow, transport, mechanical pressure, and tissue deformation in and around a tumor micrometastasis for a quasi-steady tumor on short time scales. We use an efficient, minimal lattice-free agent-based model (ABM) framework to simulate individual CRC cells and parenchyma agents in centimeter-scale liver tissues over weeks and months.

Unlike typical matched models where small-scale discrete models motivate large-scale continuum models, this study uses a detailed continuum model of flow mechanics to generate hypotheses and simplifications for scalable discrete models. We use the PVE model results to build constitutive relations for growth substrate transport at centimeter scale in tissues with hundreds of lobules. We also use the PVE results to motivate a minimal set of biological rules (hypotheses) to study tumor cells and parenchymal tissue in a centimeter-scale agent-based model (ABM) of tumor metastatic growth in liver tissue. The multi-model approach is summarized in Fig. 2.

**Figure 2.**
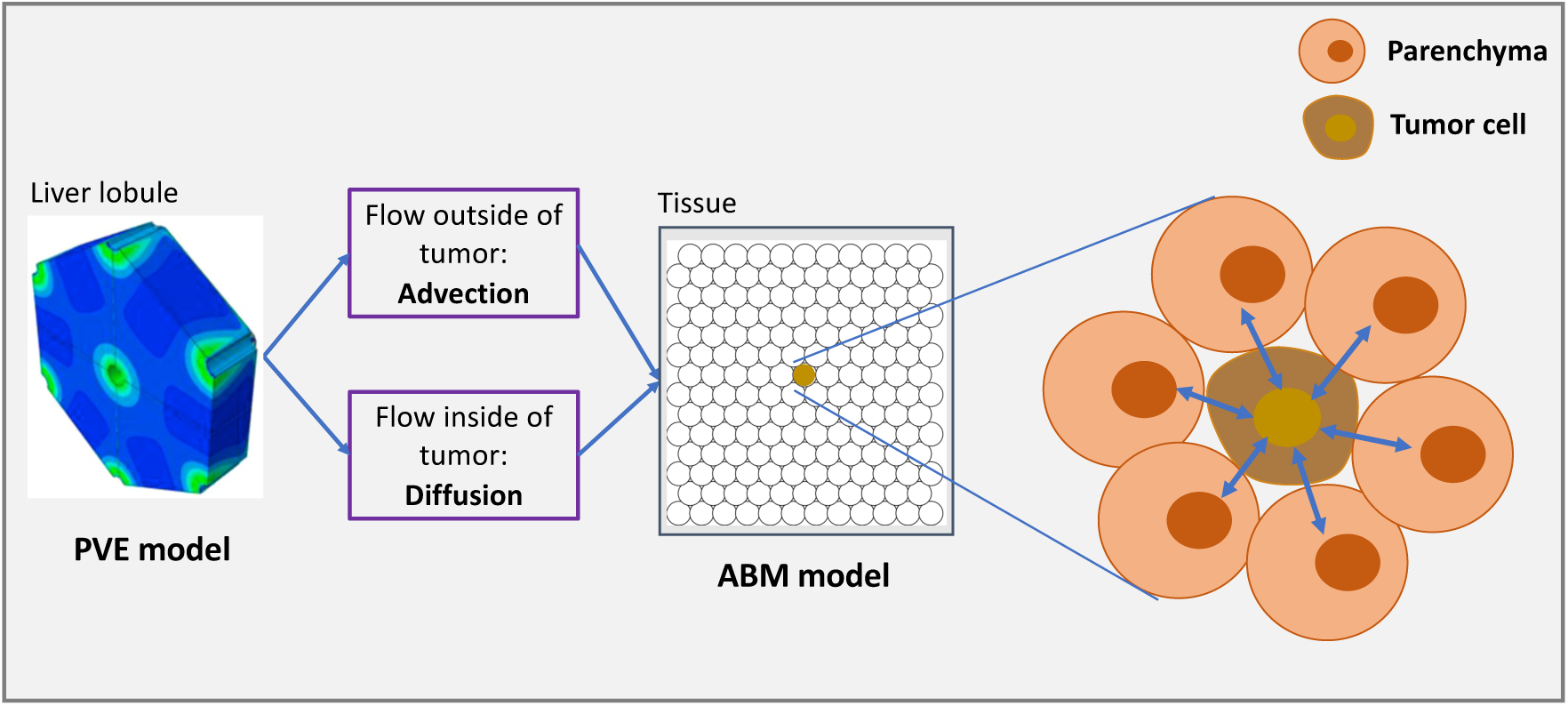
Mathematical modeling approach. Short-time simulation studies of the fine-scale poroviscoelastic (PVE) model give insights on perfusion within individual liver lobules. These results drive the development of an agent-based model (ABM) to investigate the dynamics of tumor-parenchyma interactions in larger tissues over longer times.

### Small micrometastases have little impact on flow in nearby parenchyma, but they increase strain

In the absence of a tumor (control case), our PVE model simulations show that the flow velocity increases in an approximately exponential pattern from the portal vein inlet to the central vein outlet, with velocity magnitudes ranging from 0 to ∼800 *µ*m/sec. The solid matrix strain decreases from approximately 3% at the lobule periphery to 1.7% near the central vein outlet. See *Supplementary Fig S1*.

We next seeded very small micrometastases (50 *µ*m diameter) in the lobule, with no net fluid pressure drop across the tumor boundary. The simulated flow velocity and tissue strain away from the tumor are largely unaffected by the presence of a nearby metastatic seed. See the “no tumor” paths in *Supplementary Fig S2*. Along the “tumor” path, the fluid velocities are notably elevated in the high permeability tumor condition (200 *µ*m/sec) as compared with the low permeability tumor condition (57 *µ*m/sec) or the normal parenchyma at a comparable location within the lobule (95 *µ*m/sec at 200 *µ*m from the center of the lobule). In both these scenarios, we note that interstitial flow is largely intact near the micrometastasis. (See the top left plot in *Supplementary Fig S2* for the high-permeability flow profiles, and the lower left plot for the flow profiles along the “tumor” and “non-tumor-paths” for both the high- and low-permeability cases.) The strain increases at the tumor margins, from 3% to ∼4.2%, and it drops to negligible values inside the tumor itself as a result of the increased tumor stiffness relative to the surrounding tissue (refer to Supplementary Materials for PVE Model for details regarding tumor stiffness used in the model). See the right plots in *Supplementary Fig S2*.

### Flow is stagnant within smaller micrometastases but not in the surrounding parenchyma

We next simulated the disruption of the liver parenchyma surrounding a 200 *µ*m micrometastasis between a portal vein and the central vein, with the tumor modeled as a pressure sink; see Fig. 3. For the flow path passing through the tumor, the fluid velocity entering the lobule from the portal vein is higher (800 *µ*m/sec) compared with control (∼0 *µ*m/sec), increasing to a maximum of 1370 *µ*m/sec as it approached the tumor boundary. Thus, the flow is non-stagnant in the regions surrounding the micrometastasis. Flow along the path through normal parenchyme has comparable velocities compared to the control near the portal triad and middle of the lobule, but lower velocities towards the central vein outlet. Notably, the fluid velocity drops to negligible values within the tumor itself.

**Figure 3.**
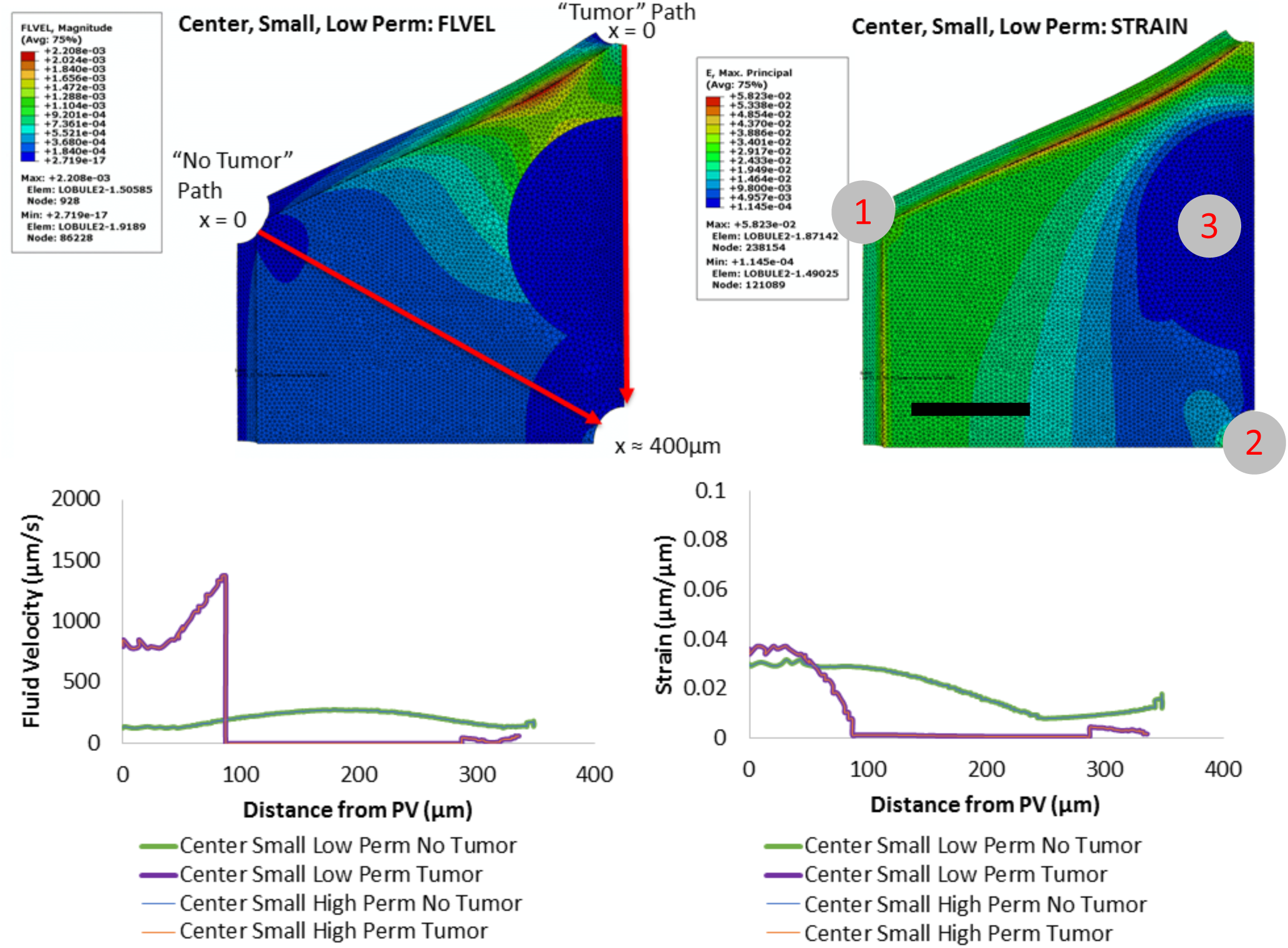
Hepatic flow: small micrometastasis. A 200-*µ*m micrometastasis disrupts a hepatic lobule, with tumor treated as a net pressure sink. Here, the fluid flow velocity increases “upstream” of the tumor (near portal triad inflow) and decreases near the central vein outlet. The fluid velocity abruptly drops to zero in micrometastases with both high- and low-permeability (left plots). It is noted that the magnitude powers of flow of “No Tumor” and “Tumor” paths are 10^−4^ and 10^−17^ respectively in left-top sub-figure. In the right-top sub-figure, 1 is the position of portal triad, 2 is the position of the central vein, 3 is the tumor position. <u>Scale bar</u>: 100 *µ*m.

This trend holds for both high and low permeability tumor conditions. Moreover, similar simulations that varied the position and tumor permeability for 200 *µ*m micrometastases similarly show stagnant flow within the tumors and non-stagnant (or increased) flow in the regions immediately surrounding the tumors, likely due to the tumor set as a pressure sink (see table in boundary conditions section of *Supplementary Materials for PVE Model*). This is true regardless of the tumor location (center or portal) or permeability (high or low).

This trend continues for larger micrometastases, but with clearer effect. When we placed a 400 *µ*m micrometastasis near a portal vein and treated the tumor as a pressure source, flow is stagnant within the tumor and intact (or accelerated) in the surrounding parenchyma. See *Supplementary Fig S3*. Fluid velocities increase sharply from zero to ∼1000 *µ*m/sec at the tumor boundary, then continue to increase up to ∼2000 *µ*m/sec at the central vein. Flow along the path through normal parenchyme shows a velocity trend similar to the control condition but with a higher peak velocity (∼2000 *µ*m/sec) at the central vein. This result holds for high and low permeability when the tumor is treated as a pressure source *(Supplementary Fig S3 and Fig S11)*, or as a net sink *(Supplementary Fig S14)*. It also holds when the 400 *µ*m micrometastasis is simulated as pressure neutral with low permeability *(Supplementary Fig S12)*.

### Larger micrometastases can exert tissue strains on the surrounding parenchyma

We observed significant tissue strains in the liver parenchyma surrounding the 400 *µ*m micrometastases *(Supplementary Fig S3*. For the smaller 200 *µ*m micrometastasis, the strain decreases from 3% to zero at the tumor boundary and remains negligible within the tumor itself. For the larger 400 *µ*m micrometastasis, the strains increase sharply at the tumor boundary to a peak value of 9%, the highest strain magnitude of any condition. We note that setting the tumor as a pressure sink produces a drop in strain at the tumor edge (as in the 200 *µ*m micrometastasis in Fig. 3), unlike the spike in strain noted in the other two conditions (*Supplementary Fig S2 and S3)*.

### Biotransport is mainly diffusive within micrometastases, and largely advective and quasi-steady in unobstructed parenchyma

In the PVE simulations, we found that flow is stagnant within most micrometastases, and non-stagnant (e.g., comparable to control or accelerated) within non-obstructed regions of parenchyma. After dimensionless analysis of a general conservation law in these two respective regions, we found that substrate transport is largely advection-driven in intact parenchyma, and primarily diffusion-dominated in tumor regions. We built on these results to obtain an approximate analytical solution to the quasi-steady distribution of cell substrates within an unobstructed lobule, as well as to approximate advection-dominated perfusion in large tissue sections. See the *Supplementary Materials* for further detail on the dimensional analysis and approximate solutions.

### Additional growth feedbacks are needed to model well-perfused liver tumors

Most current tumor growth simulation models, such as [56–63] use a diffusing growth factor (e.g., oxygen or glucose) as the main negative feedback on tumor cell proliferation by scaling cell cycling with local substrate availability. As the avascular tumor grows, it consumes and depletes the substrate, thus slowing growth in a negative feedback. Other models have focused on mechanical growth feedbacks, such as work by Drasdo and co-workers [64] or cellular automaton models that only allow proliferation when a neighboring lattice site is unoccupied [65, 66]. See the excellent review in [67]. While simulation frameworks such as PhysiCell are capable of simultaneously modeling substrate and mechanical growth feedbacks, most models consider one or the other. This in part reflects the goal of model *parsimony*, where simple models are selected over more complex models (with more degrees of freedom) when they can sufficiently fit available data and observations.

Because the PVE simulations and subsequent analyses showed that liver parenchyma is well-perfused in tumor-free regions, we questioned whether substrate-driven growth limits alone (as a parsimonious model) can prevent non-physical behaviors (e.g., extensive cell overlap) or if mechanobiologic feedbacks are also required as in [64]. To pursue this question further, we combined the analytical approximation for advection-dominated non-tumor regions from the Section “Approximate oxygen distribution in large tissue sections” in Supplementary Materials with the BioFVM diffusion solver in PhysiCell. With the methodology as outlined in Methods, we seeded a 1-cm^2^ liver tissue with 10 tumors with a distribution of sizes (diameters) from 250 *µ*m to 2 mm. We then held each tumor static (mechanics, cycling and necrosis disabled) and simulated to steady state. See Fig. 4a. From the heat map of oxygen diffusion (Fig. 4b), oxygen concentration outside of tumors varies from ∼38 mmHg to ∼60 mmHg, while pO_2_ inside of tumors depends upon the tumor size. For many of these tumors, most of the tumor cells are above the hypoxic threshold (5 mmHg; see [61, 68] for more discussion of default parameter values) below which proliferation is disallowed and cells may necrose. The statistics of oxygen distribution in different tumors are shown in (Fig. 4c), with big tumors having lower means and larger standard deviations as a result of the larger oxygen gradients in those larger tumors. In Fig. 4d, we overlay for each tumor its oxygen versus distance from the tumor edge. The 800 *µ*m tumor completely lacks a necrotic region (pO_2_ *<* 5 mmHg), the 1 mm tumor only has a small region below the threshold, and larger tumors have at least 400-500 *µ*m of viable tissue before reaching the necrotic threshold.

**Figure 4.**
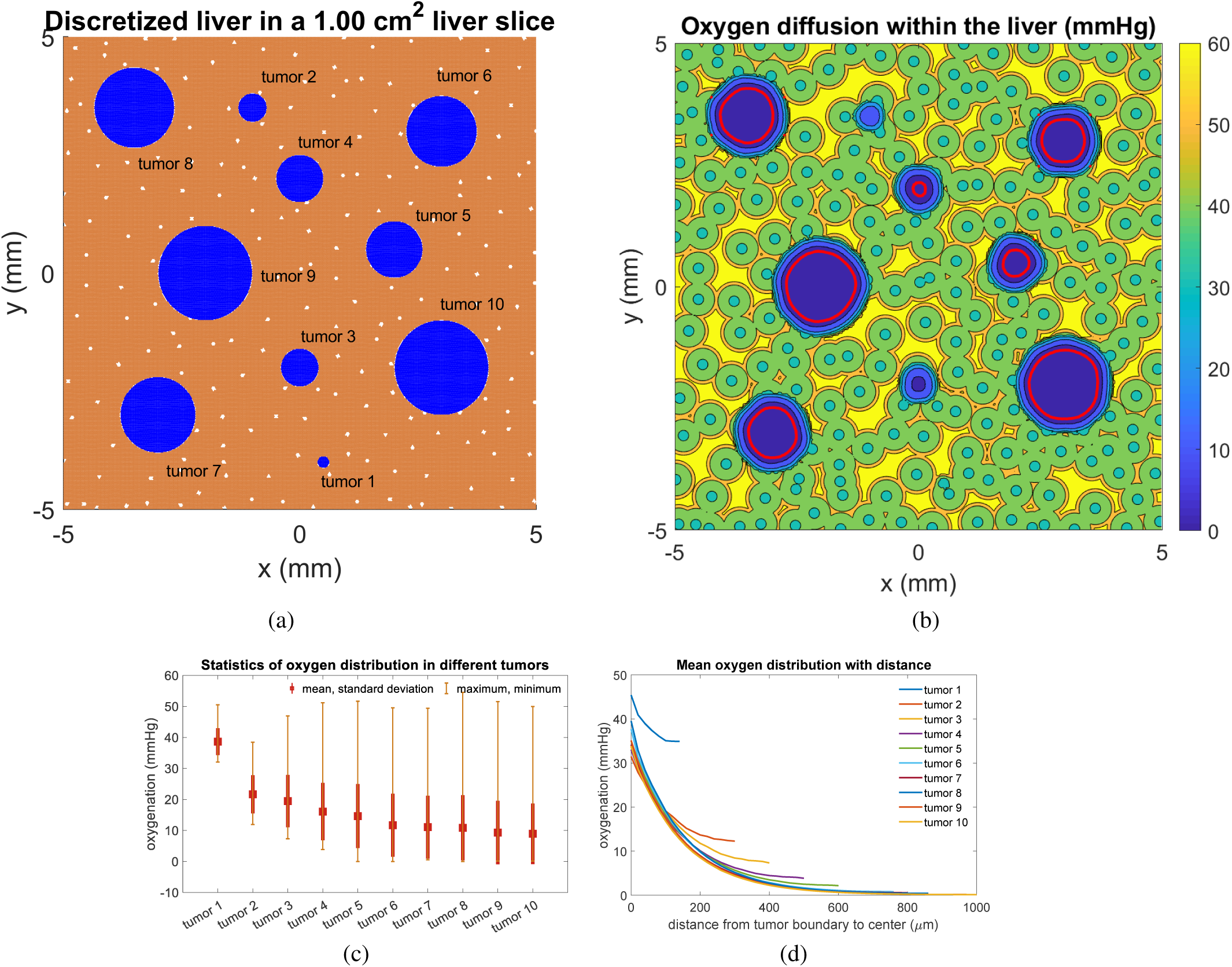
Oygen distribution inside of 10 tumors (red line is the necrosis threshold value (5 mmHg)) within liver tissue. **(a)**: the 10 tumors in liver tissue; **(b)**: corresponding oxygen map; **(c)**: statistics of oxygen distribution in the different tumors; **(d)**: corresponding mean oxygen distribution with distance

We tested growth of a micrometastasis into liver tissue with oxygen-based growth control (as in [61]) but without a mechanobiologic feedback. We found that pressure peaked at the tumor center and decreased towards the tumor-parenchyma interface, which agrees well with Stylianopoulos et al.’s experimental and mathematical results [69]. See *Supplementary Fig S4* for an example using the more advanced liver mechanics to be introduced below. While liver parenchyma apoptosis near the tumor boundary helped to locally relieve this pressure, we observed an excessive non-physical overlap of cells and (nondimensional) pressures well over 0.5 (the pressure value for a confluent, uncompressed tissue). Therefore, we surmised that an additional mechanobiologic growth feedback is required to model metastatic tumor cells in the well-perfused liver environment.

### Model behavior with mechanobiologic feedbacks

In the PVE model results, we saw evidence of significant tumor-parenchyma mechanical coupling, even for static tumors. Moreover, attempts at parsimonious models that rely solely upon diffusion-driven growth limits are insufficient to avoid non-physical model behaviors for metastases growing in well-perfused liver tissues.

To address this, we tested whether the addition of mechanobiologic feedbacks could prevent such non-physical behaviors while opening space for a growing micrometastasis. As in work by Drasdo and Byrne [70], we decreased tumor cell proliferation with increasing mechanical pressure. With this modification, rapid tumor proliferation into the elastic parenchyma leads to increasing pressure in the tumor that reduces proliferation and avoids the non-physical pileup of tumor cells. (We note that substrate-dependent proliferation is still needed to capture the effect of greatest tumor cell proliferation at the outermost edge, while substrate-dependent cell survival is necssary to capture the emergence of a necrotic core.) We also modified the parenchyma agents to scale death with increasing elastic displacement. In cases of rapid metastatic growth (relative to plastic relaxation), parenchyma death can reduce the mechanical pressure. See additional details in the Method section.

With the addition of these mechanobiologic feedbacks, we posited that the long-term fate of a metastatic seed (e.g., failed growth, arrest at a steady size, or growing without bound) would depend upon the balance between (substrate- and pressure-dependent) tumor proliferation, substrate-driven tumor necrosis, elastic resistance forces generated by displaced parenchyma, mechanics-driven parenhcyma death, and plastic reorganization of the parenchyma. We performed a parameter space investigation to determine how these effects interacted to drive metastatic growth.

We initiated a series of simulations of 90 days of growth from a single seeded metastatic cell in the center of a 0.5 cm × 0.5 cm liver tissue. We selected 27 parameter sets (combinations of 3 parameters at low, medium, and high values; see Table 1) based on previous experimental [71] and simulation [33] results. In particular, the base parameter set uses *r*_E_ = 0.1 min^*-*1^ so that elastic deformation occurs on the time scale of minutes, *r*_P_ = 0.001 min^*-*1^ so that plastic relaxation is two orders of magnitude slower than the elastic response, and *d*_max_ = 1.5 *µ*m is on the order of 10% of the size of a parenchyma agent (15 *µ*m radius). We increased and decreased each of these parameters by a factor of 2, yielding 27 parameters sets, and ran 10 simulations per parameter set due to PhysiCell’s stochasticity (a total of 270 simulations).

**Table 1.** Parameters used for the agent-based model biomechanics investigation.

|  |  |  |  |
| --- | --- | --- | --- |
| $r_E \text{ (min}^{-1}\text{)}$ | 0.05 | 0.1 | 0.2 |
| $r_P \text{ (min}^{-1}\text{)}$ | 0.0005 | 0.001 | 0.002 |
| $d_{\max} \text{ (}\mu\text{m}\text{)}$ | 0.75 | 1.5 | 3 |

In Fig. 5, we plot the final images (at t = 90 days) for a representative simulation for each parameter set. In these and all subsequent figures, blue cells are Ki67-tumor cells (quiescent), green cells are Ki67+ cells preparing for division, magenta cells are Ki67+ cells following mitosis, red cells are apoptotic, and brown cells are necrotic. Tumor cells with pressure *p* exceeding the maximum pressure *p*_2_ = 1 are colored yellow; these cells will arrest cell cycling in the Ki67-state. Liver parenchyma agents are shaded from brown to bright magenta based upon their displacement (strain) relative to their maximum displacement *d*_max_ (brightest magenta cells have the greatest displacement). Parenchyma agents with displacement exceeding 0.01 *d*_max_ were given a black outline to further indicate regions of tissue deformation. Apoptotic parenchyma are colored black.

**Figure 5.**
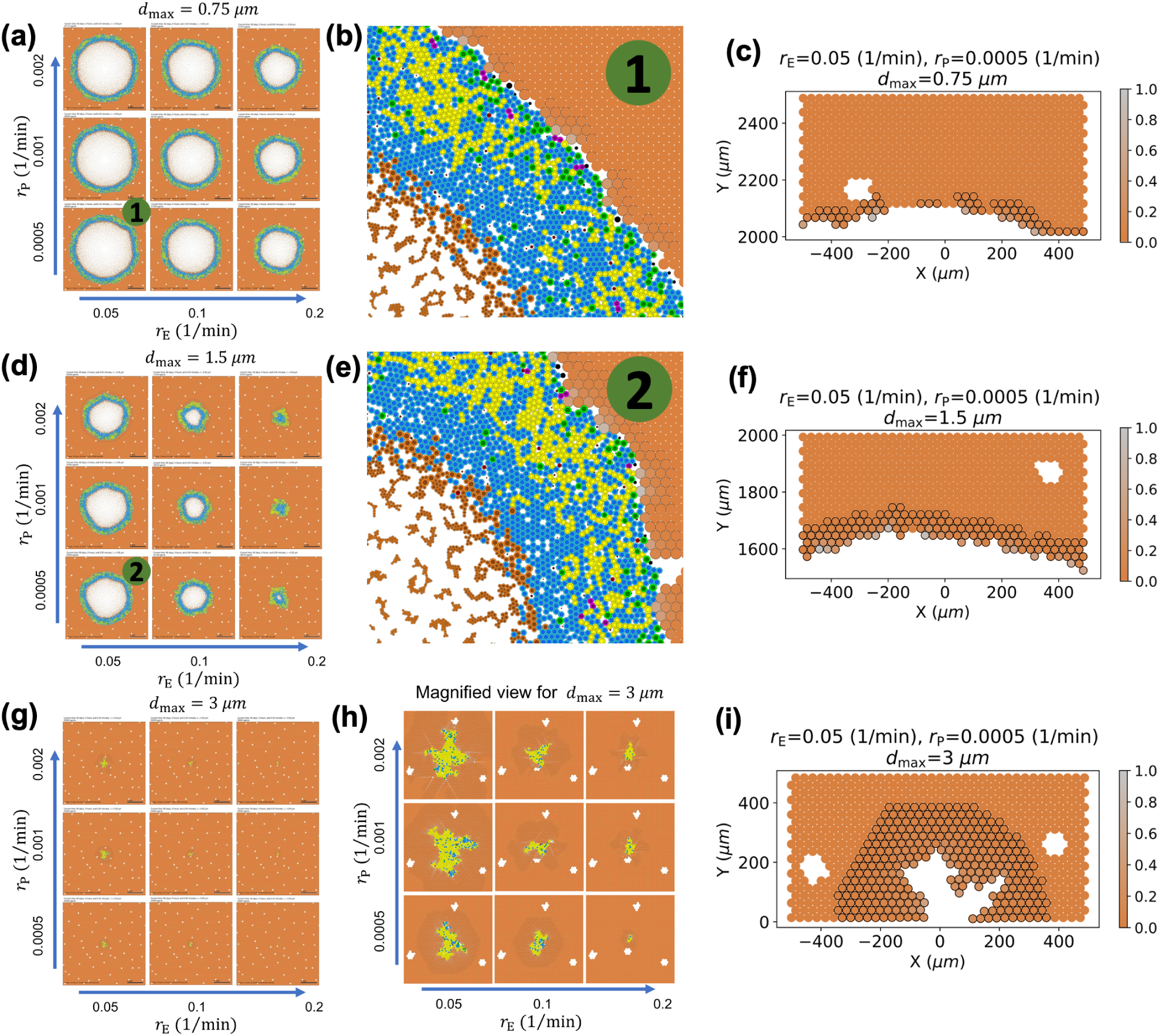
**(a, d, g):** Final tumor shapes at t = 90 days in 0.5 cm × 0.5 cm liver tissue with different biomechanical parameters (*r*_E_: elastic response rate; *r*_P_: plastic relaxation rate; *d*_max_: maximum tolerated deformation). **(h):** 5 magnification of the tumors in **(g). (b, e):** 6 × magnification of the tumor boundary section in ➀ and ➁ of corresponding parameteres in **(a, d). (c, f, i):** Crop section of viable parenchyma in tumor boundaries of three different maximum tolerated displacement. Parenchyma agents that have at least 1% of their maximum tolerated displacement are outlined with black edge color, and agents are colored by their relative strain (from bright orange to light gray). **<u>Legend</u>:** Tumor cells are shaded blue (Ki67-), green (Ki67+ prior to division), magenta (Ki67+ after division), red (apoptotic), or brown (necrotic). Yellow tumor cells have pressure *p* exceeding their tolerance and will arrest in the Ki67-state. **Liver parenchyma** is shaded from bright orange (no deformation) to light gray (maximum tolerated deformation), and black parenchyma is apoptotic. Black outlined parenchyma agents have been displaced at least 1% of their maximum tolerated displacement. Original high-resolution SVG files are included in github repository: https://github.com/MathCancer/liver-mechanobiology

The final tumor size decreases with increasing *d*_max_ (maximum tolerated deformation). For *d*_max_ = 3 *µ*m (Fig. 5g-h), the majority of tumor cells are yellow, indicating high pressures and prevalent cycle arrest. Tissue strain extends far into the surrounding parenchyma. (See bands of parenchyma with outlines in Fig. 5h-i) In these conditions, the final tumor are small with irregular morphologies. For lower values of *d*_max_, the tumor-parenchyma boundary is much smoother. Tumors exceeding approximately 1 mm in diameter showed necrotic regions.

### Tumor-parenchyma biomechanical interactions generate pressure inside tumors and tissue strains in the nearby parenchyma

For *d*_max_ ≤ 1.5 *µ*m, the greatest concentration of tumor cells exceeding their arrest pressure occur in the outer portion of the viable rim where tumor cell proliferation has the highest probability. As daughter cells are generated and grow, the tumor tissue encounters (elastic) mechanical resistance by the parenchyma, leading to the local pressure increase. (See the yellow cells in Fig. 5a-b.) In necrotic regions, shrinking necrotic cells locally reduce the pressure; this is reflected in the reduced density of yellow cells near the perinecrotic boundaries. See a cross-sectional plot of the tumor pressure and parenchyma strain (for *r*_E_ = 0.05 min^*-*1^, *r*_P_ = 0.0005 min^*-*1^, and *d*_max_ = 0.75 *µ*m–a parameter set that permits sustained tumor growth) in Fig. 6.

**Figure 6.**
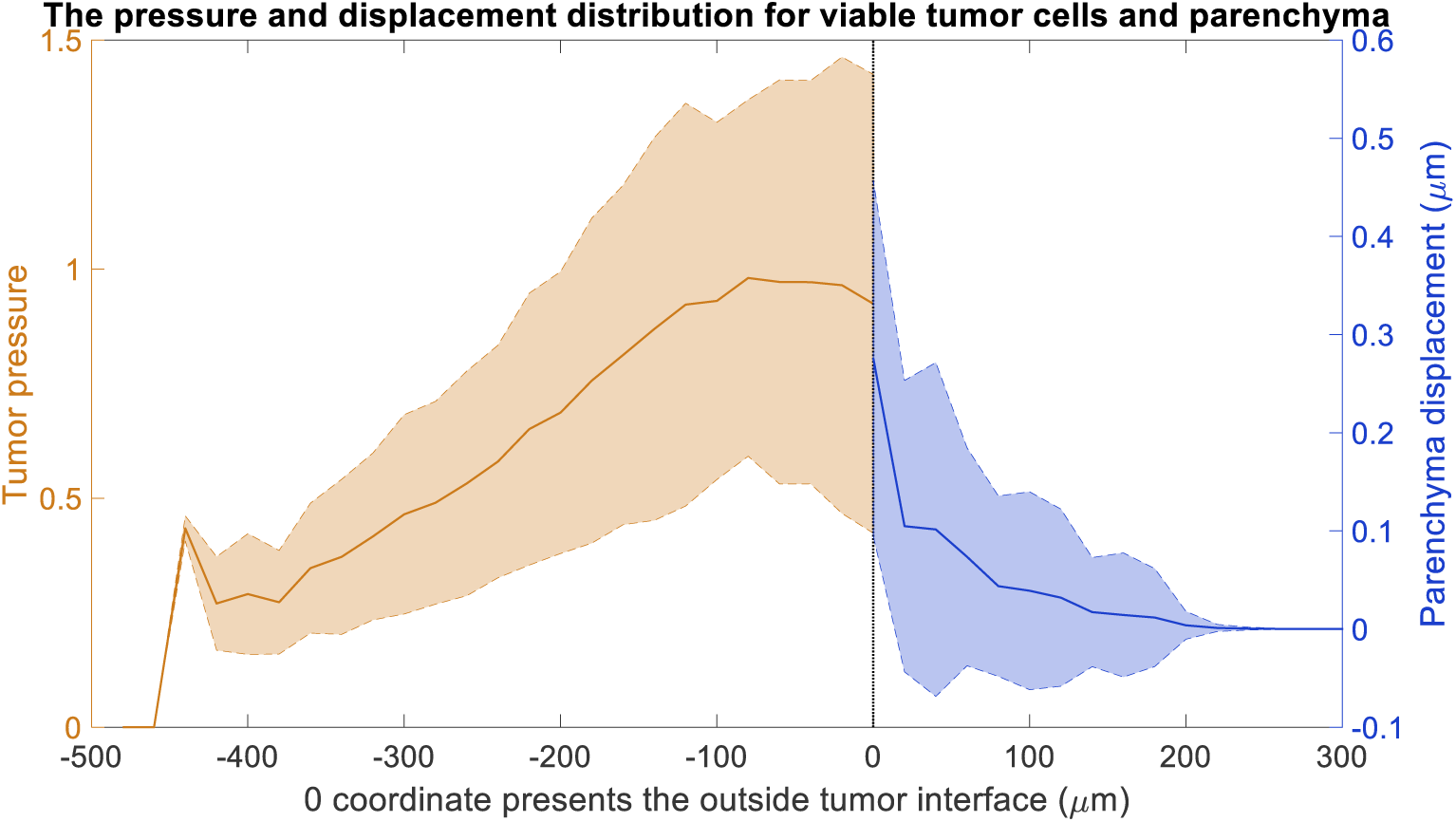
Pressure distribution of viable tumor cells and the displacement distribution of parenchyma at t = 90 days, where *r*_E_=0.05 (1/min), *r*_P_=0.0005 (1/min) and *d*_max_=0.75 *µ*m. Cells are colored as described in Fig. 5.

The proliferative tumor regions expand the volume of the tumor viable rim, thus deforming the nearby parenchyma. The parenchyma strain (brightest magenta: greatest relative deformation) reaches a maximum at the tumor-parenchyma interface and steadily dissipates with distance from the tumor, similarly to the poroviscoelastic simulation results. Apoptosis (black agents) occurs nearest to the tumor-parenchyma interface wherever the parenchyma strain exceeds its maximum tolerance. See Fig. 6.

Fig. 7 plots key properties of tumor and parenchyma versus time (days) with the same parameters as Fig. 6, visualized for all 10 simulation replicates. For the first 20 days, the fraction of the tumor exceeding the maximum pressure steadily grows, reflecting both the proliferation and the mechanical resistance of the surrounding parenchyma. Afterwards, the pressure reduces because (1) the pressure feedback reduces proliferation, and (2) necrosis acts as a pressure release. Over time, these effects balance to reach a steady state. See Fig. 7(a-b). The mean tumor pressure behaved similarly (result not shown). The mean parenchyma displacement increases steadily over time(Fig. 7(c), which reflects the increasing tumor-stroma contact area as the tumor expands into the parenchyma.

**Figure 7.**
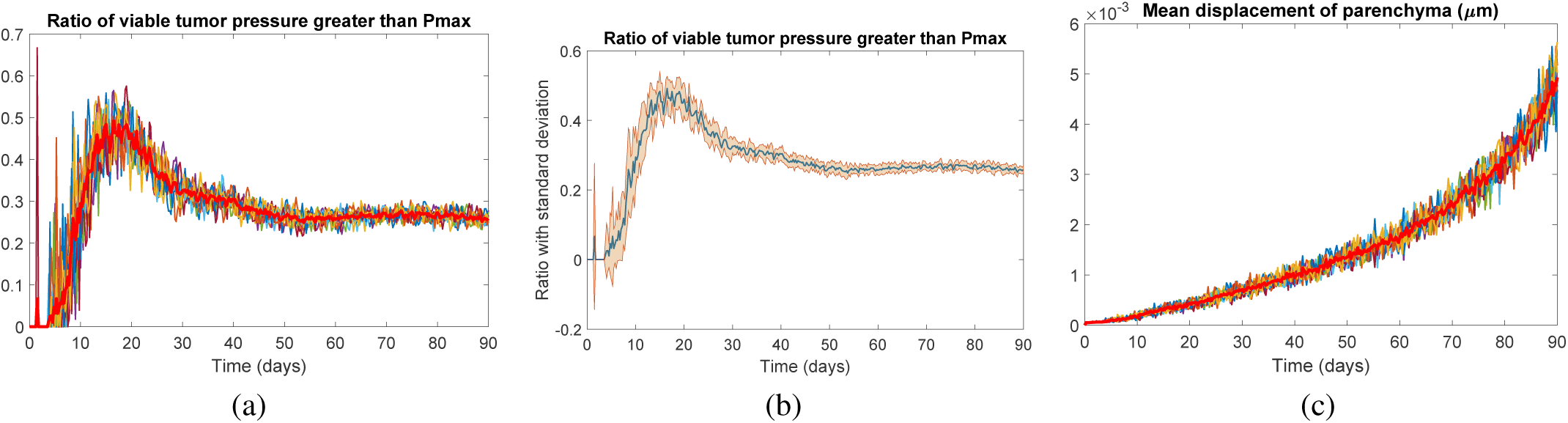
Longitudinal changes in pressure over 90 days. In **(a-b)** we plot the fraction of tumor cells exceeding the arrest threshold, showing 10 individual profiles **(a)** and the average and standard deviation of 10 runs **(b). (c)** plots the mean parenchyma displacement. Here, *r*_E_=0.05 (1/min), *r*_P_=0.0005 (1/min) and *d*_max_=0.75 *µ*m.

### Predicted parenchyma deformations are observed in human colorectal cancer metastases

The model predictions of deformed liver parenchyma can be observed in clinical samples. In Fig. 8, we show a hematoxylin and eosin (H&E) stained whole slide image of human liver tissue with a colorectal cancer metastasis. Regions of compressively strained liver parenchyma (C) are macroscopically observable near the tumor boundary (dashed line), visible as regions of more concentrated eosin staining. See Fig. 8A.

**Figure 8.**
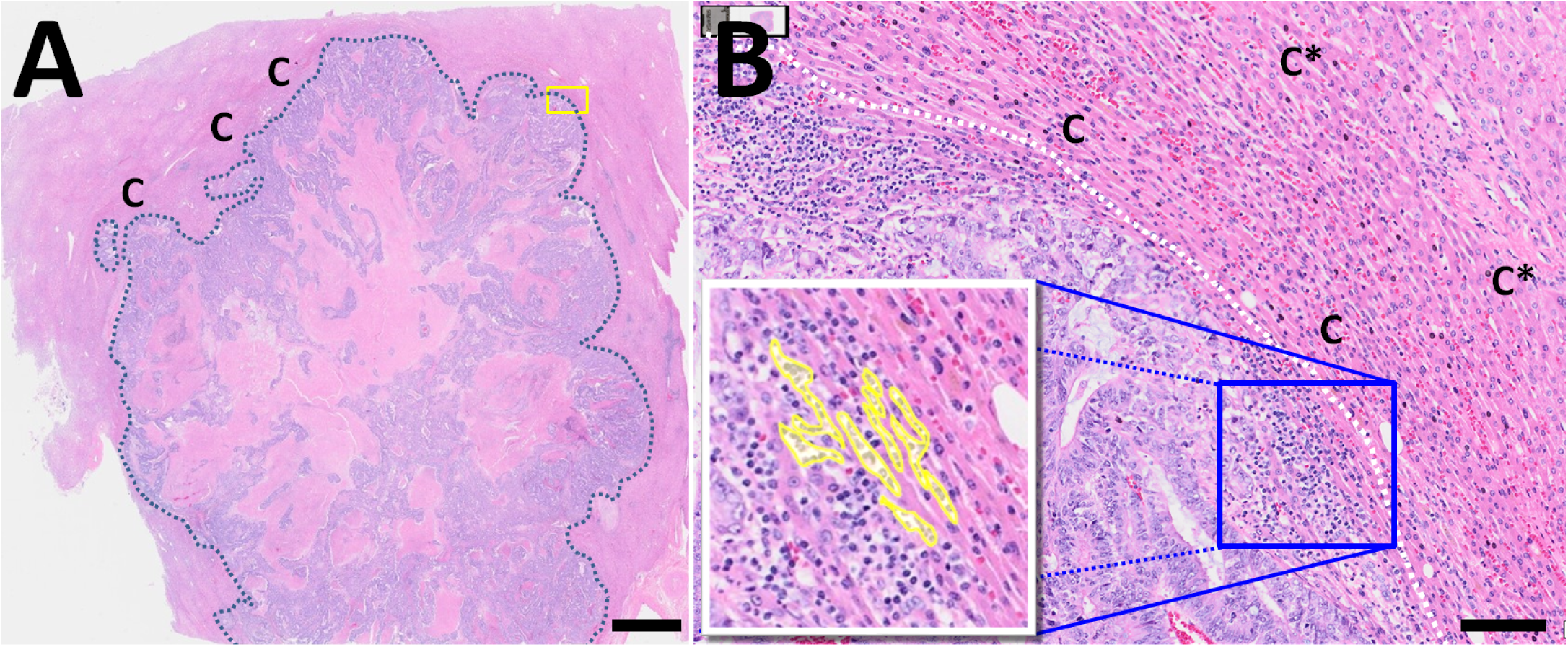
Colorectal cancer metastasis in human liver: **A:** Hematoxylin and eosin (H&E) stained whole slide image of a CRC metastasis in liver. Note the deformed liver parenchyma (C) near the tumor-parenchyma boundary (dashed line), which indicates sufficient tissue deformation (strain) to be observed macroscopically. <u>Scale bar</u>: 2 mm. **B:** Magnified view of the yellow box from A near the tumor boundary. Note that the regular liver microstructure has been deformed (strained) to be highly anisotropic, indicating compression in the direction orthogonal to the tumor boundary (dashed line) and potential stretching in the direction parallel to the boundary. This deformation is greatest near the tumor (C) and decreases farther from the tumor boundary (C*). The yellow highlighted cells in the inset show tumor cells growing along paths of mechanical least resistance in collapsed sinusoids. <u>Scale bar</u>: 100 *µ*m.

In a magnified view of the sample (Fig. 8B), we see that the parencyhma nearest to the tumor is highly anisotropic: it has been compressed in the direction parallel to the tumor boundary (in the direction of tumor growth), and stretched in the direction parallel to the boundary. This is consistent with deformation from a growing tumor: compression in the direction of growth, and stretch in the direction parallel to the interface. Moreover, just as in the simulations, the greatest parenchyma deformation happens nearest to the tumor (annotated as C), and the deformation extends out into the tissue with decreasing effect (C*).

### Metastases can invade along paths of mechanical least resistance

In the simulation results, if the parenchyma was more tolerant of deformation (higher values of *d*_max_) but readily deformable (lower *r*_E_), then a sufficient amount of parenchyma could be displaced to encapsulate the tumor. Plastic reorganization (represented by *r*_P_) results in a decreasing elastic force, which in turn slowly relieves the pressure on the tumor cells to allow sporadic proliferation. Continued tumor proliferation into this tough yet well-perfused tissue lead to local “fingering” morphologic instabilities, generally nucleated along paths of mechanical least resistance (e.g., in gaps between parenchyma agents). See *Supplementary Fig S5*. We note that this is consistent with prior predictions made with continuum methods, where fingering morphologic instabilities were observed in growth into stiff, well-perfused tisues [57]. Note that this is also observed in clinical tissues: Fig. 8B shows tumor cells growing in collapsed liver sinusoids–paths of mechanical least resistance (see the yellow highlighted regions in the figure inset).

### Micrometastases may remain dormant in highly elastic parenchyma

When the parenchyma was more tolerant of deformation (higher values of *d*_max_), but readily deformable (lower *r*_E_) with slower plastic reorganization (lower values of *r*_P_), then the tumor becomes encapsulated as when invading along paths of mechanical least resistance, With slower plastic relaxation of the parenchyma, however, the elastic force (and hence the high tumor mechanical pressure) is sustained, leading to tumor dormancy. See Fig. 9. The results show that some rigid tissue regimes can arrest the growth of micrometastases, which also agrees with the experimental observations that solid stress can inhibit the growth of multicellular tumor spheroids [36, 69, 72].

**Figure 9.**
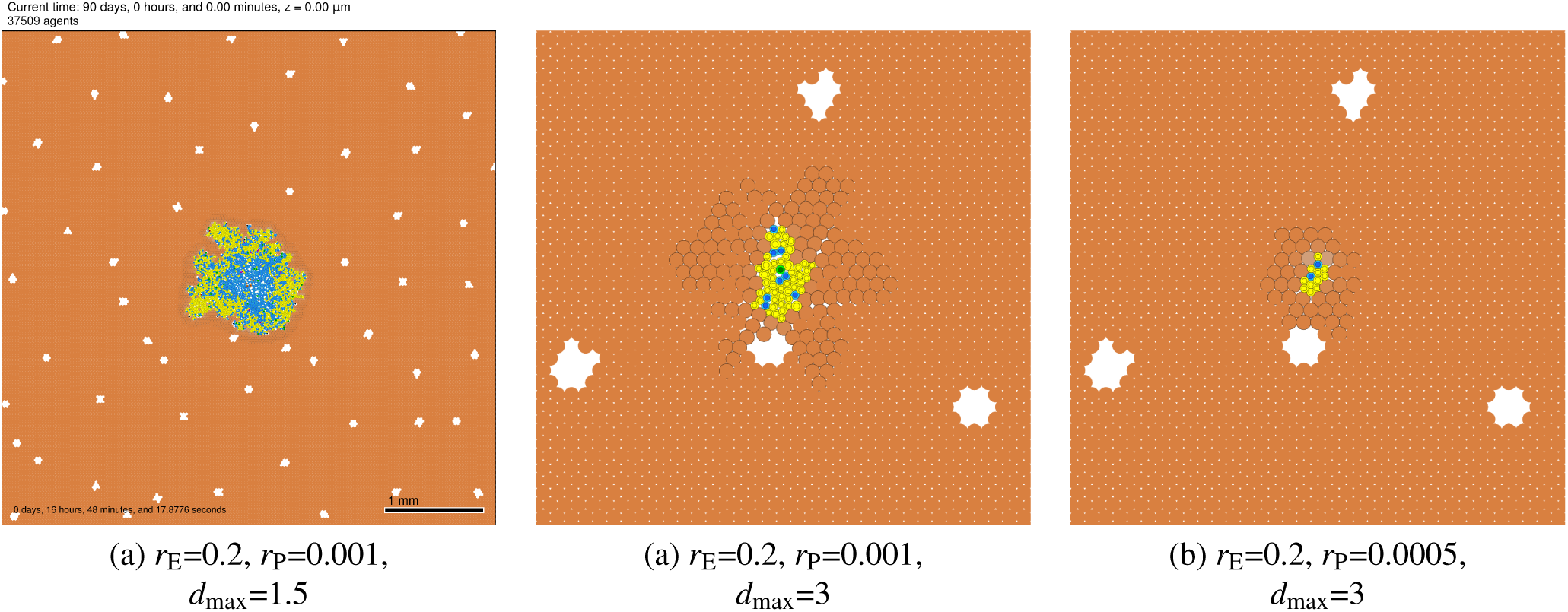
Dormant micrometastases: Metastatic growth into highly elastic parenchyma with high tolerance for deformation can lead to tumor encapsulation. If plastic relaxation is slow, mechanical feedbacks are sustained, leading to slow (but potentially invasive) growth (a) or tumor dormancy (b-c). Cells are colored as described in Fig. 5.

### Changes in parenchyma biomechanics can trigger “reawakening” of dormant micrometastases

To investigate the impact of a change in parenchyma tissue biomechanics (e.g., after an acute injury or illness that drives tissue remodeling, or during aging), we first simulated three tumors for 90 days until reaching dormancy as described above: (*r*_E_ = 0.2 min^*-*1^, *r*_P_ = 0.0005 min^*-*1^, *d*_max_ = 3 *µ*m). We then simulated an additional 90 days of growth under three scenarios: (I: *control*) left the parameters unchanged (top row in Fig. 10); (II: *increased deformability*) reduced *r*_E_ to 0.05 min^*-*1^ (middle row in Fig. 10); and (III: *reduced tolerance to deformation*) reduced *d*_max_ to 1.5 *µ*m (bottom row in Fig. 10).

**Figure 10.**
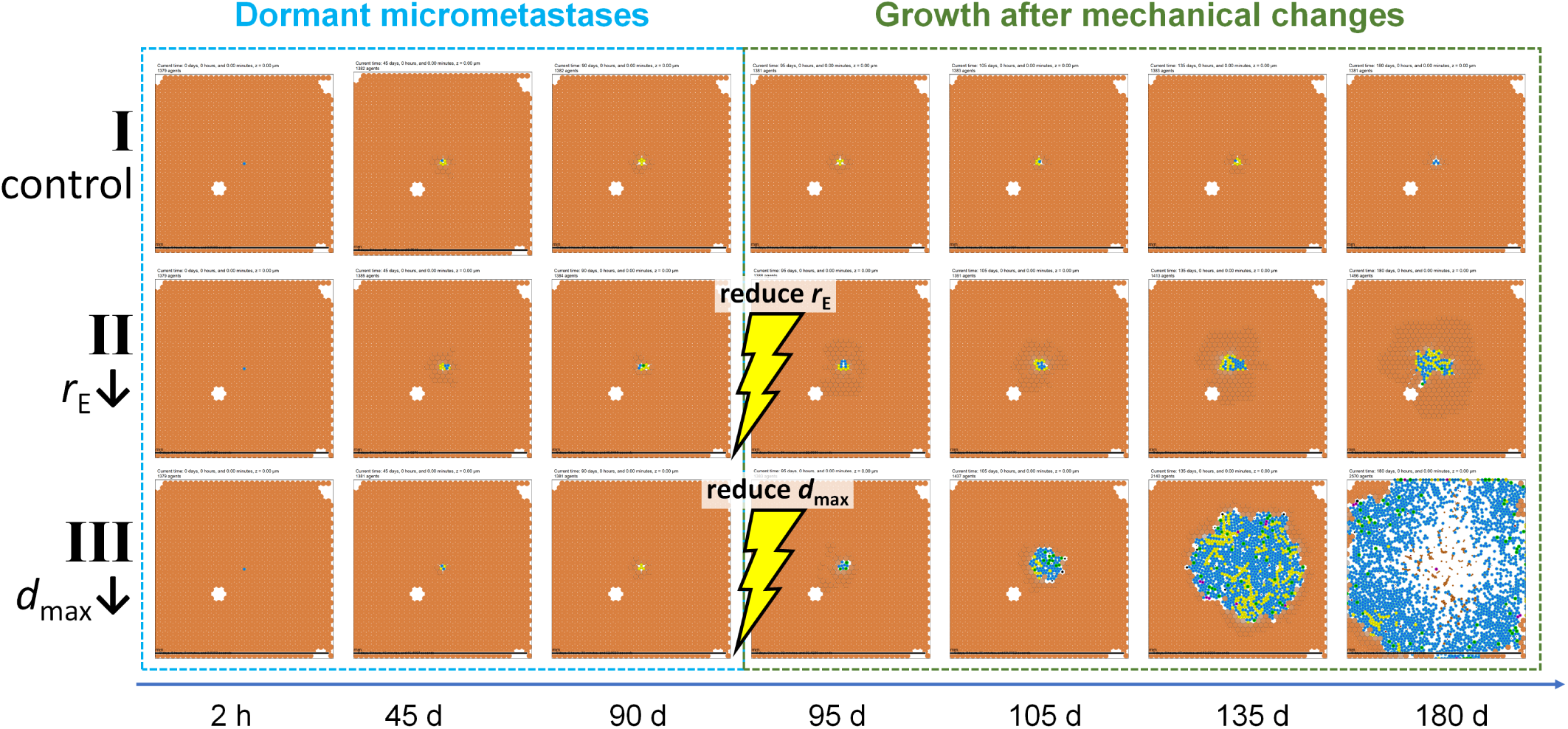
Dormant tumor reawakening after biomechanical changes: Three micrometastases were grown to dormancy over 90 days in identical conditions and then subjected to biomechanical changes in the liver parenchyma before an additional 90 days of growth. Tumor I (control) remained dormant for the extra 90 days. Tumor II (*r*_E_) “reawakened” with slow growth. Tumor III (reduced tolerance to deformation *d*_max_) reawakened to exhibit rapid growth. Cells are colored as described in Fig. 5.

With no changes in the parenchyma (tumor I in Fig. 10), the tumor remained dormant for an additional 90 days. Reducing the elastic parameter *r*_E_ as in tumor II in Fig. 10 allowed the surrounding parenchyme to more easily deform by decreasing the elastic force resisting tumor growth. Under these conditions, the dormant tumor “reawakened” to slow, invasive growth. Reducing the parencyma’s tolerance for deformation *d*_max_ as in tumor III in Fig. 10 resulted in faster liver tissue death, allowing the tumor more easily displace and replace it. In these conditions, the dormant tumor reawakened to rapidly grow through the tissue. Our simulation results showing ECM-induced mechanical restraint on tumor cells are consistent with experimental findings that evaluate the effect of collagen matrix remodeling in aging skin on melanoma metastasis [73–75].

## Discussion

Liver metastases represent the final stage for many types of cancer, with limited clinical options to offset patient mortality. Yet the biomechanical interactions that favor metastases to seed in liver tissue and enhance their growth remain poorly understood. This study presents a modeling framework to systematically evaluate the effects of mechanical interactions between liver parenchyma and tumor cells on metastatic seeding and growth (in combination with tumor cell proliferation and death that vary with substrate availability). The biomechanics of centimeter-scale liver tissue are simulated by coupling a poroviscoelastic (PVE) model with an agent-based model (ABM).

Building upon prior calibration and validation of the PVE model for *ex vivo* liver tissues, we simulated coupled flow and and mechanics in micrometastases and liver parenchyma in *in vivo* conditions. The results for fluid flow and effect on tissue strain show that changes in tumor permeability may have little to no effect when the tumor is modeled as a pressure source *(Supplementary Fig S3* or sink Fig. 3). However, when the tumor is modeled as neutral (neither source nor sink, *Supplementary Fig S2*), fluid flows faster through high permeability tumor tissue and slower through low permeability tissue as compared with normal parenchyma. For the neutral condition, high tumor permeability is also associated with higher tumor stresses (Supplementary Materials). When the tumor is modeled as a pressure sink (Fig. 3), most of the fluid entering the lobule drains through the tumor boundary, so that fluid velocities in central vein regions are negligible. Setting the tumor as a pressure sink also produces a drop in stress and strain at the tumor edge, unlike the spike in stress and strain noted with the other two conditions. Lastly, modeling the tumor as a pressure source produces the highest overall magnitudes of fluid velocity and strain (*Supplementary Fig S3*). The ABM results show that tumors reached smaller sizes when growing in liver tissues that could tolerate large deformations (larged values of *d*_max_) with reduced stress-induced apoptosis for parenchyma. The parenchyma agents imposed larger elastic forces on tumor cells, resulting in mechanobiologic growth arrest, even with readily available oxygen and nutrients. The results overall support the notion that while biotransport is largely advective and quasi-steady in unobstructed parenchyma, it is largely diffusive within micrometastases. Note that these results are difficult to obtain in animal models and prohibitively invasive for patients.

We further implemented an analytical single-lobule approximation to simulate advection-dominated perfusion in a large tissue section, and used this to evaluate whether a tumor model driven solely by growth substrate availability could parsimoniously model metastases with perfusion matched to the PVE model results and analyses. We found that for metastatic growth into an elastic, well-perfused liver tissue, a minimal tissue-scale model must include additional mechanobiologic negative feedbacks on tumor cell proliferation to avoid non-physical overlap of cells, while substrate-dependent death is still needed to model the emergence of necrotic cores. After scaling parenchyma death with mechanical disruption (a simplified model of mechanics-driven apoptosis), we performed a parameter space investigation of the emergent dynamics of this minimal model over a range of tissue elasticity, plasticity, and tolerance of mechanical disruption. The large-scale model exploration showed that tumor-parenchyma biomechanical interactions generate pressure inside tumors and tissue strains in the nearby parenchyma. The model predictions of deformed liver parenchyma around tumor boundaries can be observed in clinical samples colon cancer liver metastases. Matching long-time model behaviors with prior (static) clinical observations provides a dynamical context to understand these results, while also providing a theoretical bridge between known cell-scale cancer biology and clinical observations. Future work to fit this simple mathematical model with more detailed molecular models of signaling and micro-scale ECM interactions could further link known molecular biology to intermediate dynamics and long-time clinical observations.

Overall, the modeling framework presented here provides a platform to explore the mechanobiological conditions that lead to metastatic seeding, dormancy, and growth in liver tissue [76, 77]. Model explorations generate predictions (hypotheses) that we can presently assess qualitatively against known clinical and experimental observations. Future experiments could be tailored to these predictions could potentially further test the predictions quantitatively. The simulation results show that micrometastases may remain dormant in highly elastic parenchyma. These results are consistent with observations using a 3D *ex vivo* hepatic microphysiological system, which showed that the proportion of MDA-MB-231 breast cancer cells were more likely to enter dormancy in more deformable tissues compared to a stiffer polystyrene scaffold [78], analogous to the simulations where parenchyma agents could tolerate greater displacement. The simulation studies found that changes in parenchyma biomechanics can trigger “reawakening” of dormant micrometastases, with metastatic growth representing an invasion along paths of mechanical least resistance. In particular, simulations showed that changes in liver tissue deformability could occur when injury or aging damages the structural integrity of extracellular matrix; prior experimental cancer biology shows that such changes could occur by degradation of collagen and elastin or decreased cross-linking [9, 75, 79]. The simulations also predict that a decrease in the overall resiliency of hepatocytes due to injury or illness may additionally lead to liver tissue changes favoring metastatic growth; this is also plausible in light of known liver biology. For example, aging as well as fatty liver disease are known to reduce the regenerative capacity of the liver parenchyma [79]. Illness or injuries that compromise non-parenchymal cells (e.g., Kupffer cells, leukocytes, or sinusoidal endothelial cells) could potentially contribute to both of these scenarios [79]. In [55], a microphysiologic model system found that the presence of intact non-parenchymal cells was associated with increases in cancer attenuation signals, decreases in pro-inflammatory signals, and reduced colonization by metastatic breast cancer cells.

It is well known that aging-associated cellular and molecular changes in non-cancerous cells can contribute to a microenvironment favoring tumor progression, including biophysical changes to the ECM [75]. In particular, mechanical alterations including strain-stiffening, increased cross-linking, and fiber realignment are thought to be critical in promoting cancer growth and invasiveness [80, 81]. Further, ECM stiffening and remodeling is promoted by the secretion of factors from stromal components that become senescent with age [82]. In turn, the ECM changes affect the immune system response [83]. Interestingly, while aging can lead to increased local tissue stiffness and fibrosis, dysregulation of ECM constituents crosslinking can also yield loose, disorganized fragmented collagen [9]. These changes facilitate mechanically weak tissues, which can promote tumor invasion. Thus, aging can lead to both tissue stiffening and softening, which in their own ways facilitate tumor progression and invasiveness. These biological observations are recapitulated by the simulation results of this study. The effect of tumors growing to a steady size due to mechanical feedback (Fig. 5) realistically relates to tumor dormancy. Once injury or aging mechanically weaken the liver tissue, dormant micrometastases could resume growth. If the mechanical changes result in fibrosis that leads to local tissue stiffening, which upon plastic reorganization of the ECM is projected by the model to induce a “fingering” tumor invasion pattern to circumvent the mechanical constraints (*Supplementary Fig S5*).

Our results showing ECM-induced mechanical restraint on tumor cells are consistent with findings by [73], who performed experiments to evaluate the effect of collagen matrix remodeling in aging skin on melanoma metastasis and immune response. A mathematical model was implemented to evaluate the effect of inter-fiber crosslinks on the deformation of collagenous ECMs and the two-way interactions between the strain-stiffening behavior of fibrous ECM and cellular contractility. The simulations showed that HAPLN1, the most abundantly secreted ECM constituent by fibroblasts, significantly alters matrix organization and inhibits cell motility [73]. Related to these interactions, a mechanochemical free-energy-based method was recently developed to elucidate the two-way feedback loop between stress-dependent cell contractility and matrix fiber realignment and strain stiffening [74], validated by analyzing the invasion of melanoma cells in collagen matrix of different concentration. The model was able to find a critical stiffness (EM=0.15 kPa) required by the matrix to break intercellular adhesions and initiate cell invasion. Below this critical stiffness, spheroids kept stable as driving forces could not overcame cell adhesion, while the driving force could break the adhesions and induce invasion when the stiffness was beyond the critical value. In addition, it was found that enhancing collagen concentration in the ECM increased the density of the cell-adhesive ligands, which also in return promoted cell invasion [74].

The modeling framework presented here has several limitations, as only a subset of key biology is represented. Integrating additional biological details could refine the results, improve quantitative matching with targeted experiments, and potentially lead to further insights. For example, incorporating additional key cell types which are known to modulate metastatic inception and progression (e.g., fibroblasts, hepatic endothelial cells, and immune cells) would improve our simulation of cell-cell and cell-matrix interactions, similarly to recent modeling efforts that explored the role of M1 macrophage-induced cytotoxicty in well-vascularized tissues such as liver [42]. Immune and fibroblast cell agents recently developed for SARS-CoV-2 research could readily be adapted to the agent-based model. Similarly, varying tumor cell proliferation and apoptosis to the ECM and integrin signaling could better connect the model with known molecular cancer cell biology [85]. The structure of the liver could be more detailed, e.g., refining the microstructure of sinusoids to more faithfully represent the organ structure. In spite of these limitations, the framework shows that the synergistic integration of multiple modeling approaches may provide insight into cancer behavior. In particular, the model enables evaluating the impact of metastases on perfusion flow, the accumulation of stresses and strains in the local parenchyma, and how these effects altogether support the notion that biomechanical interactions between metastatic tumors and their microenvironment have a role in cancer behavior. Although previous work has investigated molecular-scale feedback mechanisms (e.g., inflammation and other cross-signaling), few studies have focused on purely mechanical feedbacks. Our results suggest that conditions exist for which mechanical feedback alone could explain observations of metastatic progression in the liver, which should reinforce previously studied bioechemical feedbacks. Further, the discrete model formulation presented herein was able to recreate continuum-level pressure and mechanics conditions with relatively simple cell rules, compared to more complex continuum-scale formulations (e.g., Cahn-Hilliard based models) [67]. This relative simplicity offers the promise of easier translation of experimental observations to model cellular agent hypotheses to observe macro-scale effects.

The proposed modeling framework enables evaluation of biomechanical interactions (adhesive, repulsive, and elastic forces on short time scales, and plastic reorganization on longer time scales) on metastatic tumor cell seeding and growth in liver tissue. The simulation results indicate that there exist conditions for which these interactions may arrest micrometastatic growth and prevent newly arriving cancer cells from establishing new foci. Future work will examine in more detail the role of tissue microstructure, such as sinusoids, on metastatic progression. In particular, CRC metastases create glandular structures that are not fully solid. The associated microlumens could provide some stress relief and allow cell tumor and parenchymal cells to intermix, rather than undergo more homogenous compression. An improved understanding of these as well other tumor structural characteristics when interacting with liver tissue could eventually lead to patient tumor-specific therapy to prevent and reverse metastatic seeding and progression.

## Methods

### Poroviscoelastic (PVE) Model

This section illustrates the theoretical details of the PVE model, model assumptions, computational domain and numerical implementation details. The material properties and boundary conditions as well as fluid velocity, strain, stress, and pore fluid pressure results are included in the Supplementary Materials.

#### PVE model overview

A detailed mathematical description of the mechanics of poroviscoelastic materials was published by [86], and the application of poroviscoelastic models to liver tissue has been demonstrated by [87–89] among others. The summary given here is adapted from [29, 89, 90]. A biphasic material consists of a linear elastic solid phase and an incompressible fluid phase, where relative motion between the two phases produces rate-dependent (i.e. viscoelastic) behavior [91]. If the fluid phase is inviscid, biphasic theory is equivalent to poroelasticity. Poroviscoelasticity extends poroelasticity by modeling the solid phase as viscoelastic, rather than linear elastic [92]. Equations 1-7 comprise biphasic material mechanics, Equations 8 and 9 provide for viscoelasticity of the solid phase, and Equations 10 and 11 finish the constitutive description.

The conservation of mass for incompressible solid and fluid phases can be written as:

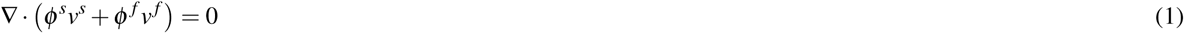

where *s* and *f* indicate solid and fluid phase, respectively, *ϕ*^*s*^ and *ϕ* ^*f*^ are solid and fluid volume fractions, and *v* is velocity. If inertial forces are negligible compared with internal frictional forces, the conservation of linear momentum can be written as:

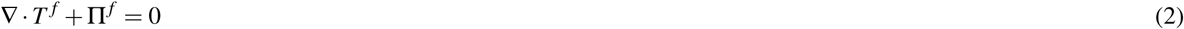

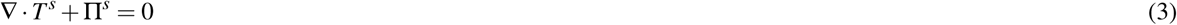

where *T* ^*f*^ and *T*^*s*^ are Cauchy stress tensors and П ^*f*^ and П^*s*^ are viscous drag forces arising from the interaction between the fluid and solid phases. П ^*f*^ and П^*s*^ are proportional to their relative velocities and inversely proportional to hydraulic permeability, *k*:

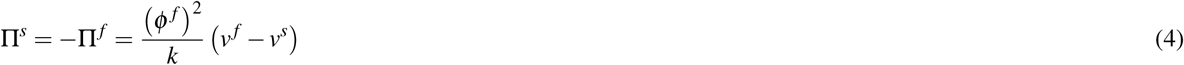

Stress tensors in the fluid and solid phases can be written as:

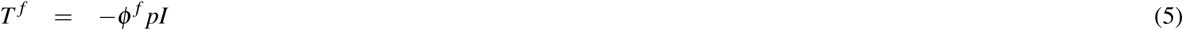

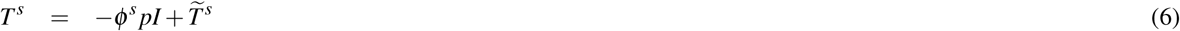

where *p* is hydrostatic pressure and 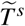 is apparent solid stress due to solid matrix deformation [91–93].

If the solid phase is linear elastic, then 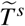 can be written as:

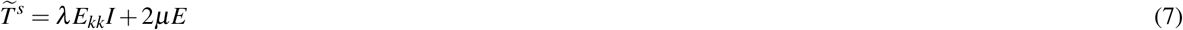

where *E* is the infinitesimal strain tensor and *λ* and *µ* are Lamé constants. To account for intrinsic viscoelasticity of the solid phase, the solid stress tensor can be replaced with:

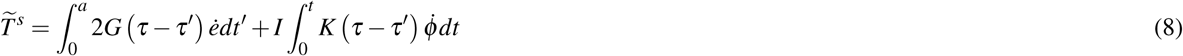

where *G* and *K* are elastic shear and bulk relaxation functions, *e* is deviatoric strain, and *ϕ* is volumetric strain [92][93]. The relaxation functions can be defined by a Prony series expansion

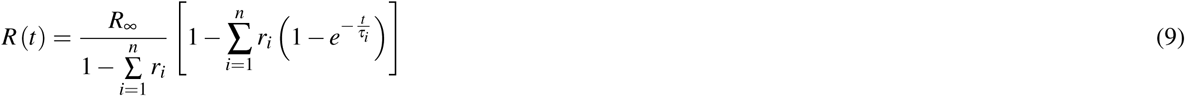

where *R* is the time-dependent modulus, *R*_∞_ is the long-term modulus, and *n, r*_*i*_, *τ*_*i*_ are Prony series constants. In the Abaqus implementation in the present work, the material behavior was defined by specifying the hydraulic conductivity (*K*), the specific weight of the liquid (*γ*), the Prony series constants, a long-term Young’s modulus (*E*_∞_), and Poisson’s ratio (*ν*) so that:

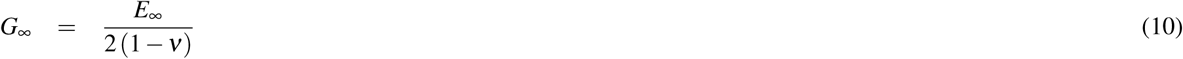

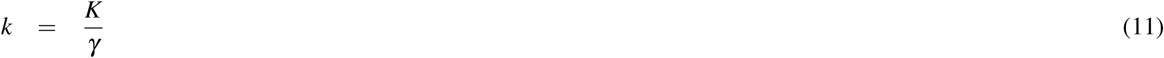

#### Starting model assumptions

It is assumed that necrosis is a source of fluid pressure as necrotic cells release fluid. It is also assumed that proliferating cells are a fluid pressure sink, as proliferating cells grow in part by absorbing fluid. Accordingly, the medium (400 *µ*m diameter) tumor is considered as a net pressure source, reflecting a release of fluid from the necrotic core that exceeds the uptake of fluid by cells in the viable rim. For comparison, in a different model iteration the medium tumor is considered as neutral, neither a source nor a sink, to reflect the possible condition that fluid release and fluid uptake are in balance. The small (200 *µ*m diameter) tumor is considered as a net pressure sink reflecting more proliferation than necrosis. Lastly, the tumor seed (50 *µ*m diameter) was considered as neutral, as a tumor of this size would consist of only a few cells and its contribution as a fluid source or sink may be neglected.

#### Model geometry

Model geometry is given in Fig. 11. Lobule dimensions and vessel radii (Table 2) were obtained from histological images in published literature [94–97]. Dimensions for ℓ and *h* are reported as the distance between the centers of the pre-tPVs (pre-terminal portal veins), and *d* is reported as the farthest distance between model faces in the z and -z direction. Each quarter-lobule model contained from 299,258 to 677,457 quadratic tetrahedral elements (Abaqus element type C3D10MP). Simulation time was 150 seconds and the minimum time step was 0.15 seconds.

**Table 2.** Quarter lobule dimensions adapted from Nishii et al. [29]

| Dimensions ($\mu\text{m}$) | |
| --- | --- |
| CV radius | 26.5 |
| tPV radius | 16 |
| pre-tPV radius | 23 |
| Edge length ($\ell$) | 350 |
| Edge width ($w$) | 303.1 |
| Edge height ($h$) | 350 |
| Lobule depth ($d$) | 30 |

**Figure 11.**
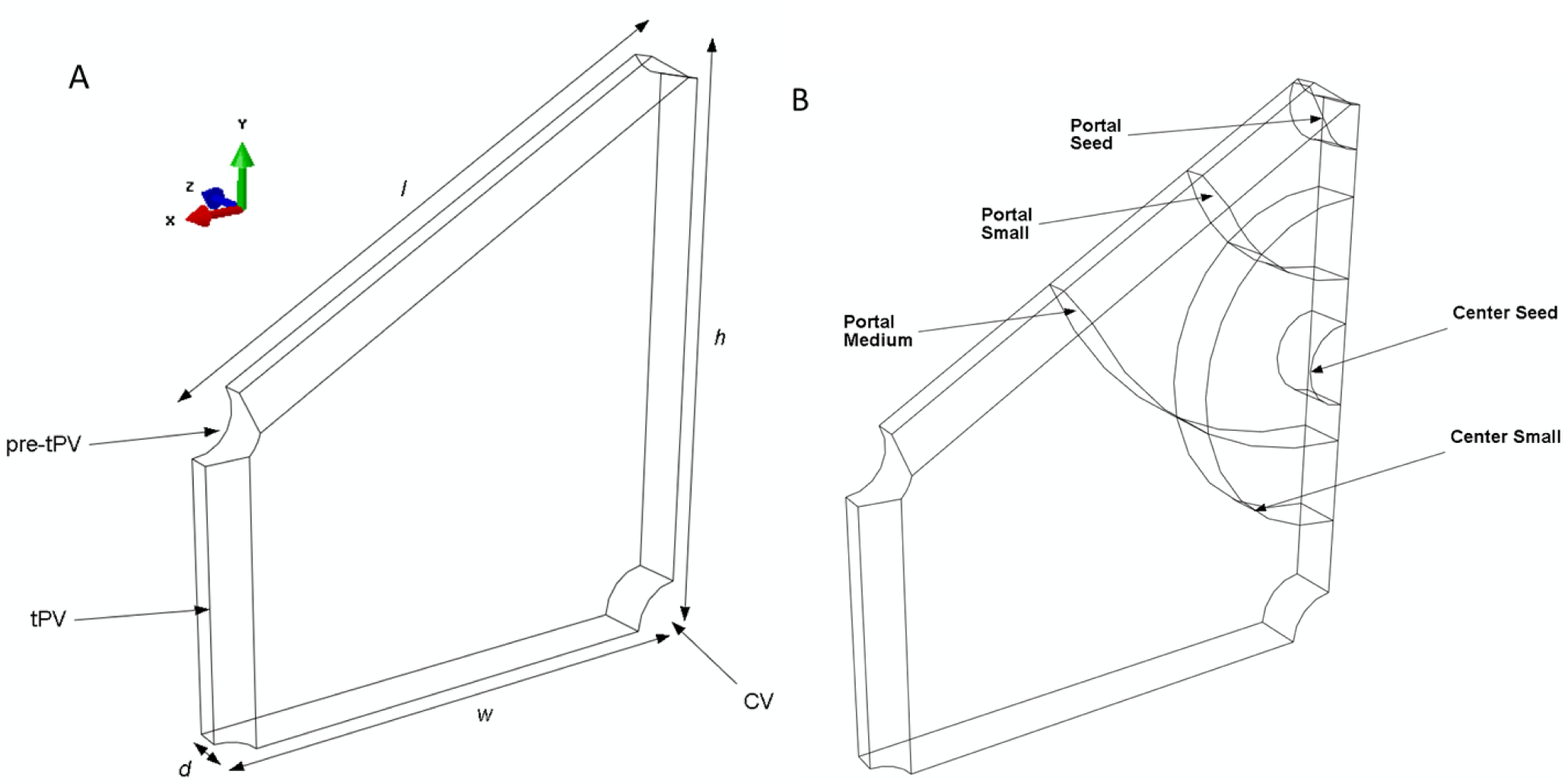
The PVE computational model. Model geometry (**A**) and simulated tumor sizes and positions (**B**). (**A**) Quarter liver lobule showing locations of the pre-terminal portal vein (pre-tPV), terminal portal vein (tPV), and central vein (CV). Dimensions ℓ, *w, h*, and *d* are given in Table 2. (**B**) Tumors of different sizes (seed, small, medium) were simulated for different lobule locations, including the midpoint of the central-portal axis (termed the “center” position) and in the portal triad (termed the “portal” position).

#### Numerical implementation

In order to model the liver microenvironment, a three-dimensional poroviscoelastic finite element model of a quarter liver lobule (Fig. 11) was created in Abaqus (v6.12, 3DS Dassault Systems, Waltham, MA), modified from the prior work of Nishii et al [29]. A poroviscoelastic material model was selected because this constitutive description of liver tissue mechanics has been shown to capture the viscoelastic stress-strain behavior of the solid matrix as well as pressure-driven flow of fluid through the matrix pores [29]. To reduce computational time, only one-fourth of the hexagonal lobule geometry was simulated, and symmetry boundary conditions were applied to X, Y and Z planes (Fig. 11). Model thickness in the Z direction was set to 30 *µ*m to approximate the diameter of one hepatocyte. Dimensions and parenchyma material properties were obtained from literature [29] as described above. Once the base model geometry was established (Fig. 11A), three different sized tumors denoted as seed (50 *µ*m diameter), small (200 *µ*m diameter), and medium (400 *µ*m diameter) tumors were included. Tumors were also modeled as poroviscoelastic solids, but with greater stiffness and altered hydraulic conductivity compared with normal parenchyma (these material properties are described in Supplementary Materials). Tumors were placed at the midpoint of the central-portal axis (termed the “center” position) and in the portal triad (termed the “portal” position). Tumor sizes and locations are shown in Fig. 11B. Each model run simulated only one tumor size at one location.

### Biotransport modeling in BioFVM

In [68], Macklin and coworkers developed *BioFVM* to simulate diffusive biotransport for large 2-D and 3-D biological problems with off-lattice, cell-based sources and sinks. Specifically, if ***ρ*** is a vector of diffusible substrates with diffusion coefficients ***D***, decay rates ***λ***, bulk source rates ***S*** (which saturate at target densities *ρ*^*\**^), and bulk uptake rates ***U***, then problems of the following form are solved:

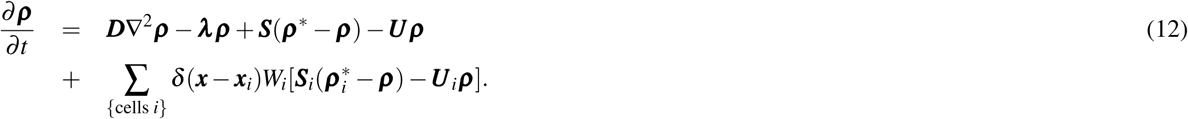

with Neumann (zero flux) boundary conditions. Here, for a collection of cells (indexed by *i*), with centers 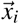 and volumes *W*_*i*_, their secretion rates are 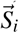 (with saturation densities 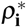) and their uptake rates are 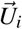.

The method tested second-order accurate in space, first-order accurate in time, and numerically stable in typical parameter regimes for cancer and tissue biology. The method was tested for scalability on systems of millions of off-lattice agents, millinos of voxels, and dozens of diffusing substrates. Setting Δ*t* = 0.01 min and Δ*x* = 20 *µ*m maintained relative errors below 5% for diffusion, decay, uptake, and source rates typical for cancer and tissue biology. See [68] for full details on the numerical method, implementation, and convergence testing. In [61], BioFVM is used as the default biotransport solver, with the system performed accurately and stably even on systems of moving cells.

A useful feature of BioFVM are *Dirichlet nodes*: any voxel *j* centered at position 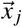 can have its substrate values set to ***ρ*** _*j*_ to approximate Dirichlet conditions on arbitrary geometries. Moreover, one can set or unset any computational voxel as a Dirichlet node at any simulation time, or overwrite the values of ***ρ*** _***j***_. This feature will be used to approximate quasi-steady conditions in selected tissue regions.

### Multicellular modeling in PhysiCell

In [61], Macklin and coworkers developed *PhysiCell* for off-lattice agent-based simulations in multicellular systems biology, with a particular focus on cancer (see [98] for details of cell-based computational modeling in cancer biology). Each cell is a software agent that moves under the balance of mechanical forces under the “inertialess” assumption as in earlier work by Drasdo, Höhme and colleagues [99]. Each cell agent with index *i* has a phenotype 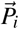: a hierarchically structured set of properties including state and rate parameters for cell cycling, death, volume changes, motility, mechanics, and secretion/uptake. Each cell can sample the microenvironment (modeled with BioFVM; see Methods) and any element of 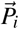 based upon these local microenvironmental conditions. Each cell agent can have custom functions assigned to it; extensive use of these functions is made to model custom mechanobiology, as when obtaining an approximate analytical solution to the quasi-steady distribution of cell substrates. PhysiCell has been extensively tested on a variety of multicellular systems problems with 10^5^ to 10^6^ cells. Its computational cost scales linearly with the domain size and number of simulated agents. See [61] for full algorithmic detail, numerical testing, and a variety of examples.

#### Implementation details

In this study, we used PhysiCell Version 1.2.1 [61] with two agent types: tumor cells (individual cancer cells arriving to colonize and grow in the parenchyma) and parenchyma agents (small portions of parenchyma tissue). For this study, we do not model the fine lobular structure (particularly hepatic cords and sinusoids), but this is a continuing topic of interest for future study.

We use the built-in “Ki67 Advanced” cell cycle model (with default parameters), where quiescent Ki67-cells can transition to a pre-mitotic Ki67+ state, grow, and divide into two Ki67+ daughter cells. These daughter cells grow and exit the cycle to the Ki67-state. Each cell can grow, apoptose, or necrose based upon user-defined functions. Except as described later, we use the standard, built-in cell mechanics. We also begin the built-in oxygen-dependent proliferation and necrosis: notably, the Ki67-to Ki67+ transition rate increases with oxygen availability above a minimal hypoxic threshold, and the necrotic death rate increases below the threshold. See [61] for further details and default parameter values.

For parenchyma agents, we set cycling and apoptosis to zero to model a homeostatic tissue in the absence of tumor cells. This investigation does not focus on single-cell invasion and EMT, but rather post-MET metastatic seeding and growth; we disable cell motility accordingly. The full source code is available as open source (see *Appendix-Code availability* in the *Supplementary Materials*), and an interactive, online version can be run at https://nanohub.org/tools/pc4livermedium.

### Generating large 2-D virtual liver tissues

This study required large 2-D (thin 3-D) sections of virtual liver tissue. Here, we describe a straightforward approach to generate a synthetic tissue dataset, based upon a Voronoi approximation of hepatic lobule packing. To generate a large liver tissue:

1. Central veins (CVs) are randomly placed across the tissue, with mean density ∼ 2 ×10^*-*6^*µ*m^*-*2^ (to ensure a mean hepatic lobular diameter of approximately 800 *µ*m, similar to analyses in whole-lever models such as [29, 96]);
2. Any closely-spaced pairs of CVs are replaced with a single CV placed at their midpoint; and
3. Remaining space is filled with (hexagonally-packed) parenchymal agents.

As needed, the lobular boundaries can be estimated by generating a Voronoi mesh from the central vein positions.

A MATLAB script is included in the github repository (see *Appendix-Code availability* in the *Supplementary Materials*) to implement this algorithm; it has the random seed hard-coded to reproduce the liver tissue in this manuscript.

### Model refinements to avoid non-physical behaviors

We combined the analytical approximation for advection-dominated non-tumor regions from the Section “Approximate oxygen distribution in large tissue sections” in Supplementary Materials with the BioFVM diffusion solver in PhysiCell: in any voxel that does not contain a tumor cell, we assume the tissue has intact flow (as in the case of not tumor cells or very small micrometastases) and apply the analytical approximation as a Dirichlet node. Prior to tumor seeding, every computational voxel is a Dirichlet node, and so the initial oxygenation matches the profile shown in the Supplementary Materials. We model any voxel containing tumor cells as permanently disrupted, as suggested by the PVE modeling results. To do this, we permanently remove the Dirichlet node from that voxel and use the usual diffusion solver in BioFVM.

### Pressure-regulated tumor cell proliferation

In a form of mechanosensing, cells can sense compression and down-regulate cell cycling to resist overcrowding (e.g., [100, 101]). As inspired by [70] and introduced in [61], cell agents can use their potential functions to construct the local compressive pressure *p*, which we normalize to 1 for a cell in a densely packed 3-D tissue (12 similarly sized neighbors in dense packing) or 0.5 in 2D (6 similarly sized neighbors in dense packing). We can therefore combine this pressure with the prior oxygen-depenent cycle entry for a mechanosensory cycle arrest:

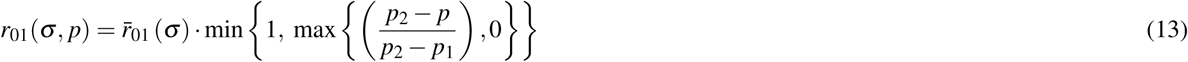

where 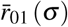 is the oxygen-dependent transition rate from the Ki67-phase to the pre-mitotic Ki67+ cycle phase. Assuming that tumor cells can tolerate greater compression than non-malignant cells, we set *p*_1_ = 0 and *p*_2_ = 1. In this functional form, the “regular” oxygen-dependent cycle entry rate is scaled by the pressure. If *p* ≥ *p*_2_, cycling is arrested in the Ki67-phase, and if *p* ≤ *p*_1_, then cycling continues without mechanosensory-based restriction.

### Elasto-plastic mechanics and mechanosensing in the parenchyma

We model the parenchyma as a collection of parenchyma agents, rather than modeling at the level of detail of individual hepatocytes and endothelial cells. Each parenchymal agent follows regular adhesion-repulsion mechanics (with a position 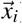) based on the standard PhysiCell potential functions [61]. In addition, we model mechanical interactions with the underlying ECM based upon an earlier model we developed in [33]. Each agent *i* with position 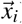 and radius *R*_*i*_ is attached to the ECM at 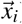ECM and experiences an additional elastic force that resists displacement (deformation) of the cell away from 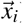ECM. Let 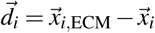, and 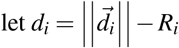 denote the elastic deformation. Then if *r*_E_ is a coefficient for the magnitude of an elastic restorative force, then we model (after the “inertialess” balance of force as in [33, 61, 99]) this restorative force an additional term in the cell’s velocity:

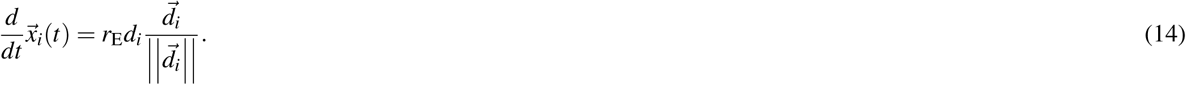

Following [31–33], if a tissue remains in a strained state, it can undergo plastic reorganization. We model this relaxation process by evolving 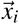_,ECM_ towards 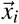 with a (slower) relaxation rate parameter *r*_P_:

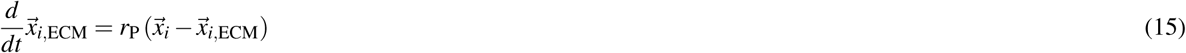

Lastly, we assume that parenchymal agents undergo apoptosis if their deformation (displacement) *d*_*i*_ exceeds a maximum tolerated deformation (*d*_max_). Taken together with the modeling tumor-parenchyma mechanobiologic feedbacks:

1. The elasto-plastic parenchyma model allows a growing micrometastasis to strain the liver tissue, potentially to the point of local tissue death. This can free space for the micrometastasis to expand.
2. The elasto-plastic parenchyma model imparts a resistive force on the cells in a growing micrometastasis, which can lead to compression and thus down-regulate cycling to slow the growth of the metastasis.

An interactive version of this model can be run online at https://nanohub.org/tools/pc4livermedium.

### Computing

We saved simulation outputs (SVG images and MultiCellDS simulation snapshots [61, 102]) every 6 hours for each run. The simulations were performed on the Big Red II supercomputer at Indiana University. Big Red II has 344 XE6 (CPU-only) compute nodes and 676 XK7 “GPU-accelerated” compute nodes, providing a total of 1020 compute nodes, 21,824 processor cores, and 43,648 GB of RAM. Each XE6 node has two AMD Opteron 16-core Abu Dhabi x86-64 CPUs and 64 GB of RAM. Each model was run on a single node using 32 threads. All jobs (270 runs) were submitted as a batch. Each simulation completed in approximately 22 hours, and with a total wall time of approximately of 292 hours.

### Creating interactive cloud-hosted PhysiCell models with xml2jupyter

In [103], we introduced *xml2jupyter*, an open source Python package that can map model parameters from PhysiCell’s XML configuration file to Jupyter widgets, which can automatically create a Jupyter-based graphical user interface (GUI) for any PhysiCell-based model, including user-friendly widgets to configure and run the model and visualize results within the notebook. In addition, the Jupyter notebooks and the accompanying simulation executable can be hosted on nanoHUB [104] as a cloud-hosted, interactive version of the simulation model. This facilitates multi-disciplinary communication of the model without need for downloading, compiling, and running the code.

The agent-based model presented herein is available as a nanoHUB app [105] at https://nanohub.org/tools/pc4livermedium. See *Supplementary Fig S6* for sample screenshots from typical use.

### Histopathology

Tumor tissues were received from colorectal cancer patients under Institutional Review Board (IRB) approval at the Norris Comprehensive Cancer Center of USC. All subjects gave their informed consent for inclusion before they participated in the study. The study was conducted in accordance with the Declaration of Helsinki, and the protocol was approved by the IRB Ethics Committee of USC (Protocol HS-06-00678; approval date 08-02-2019). The USC Translational Pathology Core processed the formalin-fixed, paraffin embedded (FFPE) tissue biospecimens. Tissue sections were prepared and stained with hematoxylin and eosin (H&E). The slides were then imaged using the Olympus VS120 Slide Scanner.

## Acknowledgements

The authors acknowledge funding by National Science Foundation (YW, PM: 1720625), the Breast Cancer Research Foundation (PM), the Jayne Koskinas Ted Giovanis Foundation for Health and Policy (PM), the National Cancer Institute (EB, KN, HBF, SM, JLS, PM: 1R01CA180149; SM, PM: 1U01CA232137). We would like to acknowledge the Miami Redhawk Cluster and the Ohio Supercomputer Center for access to computing resources to complete this work. We would also like to thank Dr. Jens Mueller of the Miami University Research Computing Center for assistance with Abaqus model development. The authors gratefully acknowledge access to FutureSystems at the Digital Science Center, Luddy School of Informatics, Computing, and Engineering, and to the Big Red II supercomputer at Indiana University, Bloomington. We would like to thank the USC Translational Pathology Core and Roy Lau (USC) for assistance with patient tissue processing, staining, and imaging. We thank Randy Heiland (IU) for support in creating and refining the nanoHUB liver app. We thank John Metzcar (IU) for helpful comments on the manuscript.

## Author contributions statement

SM, JLS, HBF and PM conceived the study and obtained funding.

EB and KN performed all poroviscoelastic (PVE) simulations and created corresponding figures.

JLS directed the PVE simulations and performed writing on PVE results.

PM, HBF, SM, and YW developed the agent-based model (ABM).

YW performed all ABM simulations and created the corresponding figures.

PM directed the ABM work and the overall study.

SM contributed the liver histopathology image and analysis.

YW, HBF, SM, JLS, and PM wrote the manuscript.

All authors approve the submission.

