## Supplementary Materials for "Impact of tumor-parenchyma biomechanics on liver metastatic progression: a multi-model approach"

#### Additional Figures

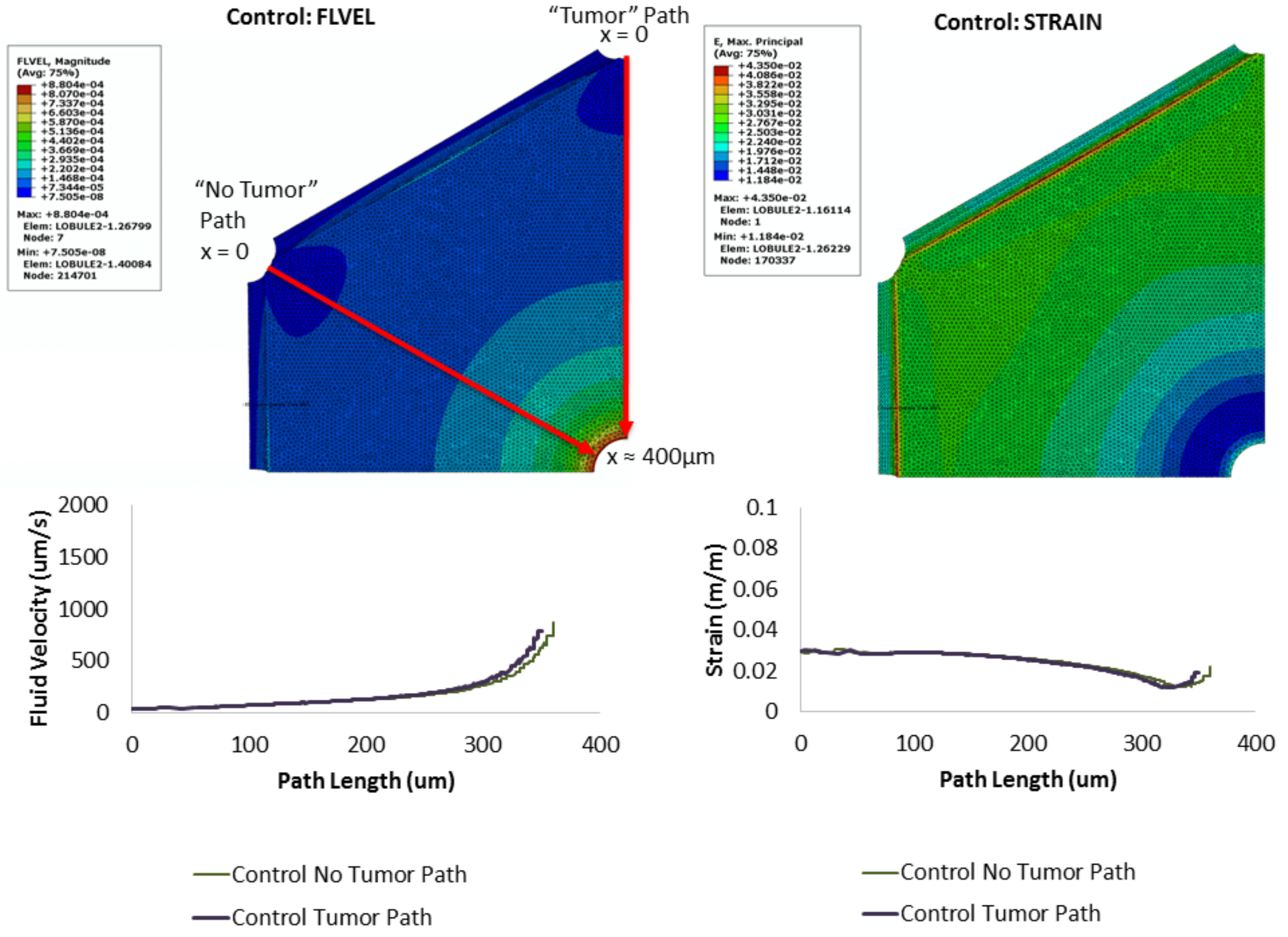


**Figure S1. Hepatic flow: no tumor.** The fluid velocity (left plots) and strain (right plots) are plotted across the lobule (top plots) and along "tumor" and "no tumor" paths (bottom plots). Flow accelerates as fluid approaches the central vein, whereas strain slightly decreases.

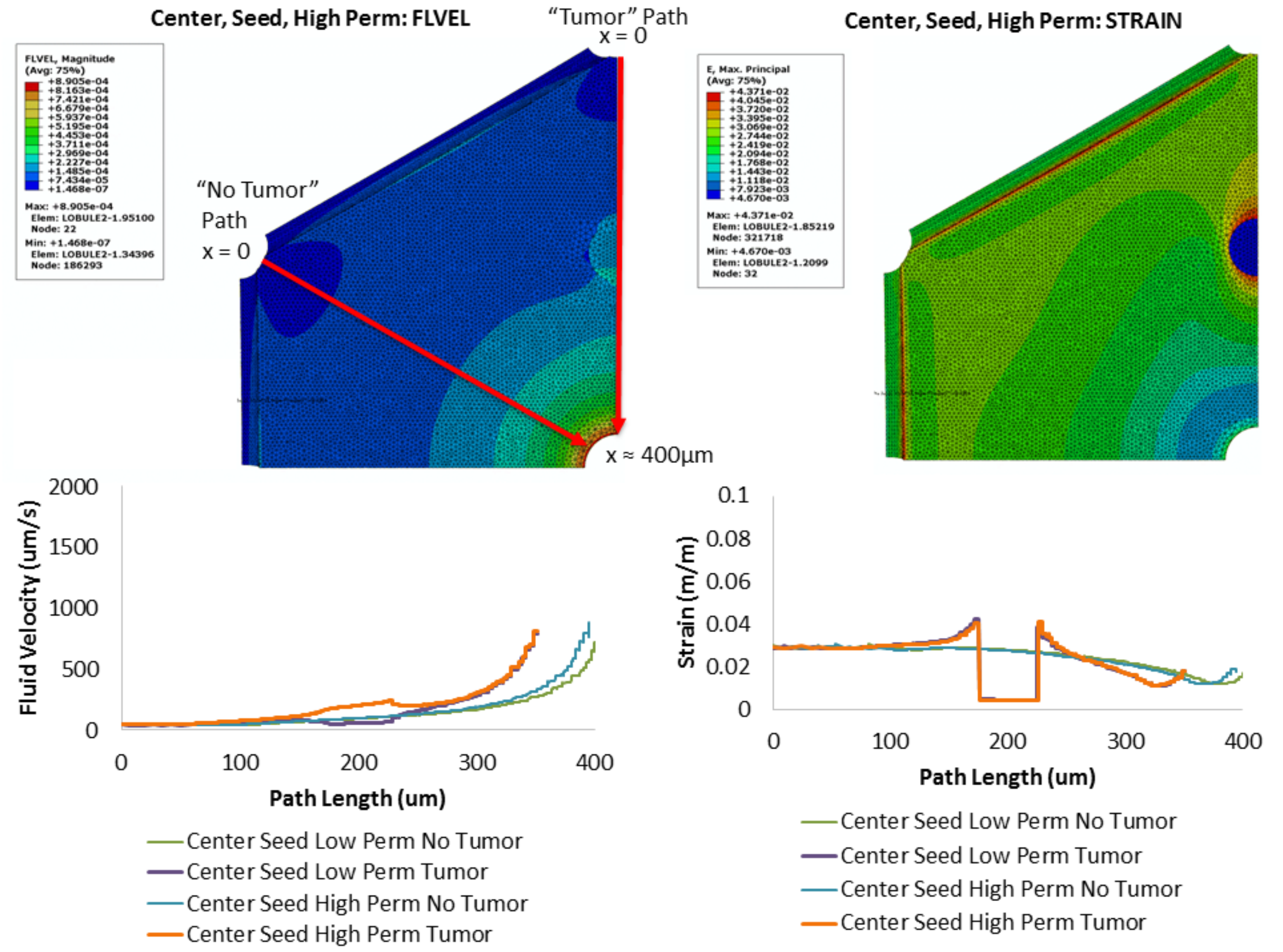


**Figure S2.** **Hepatic flow: small tumor seeds.** A 50 $\mu$m micrometastasis is seeded between the portal triad and central vein, with neutral pressure difference at the tumor boundary. The fluid velocity and strain along the non-tumor path are largely unaffected by the nearby tumor, whereas the fluid velocity increases slightly in highly permeable tumors, and decreases in less permeable metastatic seeds (left plots). The tissue strain is very low within the tumor but increases in the parenchyma immediately surrounding the tumor seed (right plots).

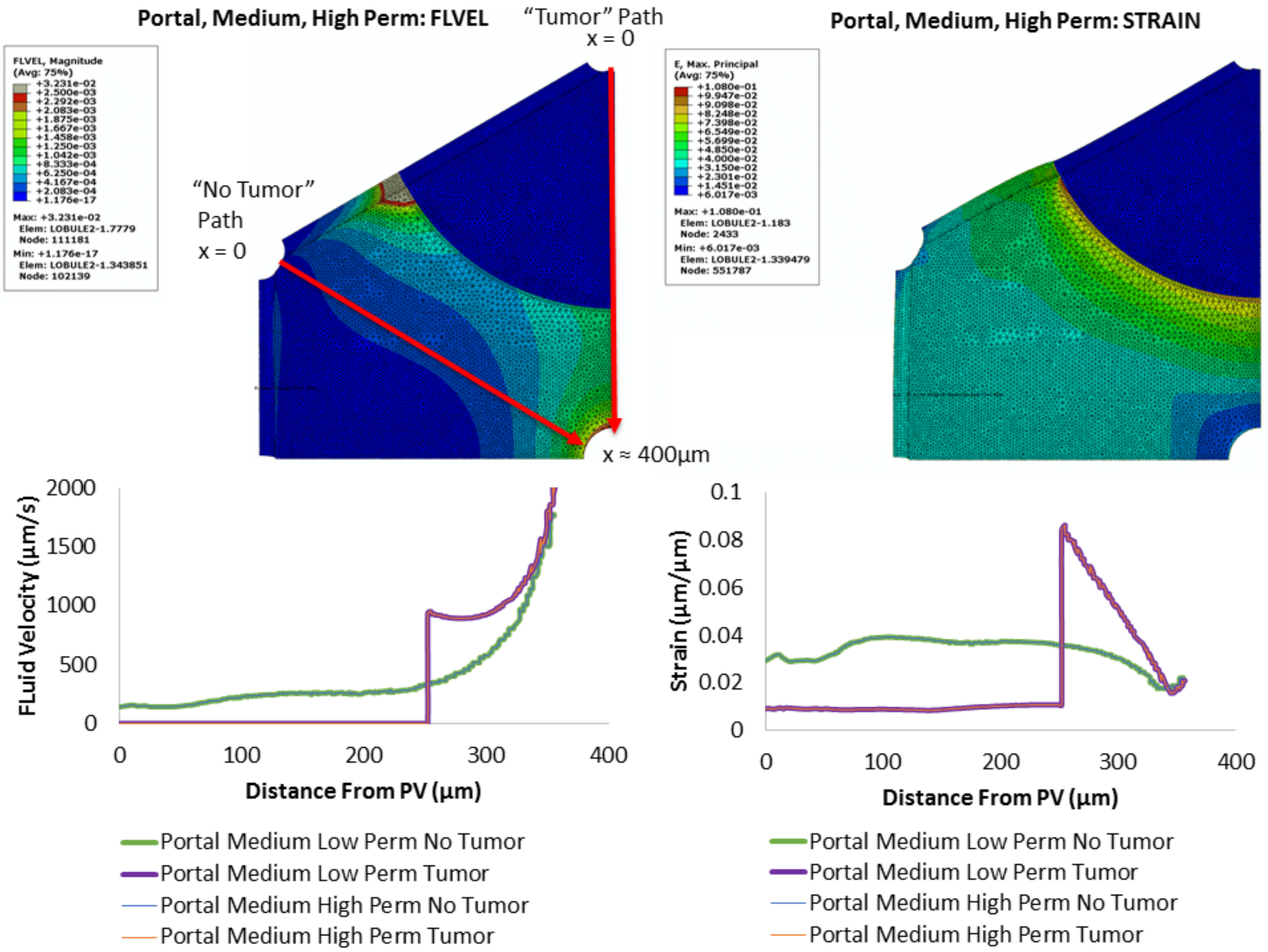


**Figure S3. Hepatic flow: medium micrometastasis.** A 400-$\mu$m micrometastasis disrupts a hepatic lobule near a portal triad, with tumor treated as a net pressure source. The fluid velocity increases in the parenchyma near the tumor but abruptly drops to zero inside the tumor. There is significant tissue strain in the parenchyma that peaks at the tumor edge and extends significantly into the surrounding tissue before dissipating. These results hold for both high and low tumor tissue permeability, and for neutral tumor pressure (results in supplementary materials.

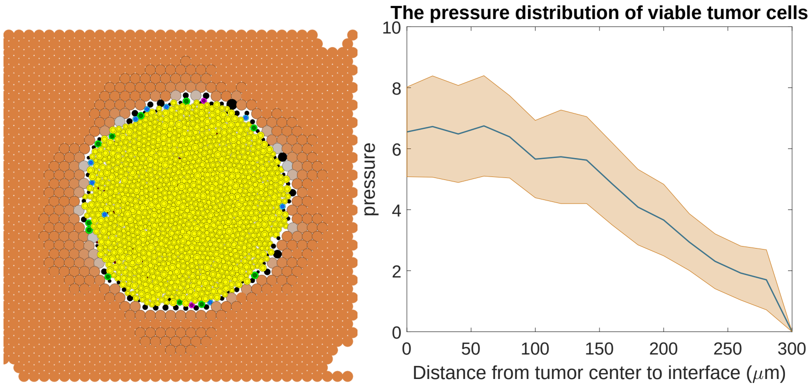


**Figure S4. Micrometastases without a mechanobiologic growth feedback.** *Left:* After 10 days of growth, almost all tumor cells have pressure exceeding 1.0, with considerable (non-physical) cell overlap. *Right:* We plot the mean (central curve) and standard deviation (shaded region) of the cell pressure versus distance from the tumor center of mass. Pressure peaks at $\sim 7$ nearest to the tumor center, gradually decreasing towards zero on the tumor edge where liver parenchyma apoptosis locally relieves the pressure. **Simulation parameters:** $r_E$=0.1 (1/min), $r_P$=0.01 (1/min) and $d_{max}$=1 $\mu$m.

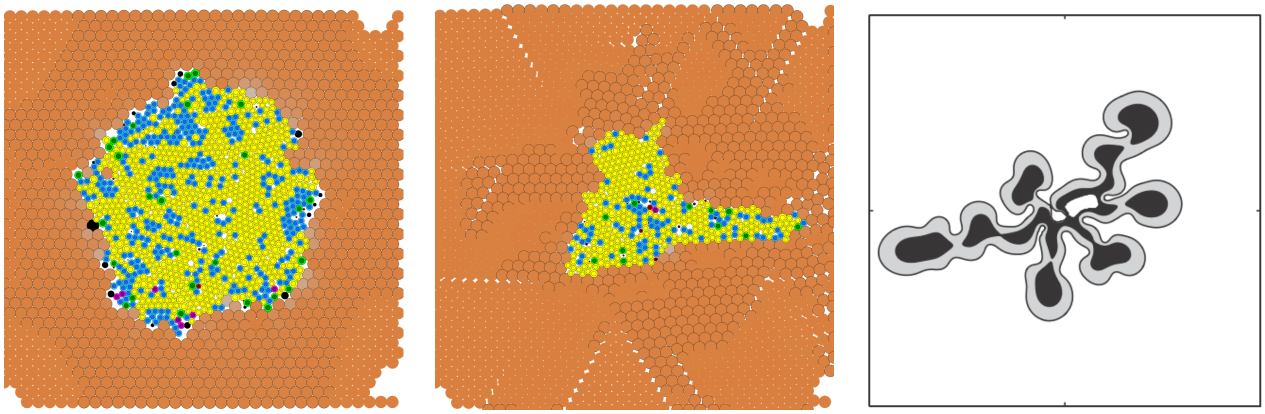

(a) $r_E$=0.1, $r_P$=0.01, $d_{max}$=1

(b) $r_E$=0.1, $r_P$=0.01, $d_{max}$=3

(c) continuum methods [1]

**Figure S5. Fingering metastatic growth: (a)-(b):** Growth into initially compliant parenchyma that tolerates deformation can lead to fingering morphologic instabilities, where tumor growth invades along directions of mechanical least resistance. Cells are colored as described in *Fig. 5* of main paper. **(c):** Macklin and Lowengrub previously used continuum methods to predict that tumor growth into relatively stiff, well-perfused tissue can lead to fingering-shaped morphological instabilities [1]. See *Fig. 8* of main paper for an example of CRC growth along paths of mechanical least resistance (yellow highlighted regions in the blue inset box).

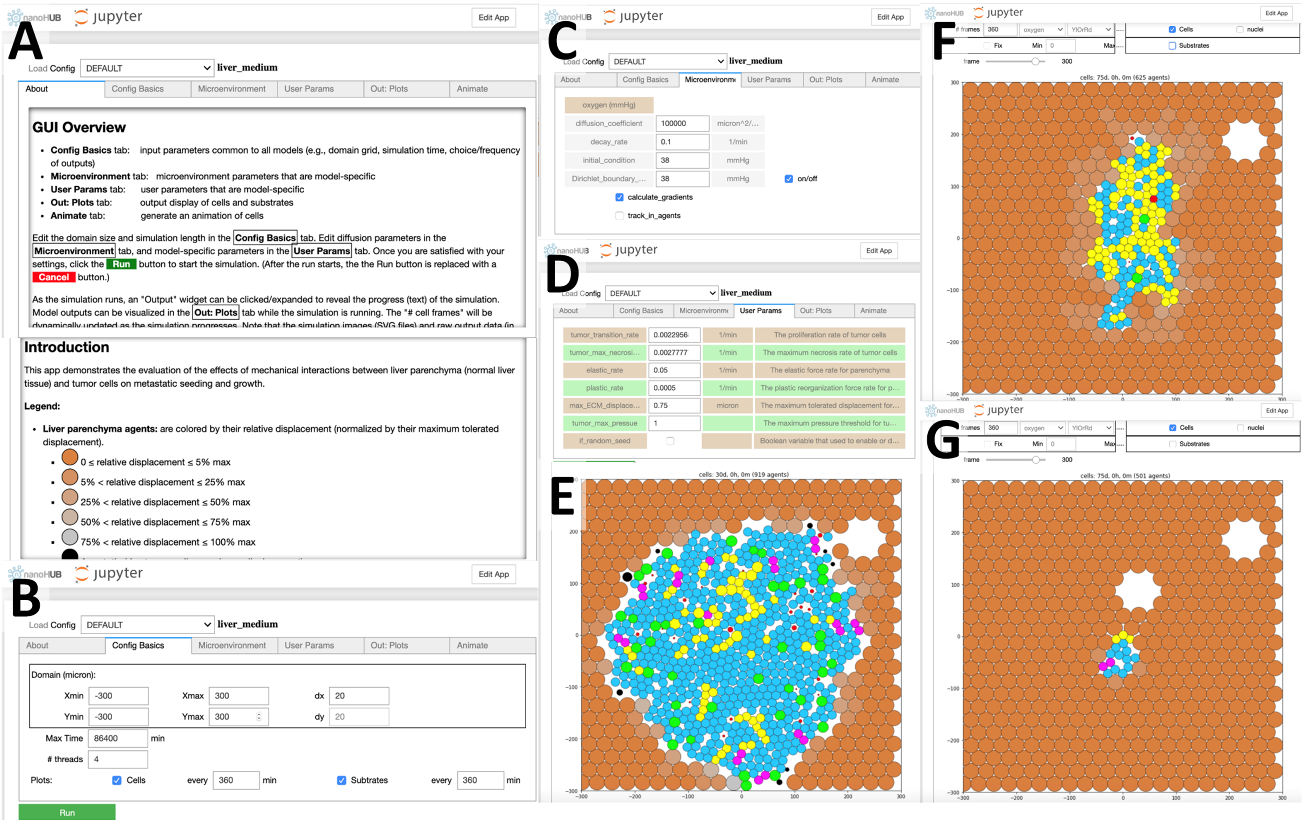

**Figure S6. Cloud-hosted interactive model:** Readers can interactively explore the agent-based model at https://nanohub.org/tools/pc4livermedium. (**A**) The *About* tab introduces the model, describes the plot legends, and gives run instructions. Users can modulate the domain size (**B**), diffusion parameters (**C**), and model-specific parameters (**D**). Simulation results can be viewed live in the *Out: Plots* tab, or animated as an mp4 movie in the *Animate* tab. Sample outputs are shown in **E**-**G**.

#### Supplementary Details on Results

##### Analysis of flow within micrometasases

Let us approximate fluid flow as identical to control conditions outside micrometastases and vanishingly small within tumors.

Using standard conservation of mass laws, we model the transport of a single growth substrate $\sigma$ (e.g., oxygen or glucose) in liver tissue as

$$\frac{\partial \sigma}{\partial t} = -\nabla \cdot \vec{J} - \lambda \sigma \quad (1)$$

$$\vec{J} = -D\nabla\sigma + \sigma\vec{u}, \quad (2)$$

where $\vec{u}$ is the fluid flow field (from the PVE model), $\lambda$ is the growth substrate consumption rate, $D$ is diffusion coefficient, and with boundary conditions to be described further below.

Based on the typical fluid velocities in unobstructed liver ($||\vec{u}|| \sim 10^{-4}$ m/sec) and tumor region ($||\vec{u}|| \sim 0$ m/sec), we find (after nondimensionalization) that growth substrate biotransport is largely advective in the parenchyma and diffusive within tumor regions. See Appendix for expanded detail.

##### An analytical approximation to oxygen distribution in unobstructed parenchyma

We can further build on these results to obtain an approximate analytical solution to the quasi-steady substrate distribution within an unobstructed lobule. Because the flow is largely advective-reactive, we can simplify the original equation to

$$\frac{\partial \sigma}{\partial t} = -\nabla \cdot (\sigma \vec{u}) - \lambda \sigma. \quad (3)$$

Next, assuming incompressible flow, approximating flow as radially symmetric ($r = 0$ corresponds to the central vein), and matching to the spatial lobule size from the PVE model results, we can obtain an approximate analytical solution for biotransport for oxygen ($\sigma$) throughout unobstructed lobules:

$$u(r) = 6 \times 10^4 e^{-0.0075r} \quad (4)$$

$$\sigma(r) = 38.9 e^{0.0223\left(e^{0.0075r}-1\right)}. \quad (5)$$

See Appendix for expanded mathematical details.

We plot the radial flow and oxygenation profiles for these parameters in a hepatic lobule with radius $R = 400\,\mu$m in Fig. S7; note that $\sigma(R) \approx 60$ mmHg, comparable to the portal venule oxygenation reported in [2].

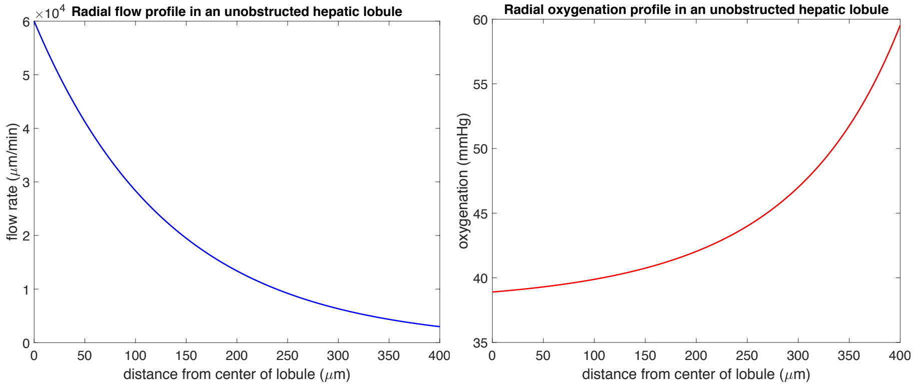


**Figure S7.** Approximate analytical solution to flow (**a**) and oxygen distribution (**b**) in an individual, unobstructed lobule

##### Approximate oxygen distribution in large tissue sections

We can extend the analytical single-lobule approximation to approximate advection-dominated perfusion in a large tissue section. We generated a large 1 cm$^2$ section of liver tissue with hundreds of hepatic lobules with central veins at positions $\{\vec{x}_i\}_{i=1}^n$. To approximate the quasi-steady oxygenation p$O_2$ at any position $\vec{x}$ of unobstructed tissue, let us define

$$r(\vec{x}) = \min\{||\vec{x}_i - \vec{x}||\}_{i=1}^n \quad (6)$$

to be the distance between $\vec{x}$ and the nearest central vein. Then we approximate

$$\mathrm{pO}_2(\vec{x}) = \sigma\left(r(\vec{x})\right). \quad (7)$$

Here, $\sigma$ is the radial oxygenation profile defined in Equation 5.

Note that this is equivalent to partitioning the 2-D tissue cross-section into a Voronoi mesh (with portal triads at the vertices (representing the flow source) and central veins in the centers) and assuming radial flow within each Voronoi polygon. We show this approximation for the 1 $cm^2$ tisue in Fig. S8.

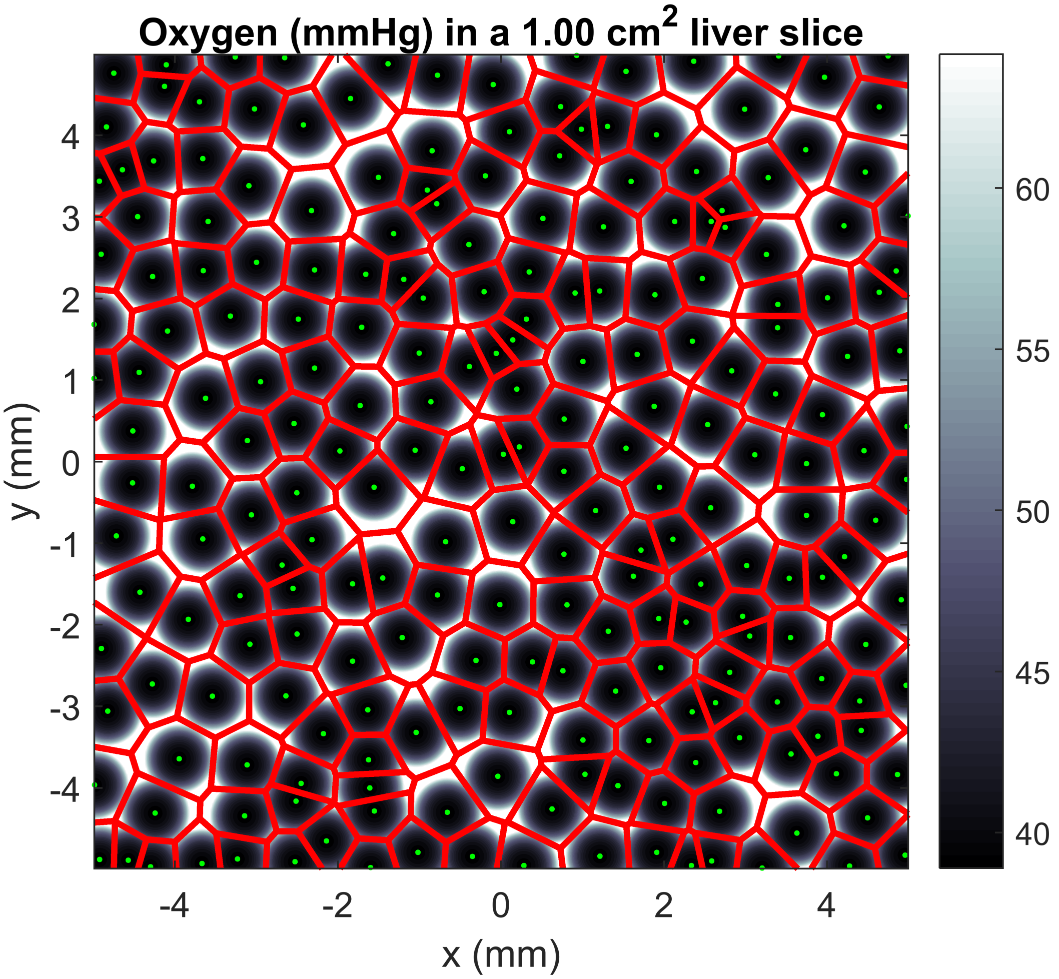


**Figure S8.** Approximate oxygen distribution (greyscale values) and Voronoi lobule boundaries (red lines) in a 1 $cm^2$ liver tissue.

### Appendix: Mathematical modeling of liver biology and metastases, and cancer mechanobiology

#### Liver regeneration

Hoehme et al. used a closed source agent-based modeling (ABM) framework similar to PhysiCell [3] to model liver damage and evaluate the cell-scale hypotheses needed for successful regeneration of the microarchitecture [4]. The model successfully replicated all hepatocytes and sinusoids within a liver lobule during regeneration [4]. More recently, they used a deformable cell-based model to explore how liver lobule cells regenerate with mechanical stress effects [5], finding that the migration forces that a cell needs to exert on its environment are much smaller than those predicted by center-based models that do not explicitly model cell morphologies. (Center-based models are off-lattice methods that track each cell's center of mass and size, typically using one agent to represent one cell [6]).

#### Liver toxicology

Schwen et al. described a methodology of exploring whole-body pharmacokinetics models based on a liver model, integrating from cells through sinusoids and the organ to the whole organism [7]. Sluka et al. developed a multi-scale, liver-centric physiologic-based pharmacokinetics (PBPK) model to study acetaminophen pharmacology and metabolism [8]. They explored drug uptake and transport at the whole body level, blood flow at the tissue/organ level, and metabolism at the sub-cellular level, showing the model is capable of predicting liver toxicity from acetaminophen overdose.

#### Interstitial flow and mechanics in liver tissues

Rani et al. presented a computational fluid dynamics (CFD) model of non-Newtonian blood flow within a portal vein, hepatic artery and two sinusoids [9]. Ricken et al. developed a two-scale, continuum multicomponent model for studying blood perfusion and cell metabolism within the liver [10]. They used porous media theory to explore perfusion within lobules, with a focus on advective and diffusive solute transport. Lettmann et al. used a porous-medium method to simulate the microcirculation fields of liver lobules [11]. They captured the blood flow velocity inside the sinusoids and oxygen consumption of Rappaport's liver acinus model. The work found that the liver microcirculation plays an important role in establishing inflammatory cytokine levels.

White et al. presented a convection-diffusion-reaction partial differential equation (PDE) model that coupled blood flow, drug distribution, and clearance in a 3-D liver [12]. Siggers et al. presented a mathematical model that considers the sinusoidal and interstitial spaces as porous media to simulate the blood, interstitial flow and lymph production in the liver [13]. Meyer et al. developed a 3D multi-scale model that explored the biliary fluid dynamics in the mouse liver lobule, predicting gradients of bile velocity and pressure within the lobule [14]. They found that osmotic effects and bile canaliculi contractility result in the bile flow. The multi-scale model is capable of predicting the biliary distribution on drug-induced liver injury. Nishii et al. developed sophisticated poro-viscoelastic (PVE) models that extend porous flow models (as described above) by coupling the flow to a moving, viscoelastic tissue [15]. We shall describe this model further below.

#### Tissue mechanics in tumor growth

Ambrosi et al. developed an elasto-visco-plastic model that used phase field methods to describe the mechanical stress of tumor cell suspensions and aggregates [16, 17]. The model illustrates tumor growth responses to mechanical stress, cellular release of stresses, the formation of tumor capsules, and compression of neighboring tissues. Antonio et al. introduced an agent-based model for studying elasto-plastic mechanical interactions between cell-basement membrane (BM)-extracellular matrix (ECM) [18]. Cell motion was modeled as resulting from a balance between adhesive, elastic, and repulsive forces. Over short times, elastic forces were used to simulate the interaction between BM and ECM; over long times, the BM underwent plastic relaxation in response to prolonged strains. Liedekerke et al. used an agent-based model to study the tumor spheroids' growth dynamics in mechanical stress environment with calibrated parameters from experiments [19]. They discovered that the growth response of cell populations on external mechanical stress is predicable with a same growth progression function.

#### Liver metastases in connection with ECM remodeling

Mathematical modeling has also been used to study liver fibrosis [20, 21], while others have used PDE methods to model the growth of liver metastases [22–24]. The kinetics of integrin receptor binding to hepatic ECM proteins following liver injury were recently modeled by Hudson et al. [25]. In follow-on work, the growth of liver metastases was evaluated in a tissue-scale model [26], taking into consideration heterogeneous macrophage populations [27]. ECM from both injured and normal liver was modeled, and parameters were set with values from mouse *in vivo* experiments designed to mimic human tissue conditions.

#### Mechanosensing and cancer cell proliferation

Others have evaluated the impact of stress on tumor growth and invasion. Voutouri et al. used both in *vivo* experiment and continuum modeling to study the effect of tumor-host mechanical interactions on the growth of solid tumors [28]. The results show that solid stresses can inhibit tumor cell proliferation, induce apoptosis in surrounding tissue, and lead the surrounding tissue to resist tumor expansion. Cheng et al. [29] built an experimental model to explore spatial distribution of compressive stress around tumor spheroids representing avascular nodules, showing that high mechanical stress results in suppression of tumor cell proliferation and apoptotic cell death.

##### Metastasis

While there has been substantial work in modeling primary tumor growth in detailed tissues [6, 30], to our knowledge there has been little work in spatial models of early metastatic seeding and growth under mechanical and flow constraints, especially in liver tissue. Macklin and Lowengrub used a level set / moving boundary method to model tumor growth in simulated brain tissue in [31], but they did not consider mechanobiologic feedback or the impact of flow. Their continuum approach was also not suited to tumors with low cell-cell adhesion or high cellular heterogeneity. Baratchart et al. recently developed a similar model (also based on a porous flow formulation, but without the level set-based sharp interface model) to simulate the merger of renal metastatic foci [32], with a very simple pressure-based tumor cell proliferation rate, and without considering any mechanobiologic feedback in the surrounding normal tissue.

Others have developed network models of metastatic seeding [33–38], but without spatially-resolved simulation of the individual metastasis sites, nor of any mechanical effects. Treatment of liver metastases via nanotherapy taking into account immune system interactions have been recently modeled [39–41], but without considering mechanical effects. In particular, detection and drug delivery to colorectal cancer liver metastases via nanoparticles were simulated in a tissue-scale model in [42, 43]. See [44] for further excellent materials on experimental and mathematical modeling of metastasis.

#### Appendix:
#### Details on dimensional analysis of PVE model

To evaluate the relative contribution of the advective and diffusive effects in liver interstitial fluid flow, we can rewrite Equations 1-2 as

$$\frac{\partial \sigma}{\partial t} = D\nabla^2\sigma - \vec{u}\cdot\nabla\sigma - \sigma\nabla\cdot\vec{u} - \lambda\sigma. \tag{8}$$

Approximating the flow field as slowly varying ($\vec{u} \approx u\vec{w}$ for some constant $u$ and a constant unit vector $\vec{w}$), $\nabla\cdot\vec{u} \approx 0$. If we nondimensionalize space with scale $L$ and time with scale $\bar{t}$, then this becomes

$$\frac{\partial \sigma}{\partial t} = \left(\frac{D\bar{t}}{L^2}\right)\nabla^2\sigma - \left(\frac{u\bar{t}}{L}\right)\vec{w}\cdot\nabla\sigma - \left(\lambda\bar{t}\right)\sigma. \tag{9}$$

Choosing

$$\frac{u\bar{t}}{L} = 1 \text{ and } \lambda\bar{t} = 1 \Longrightarrow \bar{t} = \frac{1}{\lambda} \text{ and } L = \frac{u}{\lambda}, \tag{10}$$

then the nondimensionalized equation becomes

$$\frac{\partial \sigma}{\partial t} = \left(\frac{D\lambda}{u^2}\right)\nabla^2\sigma - \vec{w}\cdot\nabla\sigma - \sigma. \tag{11}$$

By prior work [45], $D \sim 10^5 \mu\mathrm{m}^2/\mathrm{min}$ and $\lambda \sim 10\ \mathrm{min}^{-1}$. By our PVE simulations of a control case (with no tumor cells) and with small micrometastases, as well as the simulations in [15], $u \sim 10^{-4}\mathrm{m/sec} = 6\times10^3 \mu\mathrm{m/min}$ in the unobstructed liver parenchyma, and so $L \sim 600\,\mu$m, and $\bar{t} \sim 6$ sec. The diffusion term has relative order of magnitude

$$\frac{D\lambda}{u^2} \sim \frac{1}{36} \approx 0.03. \tag{12}$$

Thus, in unobstructed regions of liver lobule, flow is primarily advective-reactive, and the original equation can be simplified to

$$\frac{\partial \sigma}{\partial t} = -\nabla\cdot(\sigma\vec{u}) - \lambda\sigma. \tag{13}$$

Because the time scale is on the order of seconds, advective flow is at quasi-steady state in regions of unobstructed flow when considered on time scales of mechanics (minutes) and tumor growth (hours to days).

Because $\vec{u} \sim \mathbf{0}$ inside micrometastases, we find that biotransport in those regions satisfies

$$\frac{\partial \sigma}{\partial t} \approx D\nabla^2\sigma - \lambda\sigma. \tag{14}$$

That is, biotransport is largely diffusive-reactive within tumors.

### Appendix:
### Details on radial biotransport approximation

Here, we give expanded mathematical detail on the approximate analytical solutions for oxygen distribution in unobstructed liver tissue.

Assuming incompressible flow ($\nabla \cdot \vec{u} = 0$) and that flow within the lobule can be approximated as radially symmetric ($\vec{u} \approx -u(r)\hat{\vec{r}}$, where $r$ is the distance from the middle of the central vein), we have

$$\frac{\partial \sigma}{\partial t} = u(r)\nabla \sigma \cdot \hat{\vec{r}} - \lambda \sigma \tag{15}$$

$$= u(r)\frac{\partial \sigma}{\partial r} - \lambda \sigma. \tag{16}$$

Under quasi-steady conditions, $\sigma = \sigma(r)$, and we have

$$0 = \sigma'(r) - \frac{\lambda}{u(r)}\sigma, \tag{17}$$

whose analytical solution is given by

$$\sigma(r) = ce^{\lambda \int_0^r \frac{1}{u(s)} ds} \tag{18}$$

for some constant $c$.

For simplicity, we will approximate the flow profiles shown for no tumor cells or with small micrometastases with the form

$$u(r) \approx ae^{-br} \tag{19}$$

for constants $a$ and $b$. Notice that $u(0) = a$ and $u(R) = ae^{-bR} = u(0)e^{-bR}$, whereby

$$b = -\frac{\ln\left(\frac{u(R)}{u(0)}\right)}{R}. \tag{20}$$

For consistency with the PVE model, we set $R = 400\,\mu$m.

For this simplified form, we have

$$\sigma(r) \approx ce^{\frac{\lambda}{ab}\left(e^{br}-1\right)}. \tag{21}$$

By our PVE simulations for the control with no tumor cells or with small micrometastases, we see that $\frac{u(R)}{u(0)}$ is on the order of 0.01 to 0.1, so we choose 0.05. Thus, $b \sim 0.0075\,\mu\text{m}^{-1}$ by Equation 20. Moreover, $u(0) = a \sim 10^{-3}$ m/sec $= 6 \times 10^4\,\mu$m/min. [2] report that liver oxygenation is approximately 38.9 mmHg in central venules, 48.2 mmHg in sinusoids, and 59.8 mmHg in portal venules. So, we set $\sigma(0) = 38.9$ mmHg:

### Appendix:
### Code availability

#### .1 Poroviscoelastic model

The project files for the poroviscoelastic simulation work are available as open source under the MIT license at

https://github.com/MathCancer/PVE-liver-mets

##### Zenodo archives

**1.0.0** Version used for the PVE simulation studies
https://doi.org/10.5281/zenodo.3779512

#### Agent-based model

The core agent-based model (minus the cloud-hosted interface) is available as open source under the 3-clause BSD license at

https://github.com/MathCancer/liver-mechanobiology

##### Zenodo archives

**1.0.0** Version used for the main parameter studies
https://doi.org/10.5281/zenodo.3766870

**1.1.0** Update to allow resizeable domains
https://doi.org/10.5281/zenodo.3766979

#### nanoHUB cloud-hosted model

The cloud-hosted deployment of the agent-based model can be accessed and run for free at

https://nanohub.org/tools/pc4livermedium.

The source code is available as open source under the 3-clause BSD license at

https://github.com/yafeiwang89/liver_medium

##### Zenodo archives

**1.3** Source code for nanoHUB app version 1.3
http://doi.org/10.5281/zenodo.3775924

#### Supplementary Materials for PVE MODEL:

### Impact of tumor-parenchyma biomechanics on liver metastatic progression: a multi-model approach


Yafei Wang[†], Erik Brodin[†], Kenichiro Nishii, Hermann B Frieboes, Shannon Mumenthaler, Jessica L. Sparks[1], Paul Macklin[2]

[†]Contributed equally to this work

### 1. Material Properties

Model material properties are given in Tables S1-S3. The Young's modulus of the parenchyme was taken as 4.4 kPa and Poisson's ratio as 0.35 (Evans et al. 2013; Nishii et al. 2016). The Young's modulus for tumor tissue was set as 30 kPa (Venkatesh et al. 2008; Lu et al. 2015) and Poisson's ratio was assumed equal to normal tissue. A four-term Prony series expansion was used to model tissue viscoelasticity, with Prony series constants taken from previous nano-indentation experiments on perfused liver tissue (Evans et al. 2013).

At the macroscopic (cm) length scale, tumors have been reported to have higher hydraulic conductivity *K* relative to the surrounding tissue (Swabb et al. 1974; Netti et al. 2000; Pishko et al. 2011). However, the hydraulic conductivity of smaller tumors on the order of 50-400 μm is unknown. For each tumor size/location combination in the present study, we simulated the effects of high and low hydraulic conductivity in the tumor. For the high hydraulic conductivity case, we used a value of 3.65 times that of normal liver, based on hydraulic conductivity of hepatocarcinoma determined from measurements of (Swabb et al. 1974). For the low hydraulic conductivity case, we used 0.3 times that of normal liver, to span approximately one order of magnitude in our high/low conditions. Normal liver parenchyme hydraulic conductivity was taken as $1.85 \times 10^{-6}$ m/s as reported by (Nishii et al. 2016), based on void ratio measurements from histological image analysis. Void ratio, e, is defined as:

$$e = \frac{V_v}{V_s} = \frac{V_v}{V_t - V_v} \qquad \text{(Eq. S1)}$$

where $e$ is the void ratio, $V_v$ is the void volume, $V_s$ is the volume of solid in the lobule, and $V_t$ is the total volume of the lobule. Void ratio is related to hydraulic conductivity as follows (Nishii et al. 2016):

$$K = \frac{\rho g e r^2}{8\mu(1+e)} \qquad \text{(Eq. S2)}$$

where $K$ is the hydraulic conductivity (m/s), $\rho$ is fluid density (kg/m$^3$), $g$ is gravitational acceleration (m/s$^2$), $e$ is the void ratio, $r$ is the average radius of a sinusoid (m), and $\mu$ is the dynamic viscosity of the permeating fluid (Pa·s).

**Table S1. Elastic constants for parenchyme and tumor. Materials are assumed isotropic.**

| | Parenchyme | Tumor |
|---|---|---|
| Elastic Modulus (kPa) | 4.4 | 30 |
| Poisson's ratio | 0.35 | 0.35 |

**Table S2. Viscoelastic constants (same values used for parenchyme and tumor).**

| Term | $g_i$ (dim) | $\tau_i$ (sec) |
|---|---|---|
| 1 | 0.53 | 2.00E-05 |
| 2 | 0.376 | 1 |
| 3 | 0.027 | 7.65 |
| 4 | 0.01 | 100 |

**Table S3. Hydraulic conductivity and related properties**

| Property | Parenchyme | Tumor, High Hydraulic Conductivity | Tumor, Low Hydraulic Conductivity |
|---|---|---|---|
| Hydraulic Conductivity (m/s) | 1.85E-06 | 6.75E-06 | 5.50E-07 |
| Void Ratio | 0.8 | 1.55 | 0.052 |
| Specific Weight of Fluid ($kg/m^3$) | 9855 | 9855 | 9855 |

#### 2. Boundary Conditions

As noted in the manuscript, one-quarter of the hexagonal lobule was modeled to minimize computational cost; therefore symmetric boundary conditions were applied to the *X*, *Y*, and *Z* planes (see Model Geometry). Following the work of (Bonfiglio et al. 2010), the model assumed uniform expansion in the axial direction (in the *XY* plane) and no expansion in the longitudinal (*Z*) direction. Pressure in the CV surface was set to zero to serve as a pressure sink, and pressures in the terminal portal vein (tPV) and pre-terminal portal vein (pre-tPV) were set to 2.23 mmHg (297.3 Pa) above the CV pressure so that the physiological pressure difference was preserved (Nishii et al. 2016). The pressure condition at the tumor boundary varied depending on tumor size (Table S4). A total of 13 model runs were performed, varying tumor position and size, pressure condition at the tumor boundary, and tumor permeability relative to surrounding parenchyme (Table S4). Lastly, a control model was created with no tumor but with identical parenchyme properties and vascular pressure boundary conditions, to serve as a basis for comparison.

**Table S4. Variable parameters for finite element model runs.**

| Model Run | Tumor Position | Tumor Size (diam., μm) | Tumor as Pressure Source or Sink (Pa) | Tumor Hydraulic Conductivity (m/s) |
|---|---|---|---|---|
| 1 | Center | Seed (50) | Neutral | Low (5.50E-07) |
| 2 | Center | Seed (50) | Neutral | High (6.75E-06) |
| 3 | Center | Small (200) | Sink (0) | Low (5.50E-07) |
| 4 | Center | Small (200) | Sink (0) | High (6.75E-06) |
| 5 | Portal | Seed (50) | Neutral | Low (5.50E-07) |
| 6 | Portal | Seed (50) | Neutral | High (6.75E-06) |
| 7 | Portal | Small (200) | Sink (0) | Low (5.50E-07) |
| 8 | Portal | Small (200) | Sink (0) | High (6.75E-06) |
| 9 | Portal | Medium (400) | Source (600) | Low (5.50E-07) |
| 10 | Portal | Medium (400) | Source (600) | High (6.75E-06) |
| 11 | Portal | Medium (400) | Neutral | Low (5.50E-07) |
| 12 | Portal | Medium (400) | Neutral | High (6.75E-06) |
| 13 | Control | No Tumor | N/A | Parenchyme (1.85E-06) |

### 3. Fluid Velocity, Strain, Stress, and Pore Fluid Pressure Results

(A)

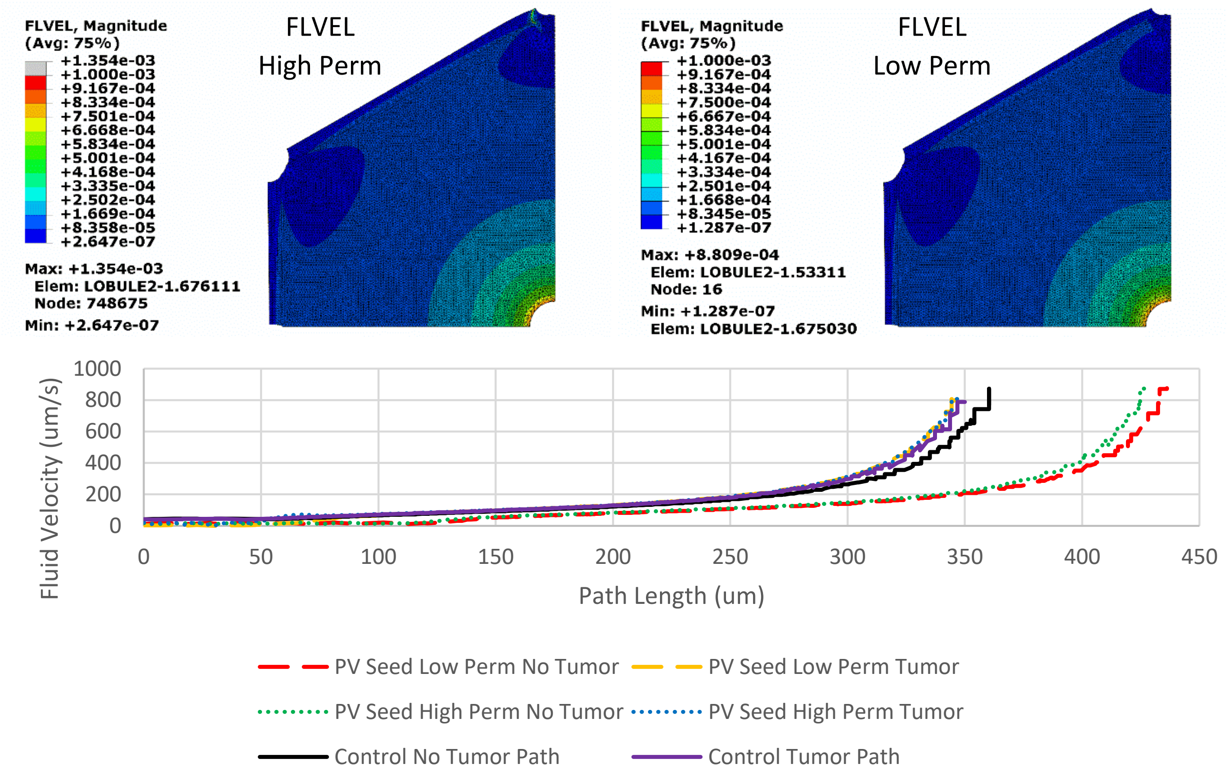


(B)

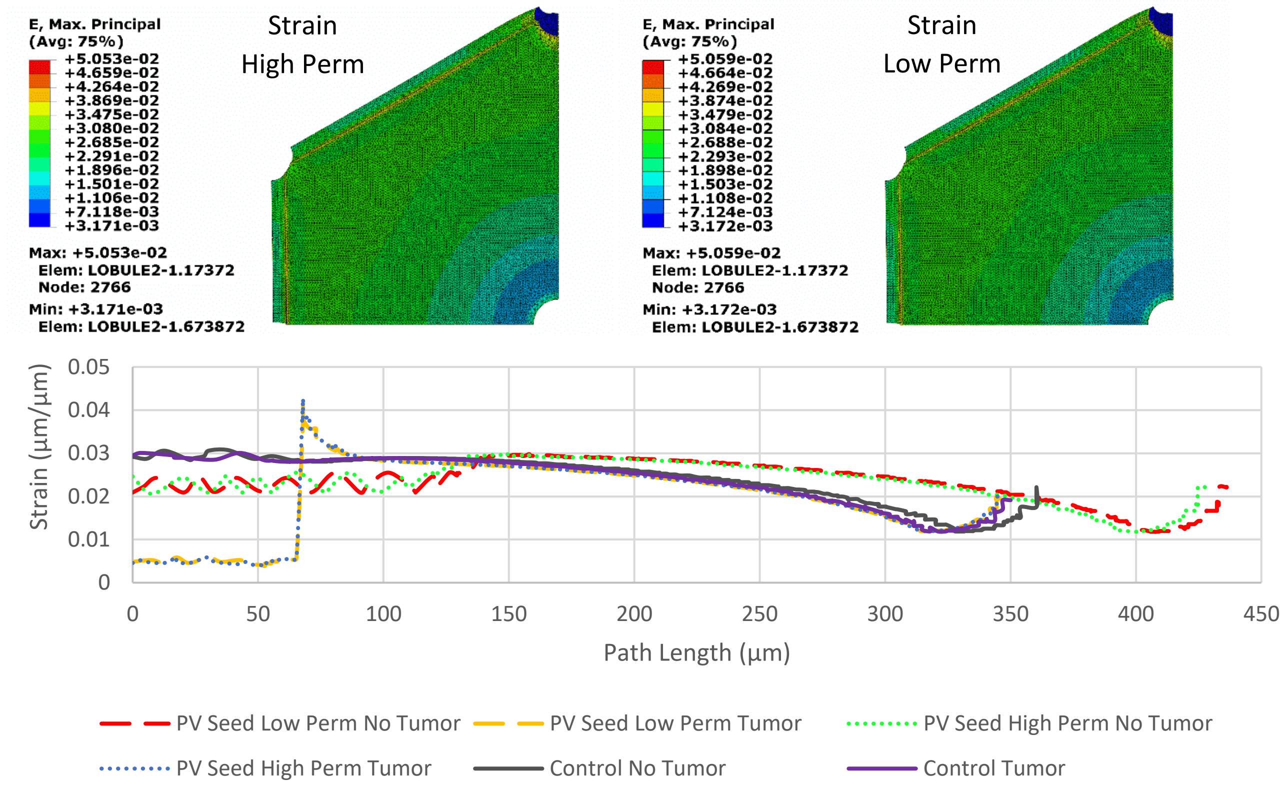

(C)

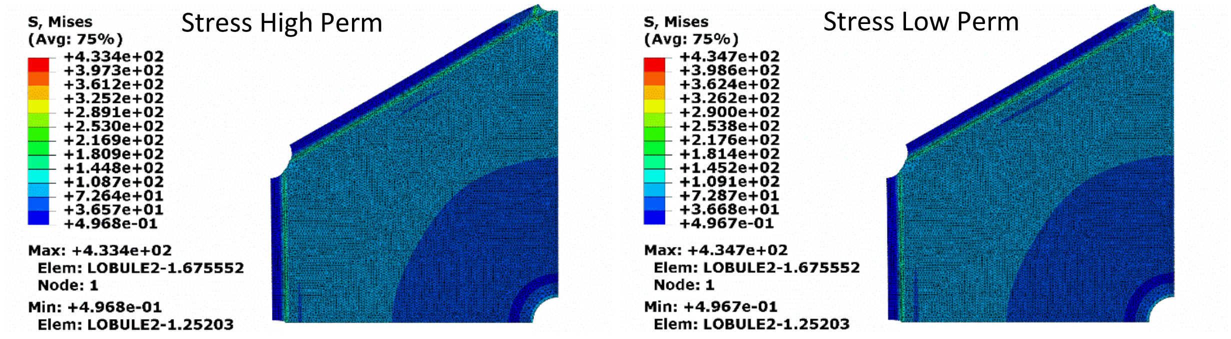


(D)

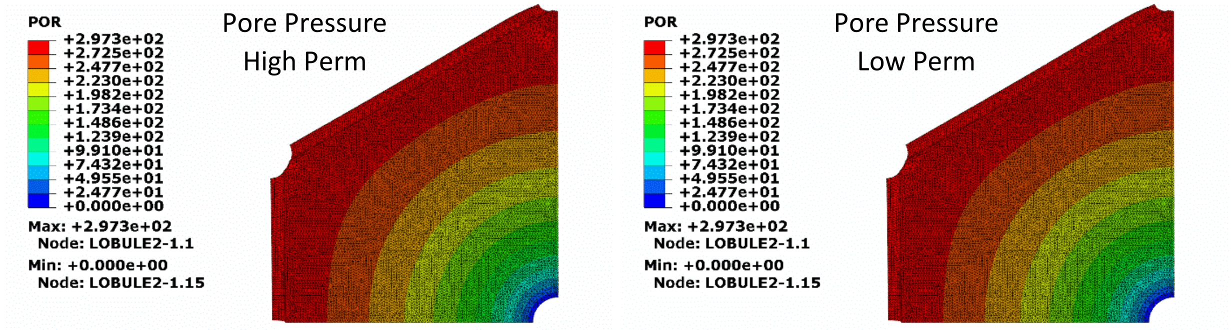


**Figure S9.** Images for the 50µm diameter tumor seed centered at the portal vein. Tumor was not set as a pressure source or sink. Images for (A) fluid velocity, (B) strain, (C) stress, and (D) pore pressure distribution throughout the quarter lobule. Graphs represent the fluid velocity and strain in the “tumor” and “no tumor” paths.

(A)

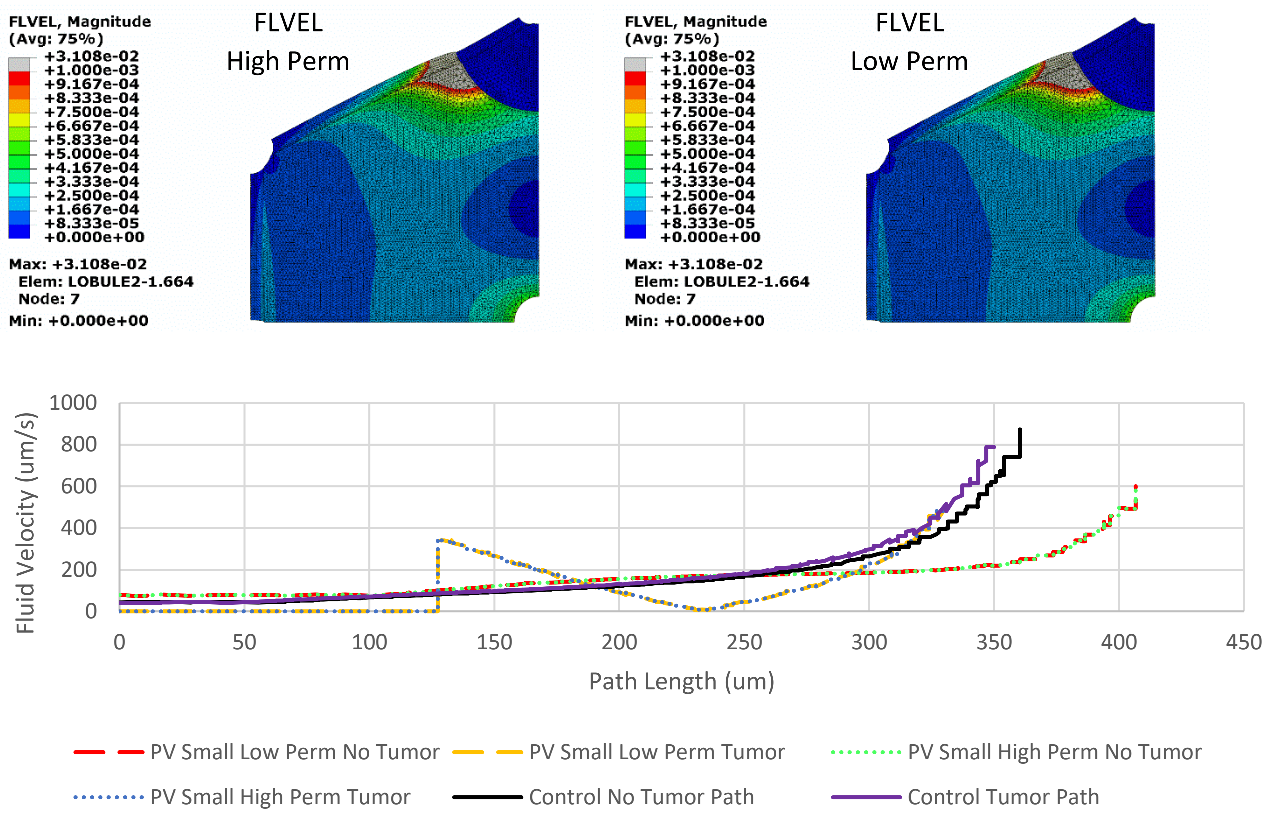
FLVEL, Magnitude
(Avg: 75%)
+3.108e-02
+1.000e-03
+9.167e-04
+8.333e-04
+7.500e-04
+6.667e-04
+5.833e-04
+5.000e-04
+4.167e-04
+3.333e-04
+2.500e-04
+1.667e-04
+8.333e-05
+0.000e+00
Max: +3.108e-02
Elem: LOBULE2-1.664
Node: 7
Min: +0.000e+00
FLVEL
High Perm
FLVEL
Low Perm
Fluid Velocity (um/s)
Path Length (um)
PV Small Low Perm No Tumor
PV Small Low Perm Tumor
PV Small High Perm No Tumor
PV Small High Perm Tumor
Control No Tumor Path
Control Tumor Path

(B)

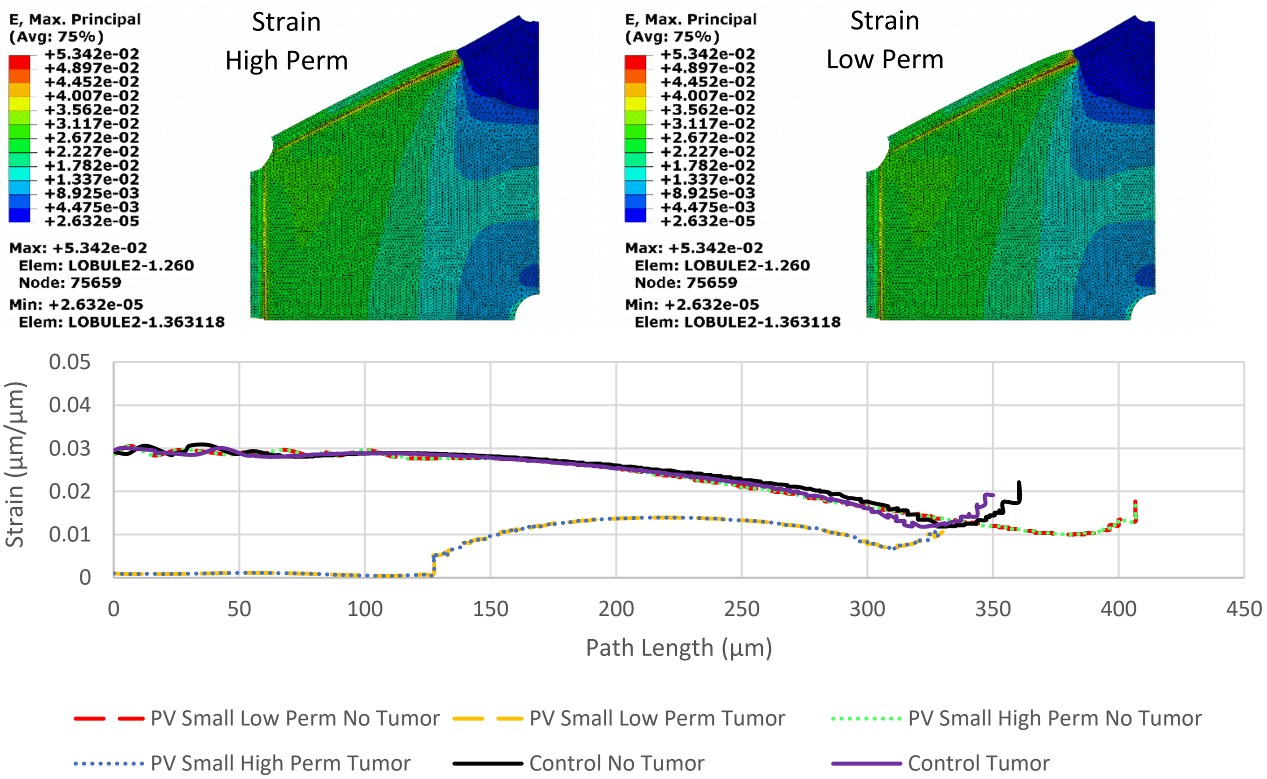
E, Max. Principal
(Avg: 75%)
+5.342e-02
+4.897e-02
+4.452e-02
+4.007e-02
+3.562e-02
+3.117e-02
+2.672e-02
+2.227e-02
+1.782e-02
+1.337e-02
+8.925e-03
+4.475e-03
+2.632e-05
Max: +5.342e-02
Elem: LOBULE2-1.260
Node: 75659
Min: +2.632e-05
Elem: LOBULE2-1.363118
Strain
High Perm
Strain
Low Perm
Strain (µm/µm)
Path Length (µm)
PV Small Low Perm No Tumor
PV Small Low Perm Tumor
PV Small High Perm No Tumor
PV Small High Perm Tumor
Control No Tumor
Control Tumor

(C)

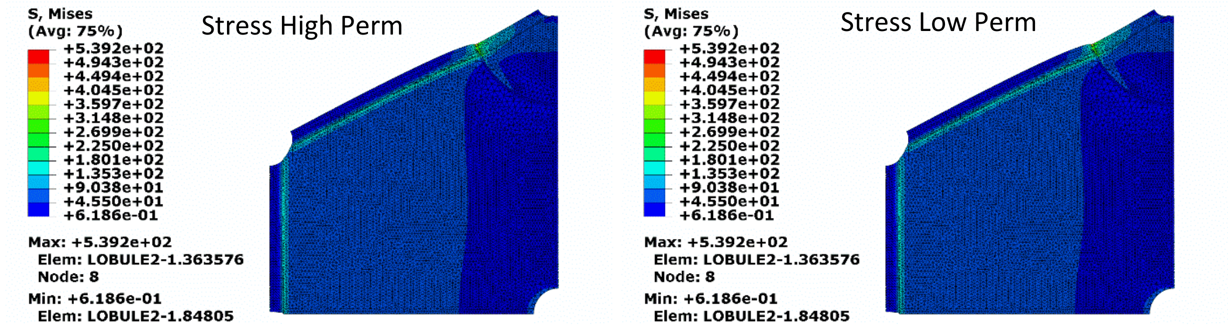


(D)

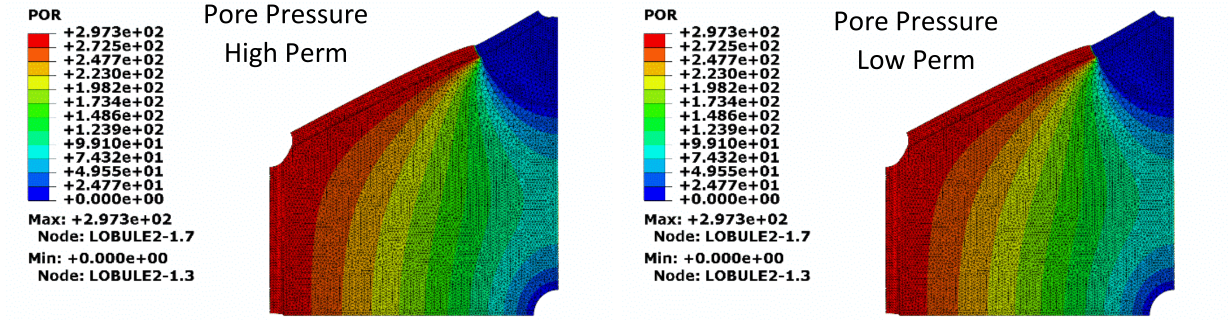


**Figure S10.** Images for the 200μm diameter tumor centered at the portal vein. Tumor was set as a pressure sink. Images for (A) fluid velocity, (B) strain, (B) stress, and (D) pore pressure distribution throughout the quarter lobule. Graphs represent the fluid velocity and strain in the “tumor” and “no tumor” paths.

(A)

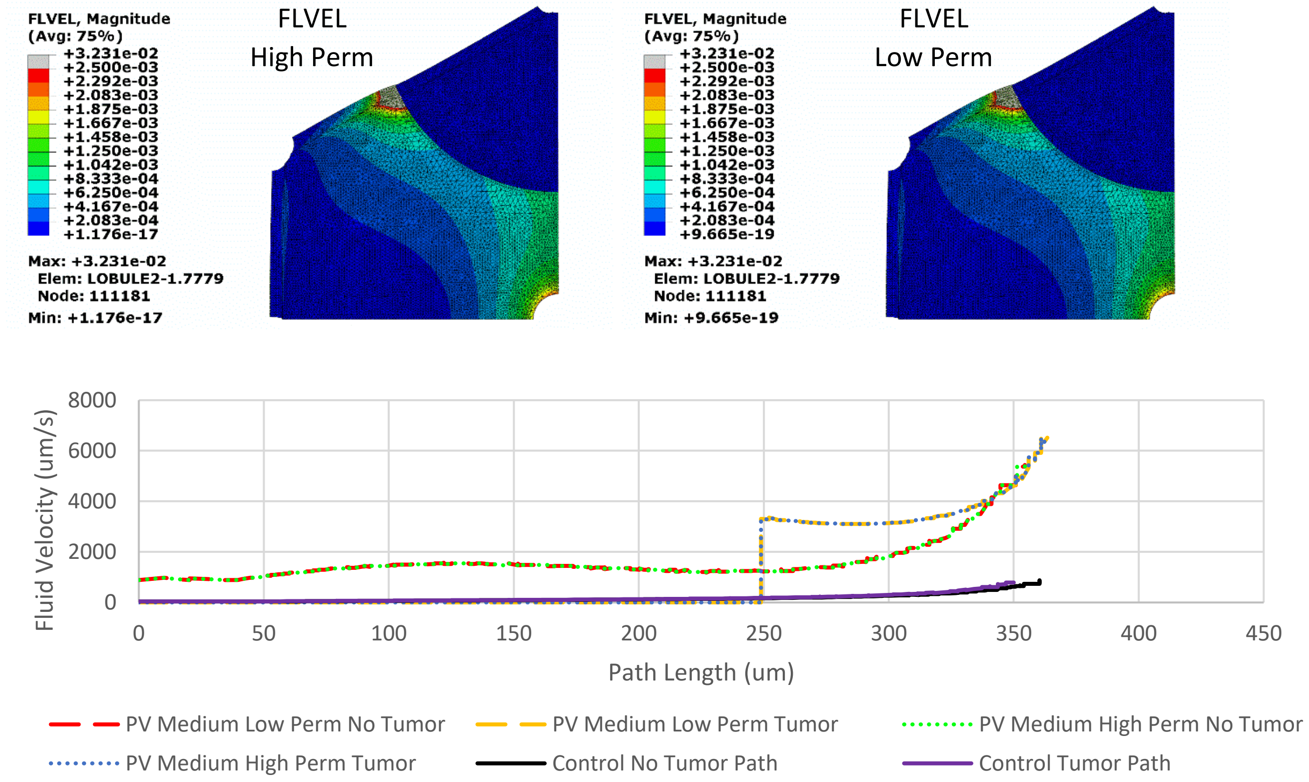

FLVEL, Magnitude
(Avg: 75%)
+3.231e-02
+2.500e-03
+2.292e-03
+2.083e-03
+1.875e-03
+1.667e-03
+1.458e-03
+1.250e-03
+1.042e-03
+8.333e-04
+6.250e-04
+4.167e-04
+2.083e-04
+1.176e-17
Max: +3.231e-02
Elem: LOBULE2-1.7779
Node: 111181
Min: +1.176e-17
FLVEL
High Perm
FLVEL, Magnitude
(Avg: 75%)
+3.231e-02
+2.500e-03
+2.292e-03
+2.083e-03
+1.875e-03
+1.667e-03
+1.458e-03
+1.250e-03
+1.042e-03
+8.333e-04
+6.250e-04
+4.167e-04
+2.083e-04
+9.665e-19
Max: +3.231e-02
Elem: LOBULE2-1.7779
Node: 111181
Min: +9.665e-19
FLVEL
Low Perm
Fluid Velocity (um/s)
8000
6000
4000
2000
0
0
50
100
150
200
250
300
350
400
450
Path Length (um)
PV Medium Low Perm No Tumor
PV Medium Low Perm Tumor
PV Medium High Perm No Tumor
PV Medium High Perm Tumor
Control No Tumor Path
Control Tumor Path


(B)

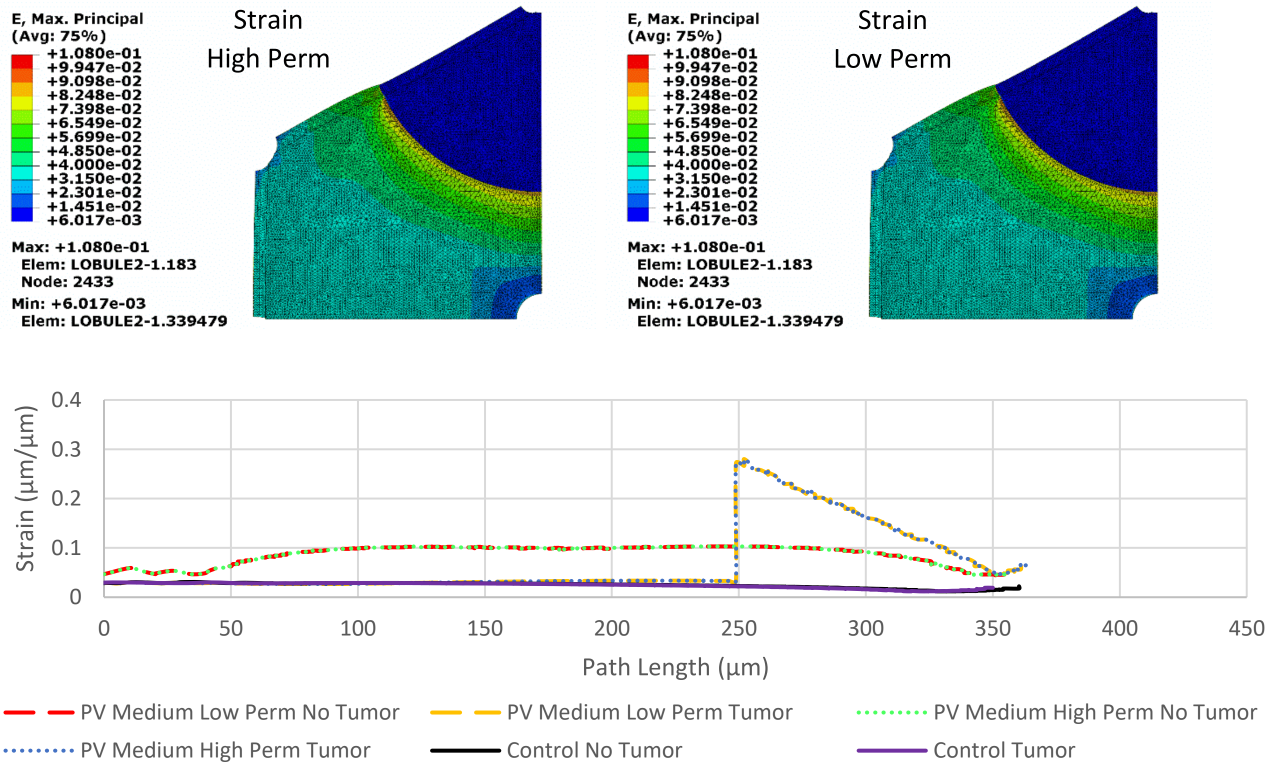

E, Max. Principal
(Avg: 75%)
+1.080e-01
+9.947e-02
+9.098e-02
+8.248e-02
+7.398e-02
+6.549e-02
+5.699e-02
+4.850e-02
+4.000e-02
+3.150e-02
+2.301e-02
+1.451e-02
+6.017e-03
Max: +1.080e-01
Elem: LOBULE2-1.183
Node: 2433
Min: +6.017e-03
Elem: LOBULE2-1.339479
Strain
High Perm
E, Max. Principal
(Avg: 75%)
+1.080e-01
+9.947e-02
+9.098e-02
+8.248e-02
+7.398e-02
+6.549e-02
+5.699e-02
+4.850e-02
+4.000e-02
+3.150e-02
+2.301e-02
+1.451e-02
+6.017e-03
Max: +1.080e-01
Elem: LOBULE2-1.183
Node: 2433
Min: +6.017e-03
Elem: LOBULE2-1.339479
Strain
Low Perm
Strain (µm/µm)
0.4
0.3
0.2
0.1
0
0
50
100
150
200
250
300
350
400
450
Path Length (µm)
PV Medium Low Perm No Tumor
PV Medium Low Perm Tumor
PV Medium High Perm No Tumor
PV Medium High Perm Tumor
Control No Tumor
Control Tumor

(C)

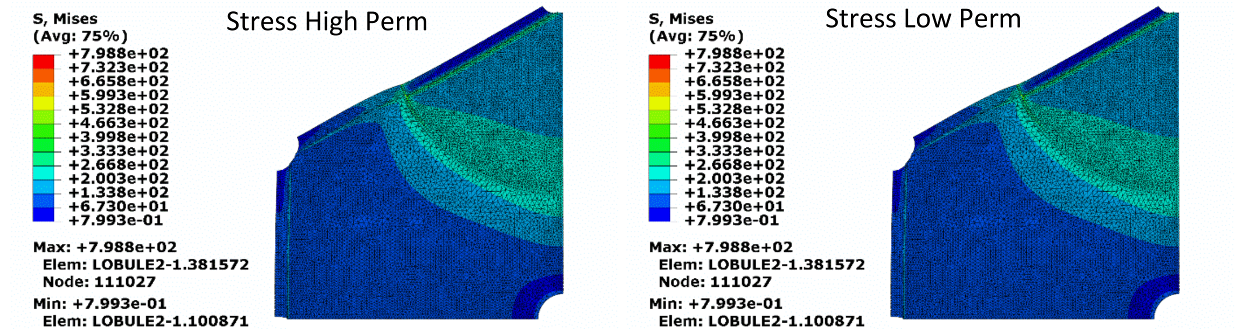


(D)

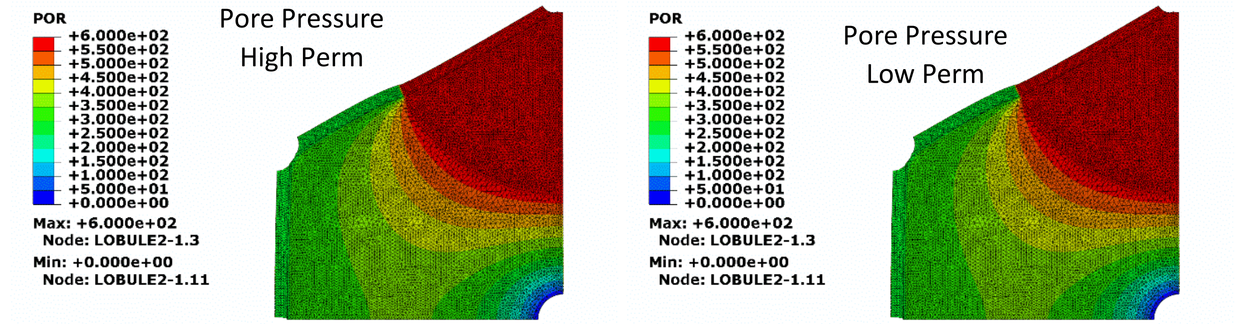


**Figure S11.** Images for the 400µm diameter tumor centered at the portal vein. Tumor was set as a 600 Pa pressure source. Images for (A) fluid velocity, (B) strain, (C) stress, and (D) pore pressure distribution throughout the quarter lobule. Graphs represent the fluid velocity and strain in the “tumor” and “no tumor” paths. Note the scale of fluid velocity and strain is much higher than other models.

(A)

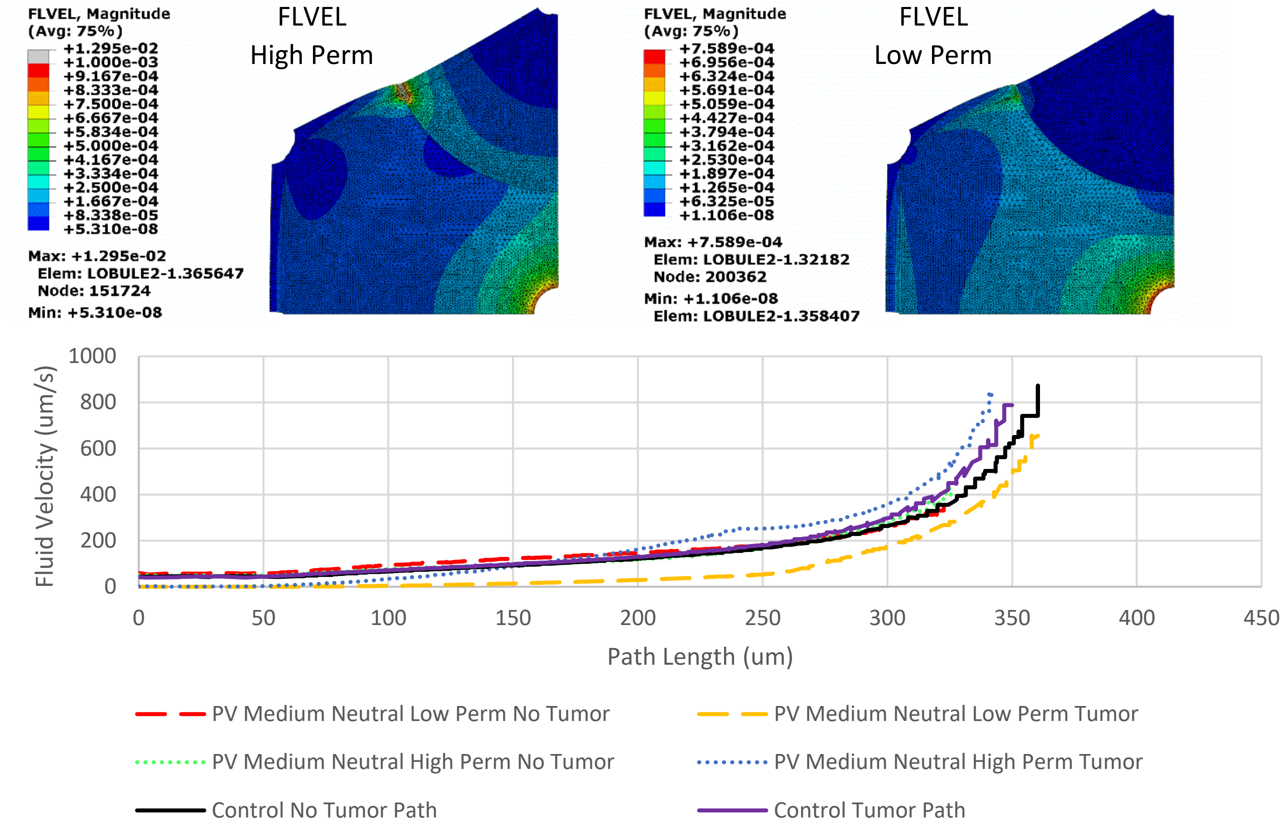
FLVEL, Magnitude
(Avg: 75%)
+1.295e-02
+1.000e-03
+9.167e-04
+8.333e-04
+7.500e-04
+6.667e-04
+5.834e-04
+5.000e-04
+4.167e-04
+3.334e-04
+2.500e-04
+1.667e-04
+8.338e-05
+5.310e-08
Max: +1.295e-02
Elem: LOBULE2-1.365647
Node: 151724
Min: +5.310e-08
FLVEL
High Perm
FLVEL, Magnitude
(Avg: 75%)
+7.589e-04
+6.956e-04
+6.324e-04
+5.691e-04
+5.059e-04
+4.427e-04
+3.794e-04
+3.162e-04
+2.530e-04
+1.897e-04
+1.265e-04
+6.325e-05
+1.106e-08
Max: +7.589e-04
Elem: LOBULE2-1.32182
Node: 200362
Min: +1.106e-08
Elem: LOBULE2-1.358407
FLVEL
Low Perm
Fluid Velocity (um/s)
1000
800
600
400
200
0
0
50
100
150
200
250
300
350
400
450
Path Length (um)
PV Medium Neutral Low Perm No Tumor
PV Medium Neutral Low Perm Tumor
PV Medium Neutral High Perm No Tumor
PV Medium Neutral High Perm Tumor
Control No Tumor Path
Control Tumor Path

(B)

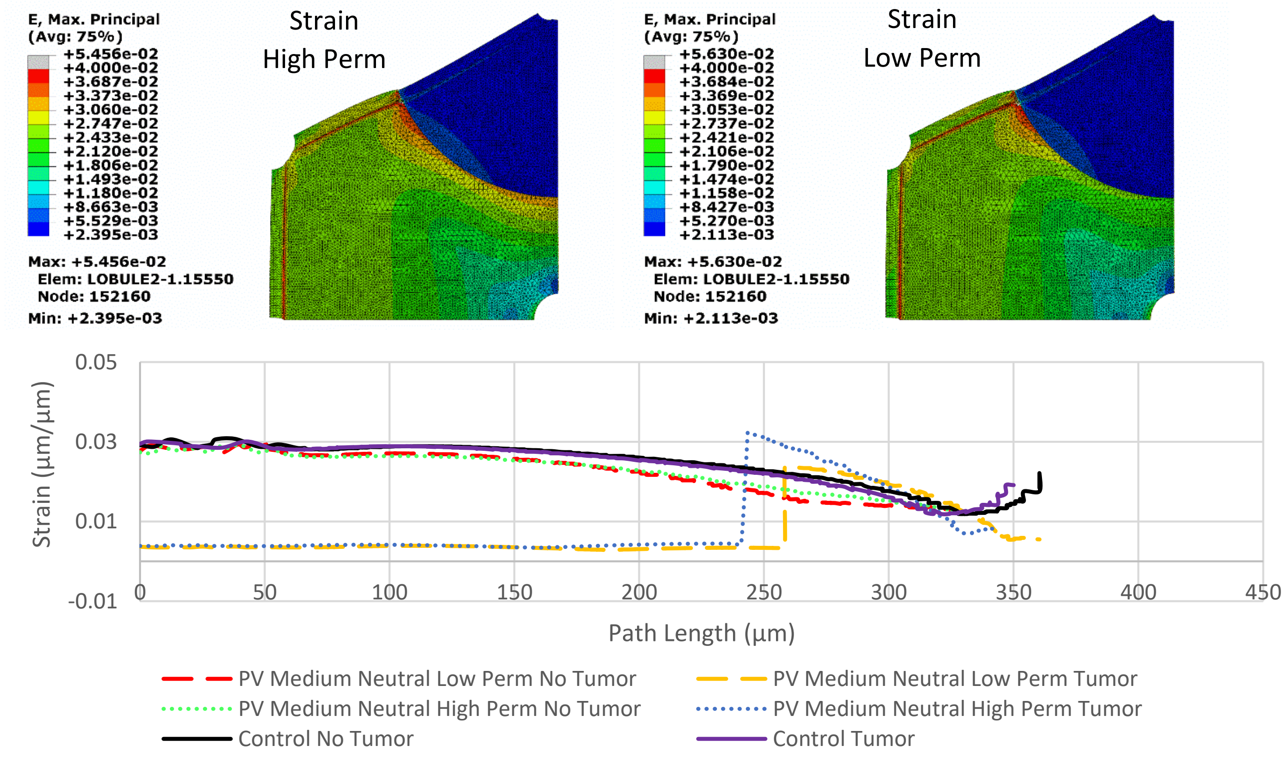
E, Max. Principal
(Avg: 75%)
+5.456e-02
+4.000e-02
+3.687e-02
+3.373e-02
+3.060e-02
+2.747e-02
+2.433e-02
+2.120e-02
+1.806e-02
+1.493e-02
+1.180e-02
+8.663e-03
+5.529e-03
+2.395e-03
Max: +5.456e-02
Elem: LOBULE2-1.15550
Node: 152160
Min: +2.395e-03
Strain
High Perm
E, Max. Principal
(Avg: 75%)
+5.630e-02
+4.000e-02
+3.684e-02
+3.369e-02
+3.053e-02
+2.737e-02
+2.421e-02
+2.106e-02
+1.790e-02
+1.474e-02
+1.158e-02
+8.427e-03
+5.270e-03
+2.113e-03
Max: +5.630e-02
Elem: LOBULE2-1.15550
Node: 152160
Min: +2.113e-03
Strain
Low Perm
Strain (µm/µm)
0.05
0.03
0.01
-0.01
0
50
100
150
200
250
300
350
400
450
Path Length (µm)
PV Medium Neutral Low Perm No Tumor
PV Medium Neutral Low Perm Tumor
PV Medium Neutral High Perm No Tumor
PV Medium Neutral High Perm Tumor
Control No Tumor
Control Tumor

C)

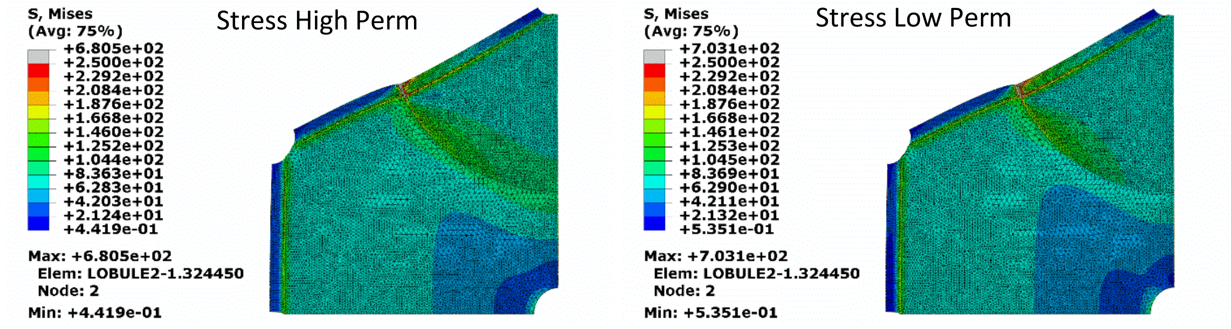


(D)

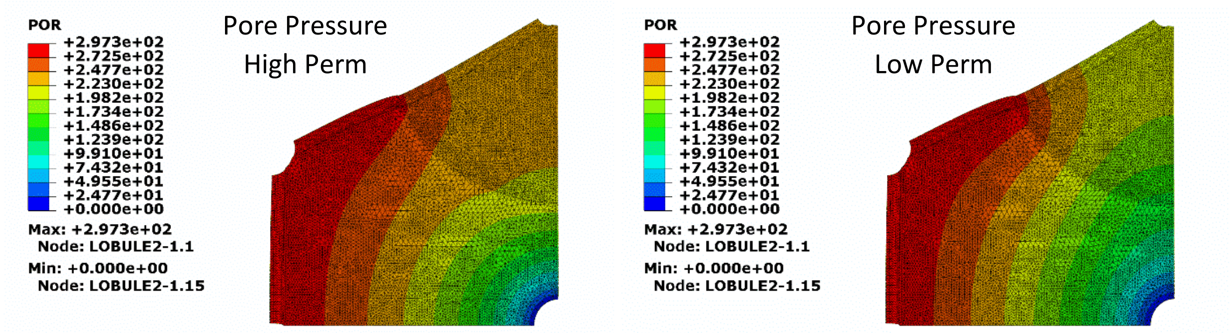


**Figure S12.** Images for the 400μm diameter tumor centered at the portal vein. Tumor was not set as a pressure source or sink. Images for (A) fluid velocity, (B) strain, (C) stress, and (D) pore pressure distribution throughout the quarter lobule. Graphs represent the fluid velocity and strain in the “tumor” and “no tumor” paths.

(A)

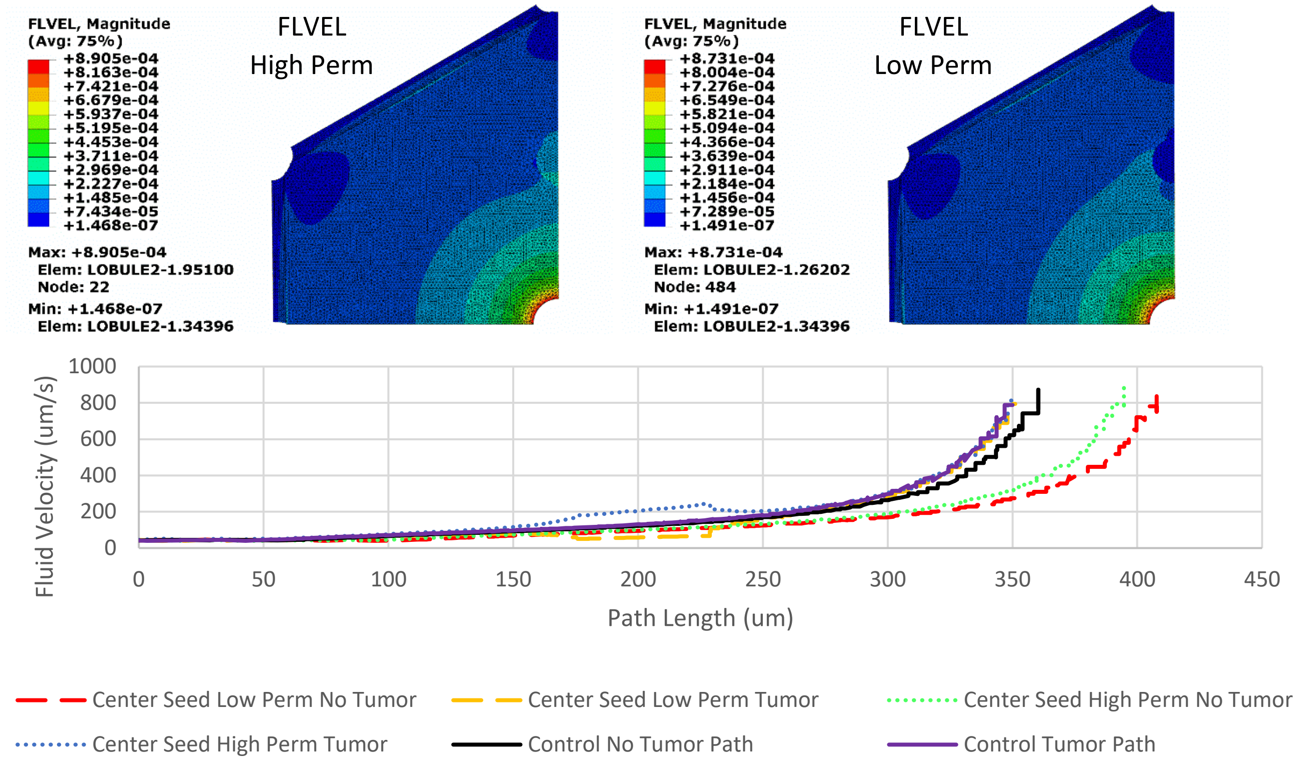
FLVEL, Magnitude
(Avg: 75%)
+8.905e-04
+8.163e-04
+7.421e-04
+6.679e-04
+5.937e-04
+5.195e-04
+4.453e-04
+3.711e-04
+2.969e-04
+2.227e-04
+1.485e-04
+7.434e-05
+1.468e-07
Max: +8.905e-04
Elem: LOBULE2-1.95100
Node: 22
Min: +1.468e-07
Elem: LOBULE2-1.34396
FLVEL
High Perm
FLVEL, Magnitude
(Avg: 75%)
+8.731e-04
+8.004e-04
+7.276e-04
+6.549e-04
+5.821e-04
+5.094e-04
+4.366e-04
+3.639e-04
+2.911e-04
+2.184e-04
+1.456e-04
+7.289e-05
+1.491e-07
Max: +8.731e-04
Elem: LOBULE2-1.26202
Node: 484
Min: +1.491e-07
Elem: LOBULE2-1.34396
FLVEL
Low Perm
Fluid Velocity (um/s)
1000
800
600
400
200
0
0
50
100
150
200
250
300
350
400
450
Path Length (um)
Center Seed Low Perm No Tumor
Center Seed Low Perm Tumor
Center Seed High Perm No Tumor
Center Seed High Perm Tumor
Control No Tumor Path
Control Tumor Path

(B)

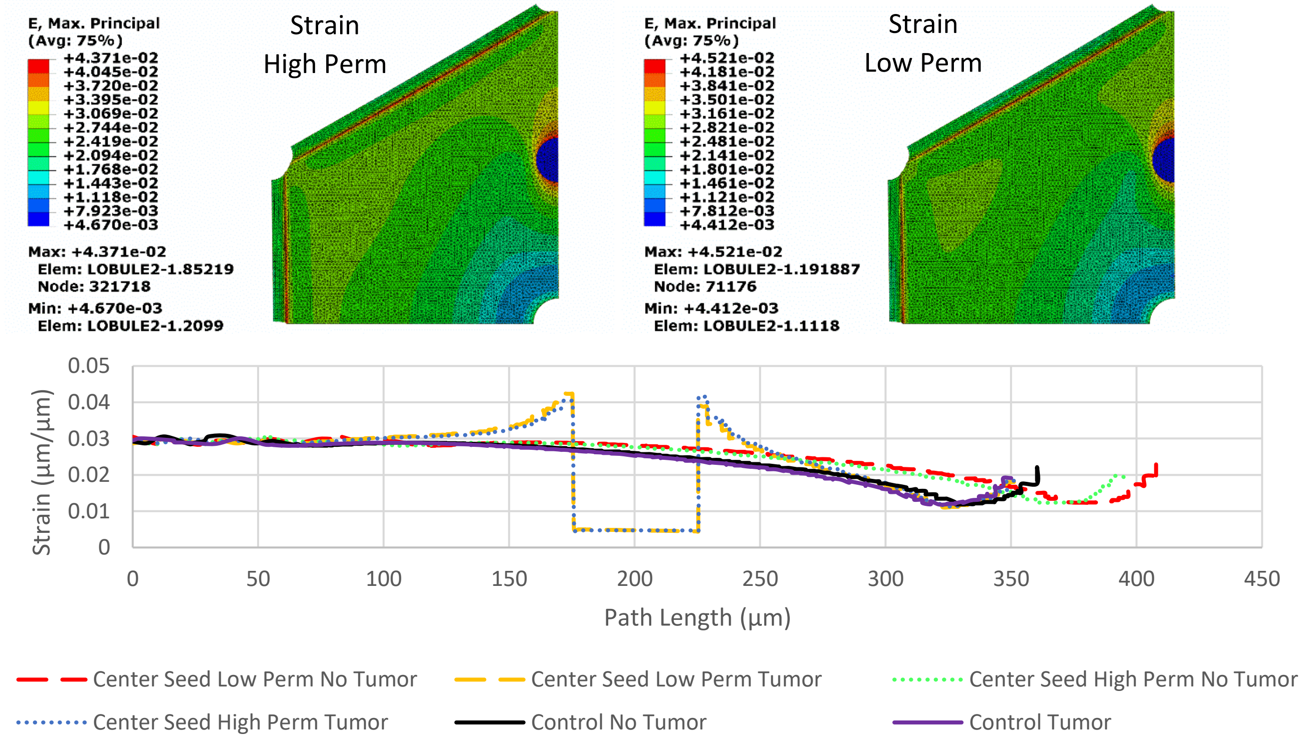
E, Max. Principal
(Avg: 75%)
+4.371e-02
+4.045e-02
+3.720e-02
+3.395e-02
+3.069e-02
+2.744e-02
+2.419e-02
+2.094e-02
+1.768e-02
+1.443e-02
+1.118e-02
+7.923e-03
+4.670e-03
Max: +4.371e-02
Elem: LOBULE2-1.85219
Node: 321718
Min: +4.670e-03
Elem: LOBULE2-1.2099
Strain
High Perm
E, Max. Principal
(Avg: 75%)
+4.521e-02
+4.181e-02
+3.841e-02
+3.501e-02
+3.161e-02
+2.821e-02
+2.481e-02
+2.141e-02
+1.801e-02
+1.461e-02
+1.121e-02
+7.812e-03
+4.412e-03
Max: +4.521e-02
Elem: LOBULE2-1.191887
Node: 71176
Min: +4.412e-03
Elem: LOBULE2-1.1118
Strain
Low Perm
Strain (µm/µm)
0.05
0.04
0.03
0.02
0.01
0
0
50
100
150
200
250
300
350
400
450
Path Length (µm)
Center Seed Low Perm No Tumor
Center Seed Low Perm Tumor
Center Seed High Perm No Tumor
Center Seed High Perm Tumor
Control No Tumor
Control Tumor

(C)

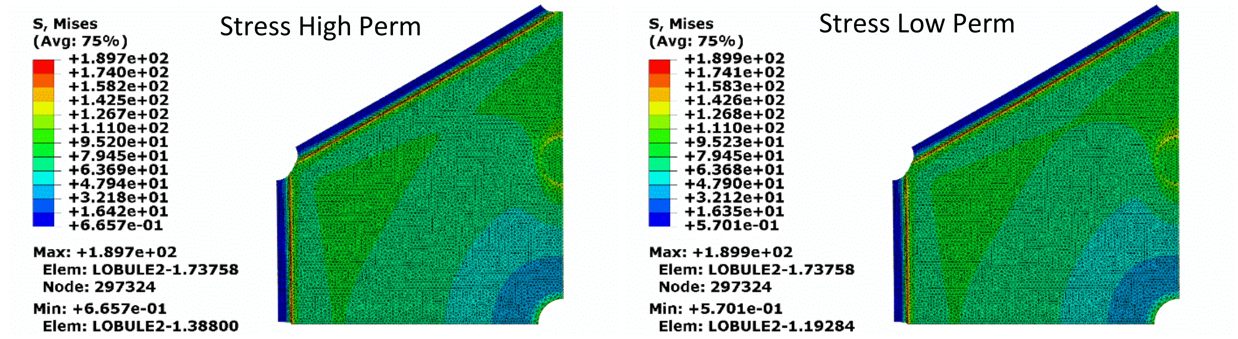


(D)

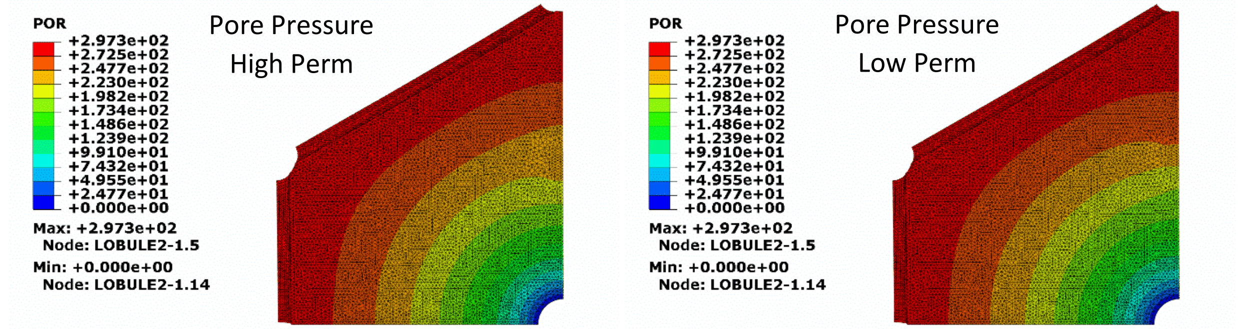


**Figure S13.** Images for the 50μm diameter tumor seed centered between the portal vein and central vein. Tumor was not set as a pressure source or sink. Images for (A) fluid velocity, (B) strain, (C) stress, and (D) pore pressure distribution throughout the quarter lobule. Graphs represent the fluid velocity and strain in the “tumor” and “no tumor” paths.

(A)

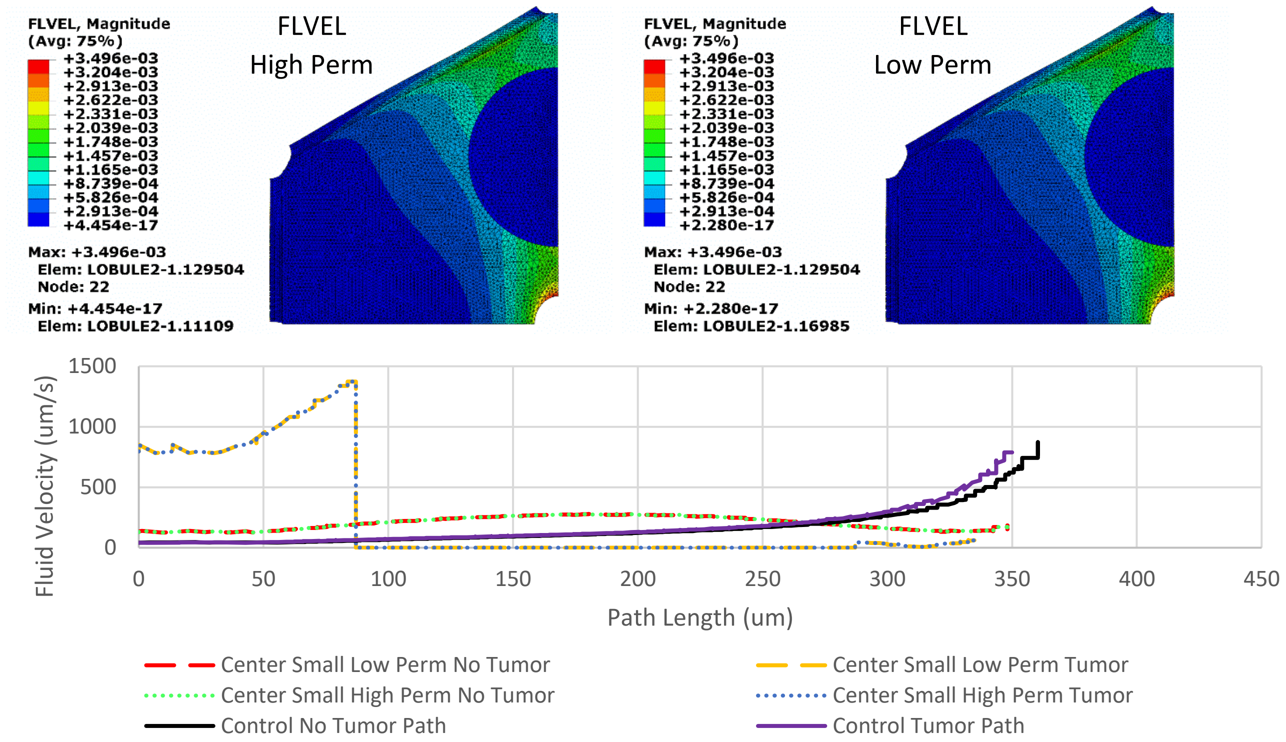
FLVEL, Magnitude
(Avg: 75%)
+3.496e-03
+3.204e-03
+2.913e-03
+2.622e-03
+2.331e-03
+2.039e-03
+1.748e-03
+1.457e-03
+1.165e-03
+8.739e-04
+5.826e-04
+2.913e-04
+4.454e-17
Max: +3.496e-03
Elem: LOBULE2-1.129504
Node: 22
Min: +4.454e-17
Elem: LOBULE2-1.11109
FLVEL
High Perm
FLVEL, Magnitude
(Avg: 75%)
+3.496e-03
+3.204e-03
+2.913e-03
+2.622e-03
+2.331e-03
+2.039e-03
+1.748e-03
+1.457e-03
+1.165e-03
+8.739e-04
+5.826e-04
+2.913e-04
+2.280e-17
Max: +3.496e-03
Elem: LOBULE2-1.129504
Node: 22
Min: +2.280e-17
Elem: LOBULE2-1.16985
FLVEL
Low Perm
Fluid Velocity (um/s)
1500
1000
500
0
0
50
100
150
200
250
300
350
400
450
Path Length (um)
Center Small Low Perm No Tumor
Center Small High Perm No Tumor
Control No Tumor Path
Center Small Low Perm Tumor
Center Small High Perm Tumor
Control Tumor Path

(B)

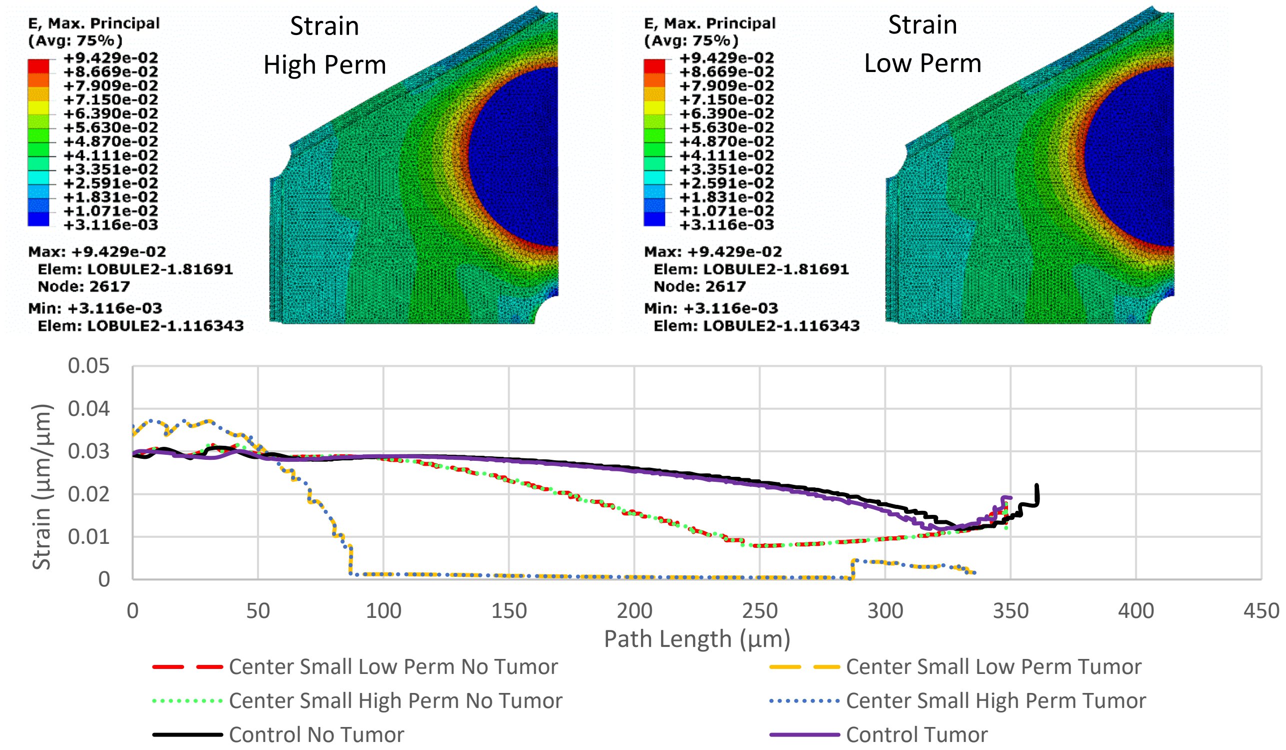
E, Max. Principal
(Avg: 75%)
+9.429e-02
+8.669e-02
+7.909e-02
+7.150e-02
+6.390e-02
+5.630e-02
+4.870e-02
+4.111e-02
+3.351e-02
+2.591e-02
+1.831e-02
+1.071e-02
+3.116e-03
Max: +9.429e-02
Elem: LOBULE2-1.81691
Node: 2617
Min: +3.116e-03
Elem: LOBULE2-1.116343
Strain
High Perm
E, Max. Principal
(Avg: 75%)
+9.429e-02
+8.669e-02
+7.909e-02
+7.150e-02
+6.390e-02
+5.630e-02
+4.870e-02
+4.111e-02
+3.351e-02
+2.591e-02
+1.831e-02
+1.071e-02
+3.116e-03
Max: +9.429e-02
Elem: LOBULE2-1.81691
Node: 2617
Min: +3.116e-03
Elem: LOBULE2-1.116343
Strain
Low Perm
Strain (µm/µm)
0.05
0.04
0.03
0.02
0.01
0
0
50
100
150
200
250
300
350
400
450
Path Length (µm)
Center Small Low Perm No Tumor
Center Small High Perm No Tumor
Control No Tumor
Center Small Low Perm Tumor
Center Small High Perm Tumor
Control Tumor

(C)

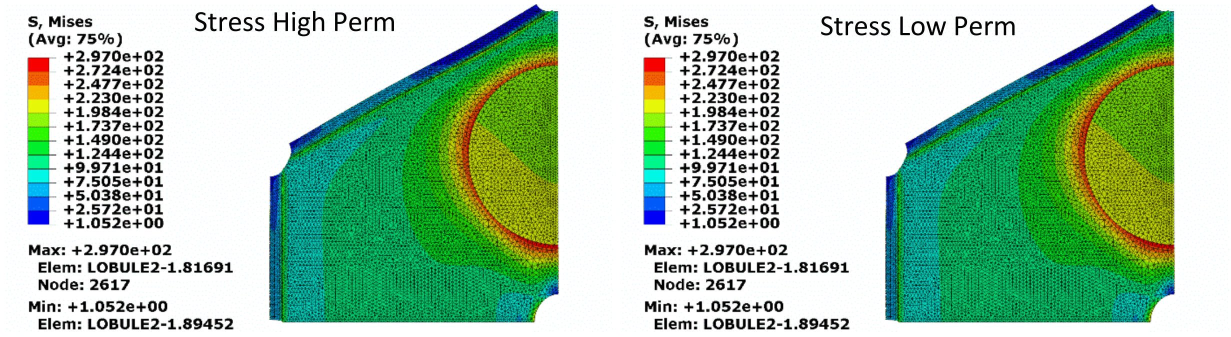


(D)

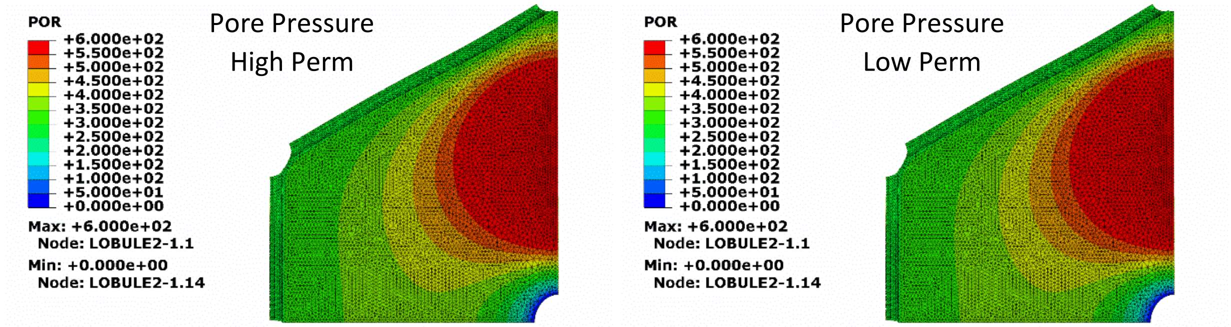


**Figure S14**. Images for the 200μm diameter tumor seed centered between the portal vein and central vein. Tumor was set as a pressure sink. Images for (A) fluid velocity, (B) strain, (C) stress, and (D) pore pressure distribution throughout the quarter lobule. Graphs represent the fluid velocity and strain in the “tumor” and “no tumor” paths. Note the scale of fluid velocity is higher than other models.

(A)

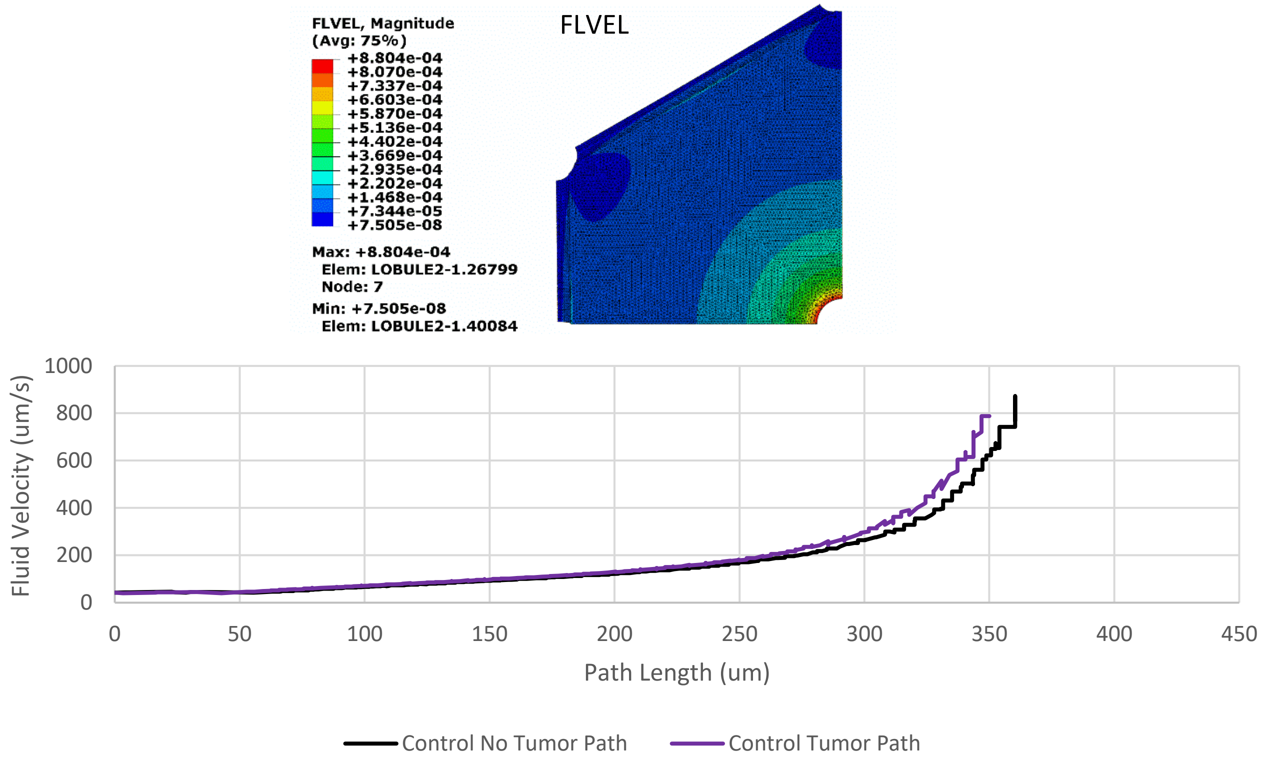
FLVEL, Magnitude
(Avg: 75%)
+8.804e-04
+8.070e-04
+7.337e-04
+6.603e-04
+5.870e-04
+5.136e-04
+4.402e-04
+3.669e-04
+2.935e-04
+2.202e-04
+1.468e-04
+7.344e-05
+7.505e-08
Max: +8.804e-04
Elem: LOBULE2-1.26799
Node: 7
Min: +7.505e-08
Elem: LOBULE2-1.40084
FLVEL
Fluid Velocity (um/s)
1000
800
600
400
200
0
0
50
100
150
200
250
300
350
400
450
Path Length (um)
Control No Tumor Path
Control Tumor Path

(B)

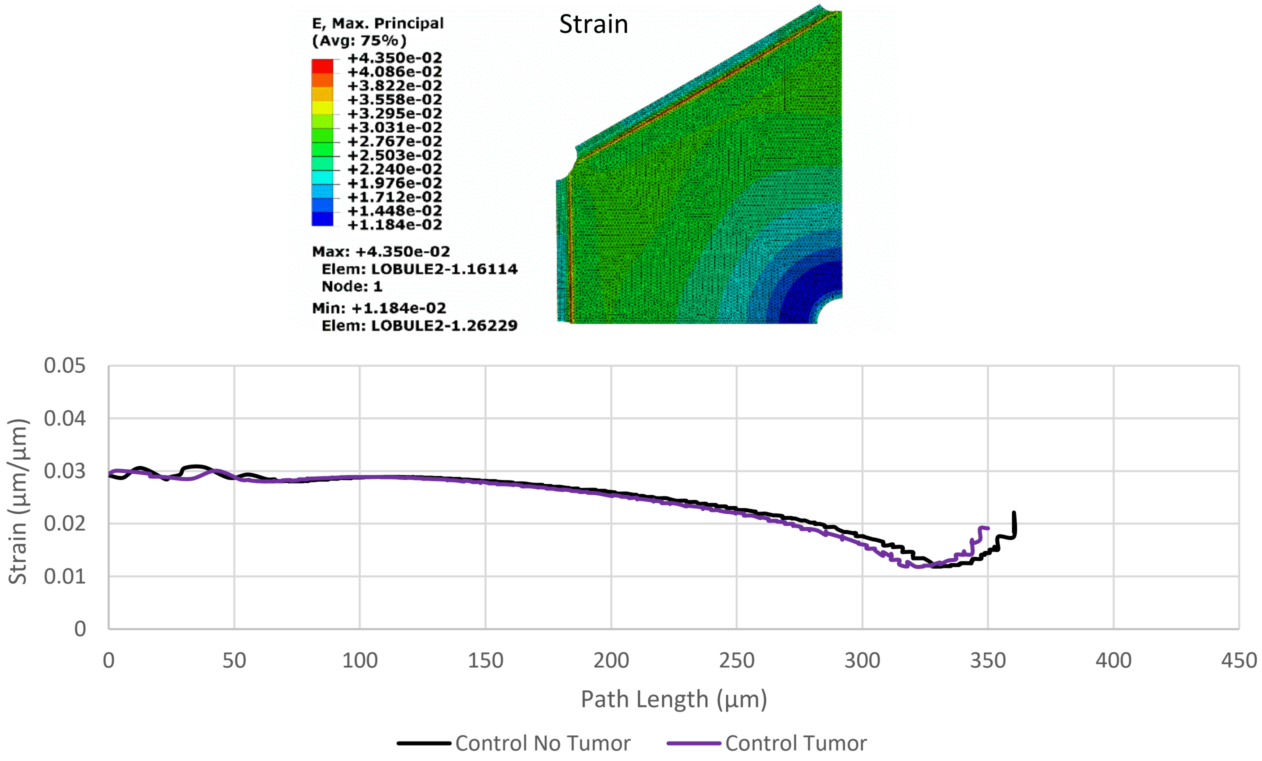
E, Max. Principal
(Avg: 75%)
+4.350e-02
+4.086e-02
+3.822e-02
+3.558e-02
+3.295e-02
+3.031e-02
+2.767e-02
+2.503e-02
+2.240e-02
+1.976e-02
+1.712e-02
+1.448e-02
+1.184e-02
Max: +4.350e-02
Elem: LOBULE2-1.16114
Node: 1
Min: +1.184e-02
Elem: LOBULE2-1.26229
Strain
Strain (µm/µm)
0.05
0.04
0.03
0.02
0.01
0
0
50
100
150
200
250
300
350
400
450
Path Length (µm)
Control No Tumor
Control Tumor

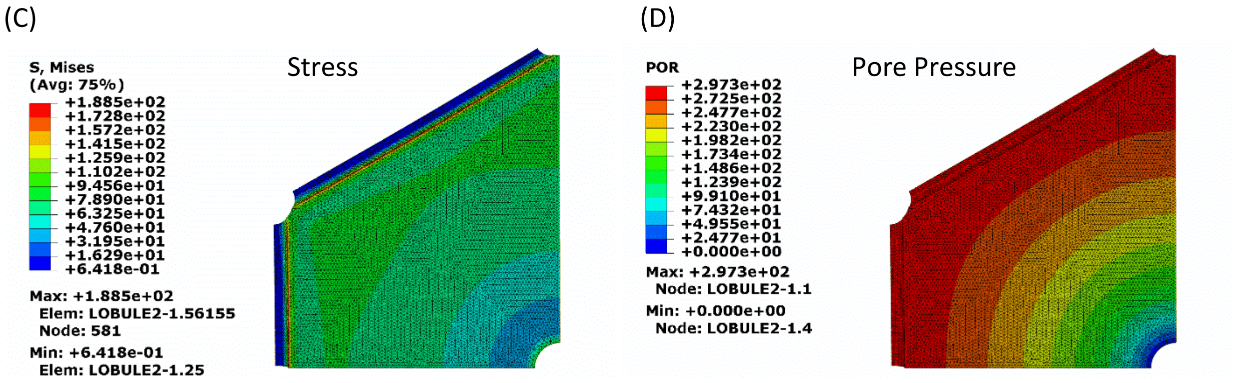


**Figure S15.** Images for the control lobule (there is no tumor anywhere in this model). Images for (A) fluid velocity, (B) strain, (C) stress, and (D) pore pressure distribution throughout the quarter lobule. Graphs represent the fluid velocity and strain in the “tumor” and “no tumor” paths (to keep path notation consistent with plots above). Data for control fluid velocity and strain can be found in other model graphs, but is also displayed here for clarity.

#### Supplemental Materials Bibliography


Bonfiglio A, Leungchavaphongse K, Repetto R, Siggers JH (2010) Mathematical Modeling of the Circulation in the Liver Lobule. J Biomech Eng 132:111011–111011. doi: 10.1115/1.4002563

Evans DW, Moran EC, Baptista PM, et al (2013) Scale-dependent mechanical properties of native and decellularized liver tissue. Biomech Model Mechanobiol 12:569–580. doi: 10.1007/s10237-012-0426-3

Lu Q, Ling W, Lu C, et al (2015) Hepatocellular Carcinoma: Stiffness Value and Ratio to Discriminate Malignant from Benign Focal Liver Lesions. Radiology 275:880–888. doi: 10.1148/radiol.14131164

Netti PA, Berk DA, Swartz MA, et al (2000) Role of Extracellular Matrix Assembly in Interstitial Transport in Solid Tumors. Cancer Res 60:2497–2503.

Nishii K, Reese G, Moran EC, Sparks JL (2016) Multiscale computational model of fluid flow and matrix deformation in decellularized liver. J Mech Behav Biomed Mater 57:201–214. doi: 10.1016/j.jmbbm.2015.11.033

Pishko GL, Astary GW, Mareci TH, Sarntinoranont M (2011) Sensitivity Analysis of an Image-Based Solid Tumor Computational Model with Heterogeneous Vasculature and Porosity. Ann Biomed Eng 39:2360–2373. doi: 10.1007/s10439-011-0349-7

Swabb EA, Wei J, Gullino PM (1974) Diffusion and Convection in Normal and Neoplastic Tissues. Cancer Res 34:2814–2822.

Venkatesh SK, Yin M, Glockner JF, et al (2008) Magnetic Resonance Elastography of Liver Tumors- Preliminary Results. AJR Am J Roentgenol 190:1534–1540. doi: 10.2214/AJR.07.3123
